# Designing viral diagnostics with model-based optimization

**DOI:** 10.1101/2020.11.28.401877

**Authors:** Hayden C. Metsky, Nicole L. Welch, Priya P. Pillai, Nicholas J. Haradhvala, Laurie Rumker, Sreekar Mantena, Yibin B. Zhang, David K. Yang, Cheri M. Ackerman, Juliane Weller, Paul C. Blainey, Cameron Myhrvold, Michael Mitzenmacher, Pardis C. Sabeti

## Abstract

Diagnostics, particularly for rapidly evolving viruses, stand to benefit from a principled, measurement-driven design that harnesses machine learning and vast genomic data—yet the capability for such design has not been previously built. Here, we develop and extensively validate an approach to designing viral diagnostics that applies a learned model within a combinatorial optimization framework. Concentrating on CRISPR-based diagnostics, we screen a library of 19,209 diagnostic–target pairs and train a deep neural network that predicts, from RNA sequence alone, diagnostic signal better than contemporary techniques. Our model then makes it possible to design assays that are maximally sensitive over the spectrum of a virus’s genomic variation. We introduce ADAPT (https://adapt.guide), a system for fully-automated design, and use ADAPT to design optimal diagnostics for the 1,933 vertebrate-infecting viral species within 2 hours for most species and 24 hours for all but 3. We experimentally show ADAPT’s designs are sensitive and specific down to the lineage level, including against viruses that pose challenges involving genomic variation and specificity. ADAPT’s designs exhibit significantly higher fluorescence and permit lower limits of detection, across a virus’s entire variation, than the outputs of standard design techniques. Our model-based optimization strategy has applications broadly to viral nucleic acid diagnostics and other sequence-based technologies, and, paired with clinical validation, could enable a critically-needed, proactive resource of assays for surveilling and responding to pathogens.

## Introduction

Diagnostics and routine surveillance underpin infectious disease countermeasures, and recent developments in nucleic acid detection have enhanced these efforts^1–8^. Yet there has been limited progress enhancing diagnostics and surveillance through innovations in computational design. This is surprising in light of new machine learning and optimization methods, and the explosion of viral genomic data. Indeed, designing viral assays from genomic data is still done largely by hand, without well-defined objectives and with a great deal of trial and error.

Harnessing predictive models, together with an optimization algorithm, promises many benefits for diagnostics and routine surveillance. It could design assays that are more sensitive than existing ones at low viral concentrations and across viral variation. It would enable a large resource of assays, developed in advance of outbreaks, that are broadly effective against known viruses and even many novel ones. And it would allow for rapid design of new assays and efficient redesign to reflect new viral variants over time. Here, we demonstrate these capabilities by developing and experimentally validating a design approach that combines a deep learning model with combinatorial optimization across vast viral genomic data.

We provide significant advances in three areas: (i) modeling and predicting the efficiency of a viral diagnostic; (ii) integrating a virus’s variation optimally into a diagnostic’s design; and (iii) designing assays rapidly at scale using a fully-automated system.

The first challenge we address is predicting a diagnostic’s efficiency in detecting a nucleic acid target. The paradigm for the most advanced diagnostic design methods, which focus on qPCR^9–15^, is to make binary decisions about efficiency—an assay will detect a viral genome or will not—according to thermodynamic criteria and simple heuristics. These heuristics include constraints on the number or positions of assay-target mismatches. Yet binary decisions are rudimentary—their accuracy is unclear and efficiency runs on a continuous scale—so they still impel experimental comparisons of assays and likely miss the optimum. Quantitative predictions of detection efficiency would enable a more sensitive diagnostic. In contrast to the latest techniques for diagnostics, our approach uses extensive experimental data and a machine-learned model to predict efficiency in detecting a viral target. Concentrating on CRISPR-based diagnostics, we screen 19,209 guide-target pairs, forming the largest dataset on diagnostic efficacy to our knowledge; our dataset exposes the effect of viral variation on a diagnostic by emphasizing imperfect guide-target homology. We then train a deep neural network that predicts, from nucleotide sequence alone, the enzymatic activity in a detection reaction, which corresponds to a diagnostic’s sensitivity.

Predictive models have been previously built for various CRISPR systems^16–18^. These models predict on-target cleavage by measuring knockdown efficacy or indel frequency, and several^18, 19^ have focused on predictions for CRISPR-Cas13 enzymes using manually-defined features; Cas13 enzymes have applications in diagnostics and are our modeling focus in this work. Subsequent studies^20,21^ have applied one of these models^18^ to improve Cas13d guide design for viral RNA knockdown, having use in antivirals. To our knowledge, our modeling is the first that applies deep learning to predict Cas13 guide activity. And the collateral cleavage activity exhibited by certain CRISPR enzymes, such as Cas13a, underlies nucleic acid detection yet that activity is more challenging to measure in high-throughput than RNA knockdown. Our dataset and modeling is also the first, to our knowledge, applied to collateral cleavage activity and designed principally to evaluate viral diagnostic performance. While we concentrate on CRISPR-based viral diagnostics, our same predictive approach can be applied to other nucleic acid detection technologies, such as qPCR, and to non-viral targets.

The second challenge we confront is viral variation. Influenza A virus (FLUAV) RT-qPCR tests often have false-negative rates over 10%^22–24^ (nearly 100% on some circulating strains) owing to extensive sequence variation, and the issue besets diagnostics and surveillance for many other viruses^25–29^. State-of-the-art design methods that account for variation generally follow one of two paradigms. One paradigm^10, 12, 13^ is to identify the most conserved genomic regions and then design an assay targeting such regions, usually matching a single genome sequence: this is inadequate because conserved viral regions are rarely free of variation, and targeting them may not give the optimal sensitivity and antagonizes specificity between related viruses. The other paradigm^9, 11, 14, 15^ is to minimize the complexity of an assay (number of primer/probes) subject to a constraint that a sufficient amount of known variation is detected (or “covered”): a downside is that, by handling variation through a constraint, this approach does not expressly optimize for sensitivity. Our predictive model allows us to directly integrate variation into the objective function and expressly optimize sensitivity over it—a formulation that differs from and advances on current techniques. Specifically, we design assays that maximize predicted detection efficiency in expectation over a virus’s genomic variation.

The third challenge we tackle involves scalability. The number of viral genomes is growing exponentially^30, 31^, encompassing well-known viruses and novel strains that may cause epidemics (Supplementary Fig. 1). This growth complicates the design of diagnostic and surveillance assays, which is based on those genomes. Moreover, viral evolution can necessitate periodic updating as variants emerge. FLUAV subtyping assays will lose sensitivity over time, especially profoundly following a pandemic (Fig. 1a and Supplementary Fig. 2). And in the case of COVID-19, as in other outbreaks, mutations quickly accumulated on top of the early genome sequences (Fig. 1b) and some have created failures in widely-used SARS-CoV-2 diagnostics^32, 33^, compelling redesigns that accommodate variation. Yet the current paradigm for designing an assay requires curating viral genome data, which in practice is laborious and difficult to keep pace with viral evolution. To overcome this obstacle, we also introduce ADAPT (**A**ctivity-informed **D**esign with **A**ll-inclusive **P**atrolling of **T**argets), a system that implements our approach and automatically integrates the latest viral genomes from public databases into the design process. It is, to our knowledge, the first fully-automated system for viral diagnostic design, and it operates at a massive scale, rapidly designing assays that integrate all genomes on NCBI for all vertebrate-infecting viruses.

**Figure 1.**
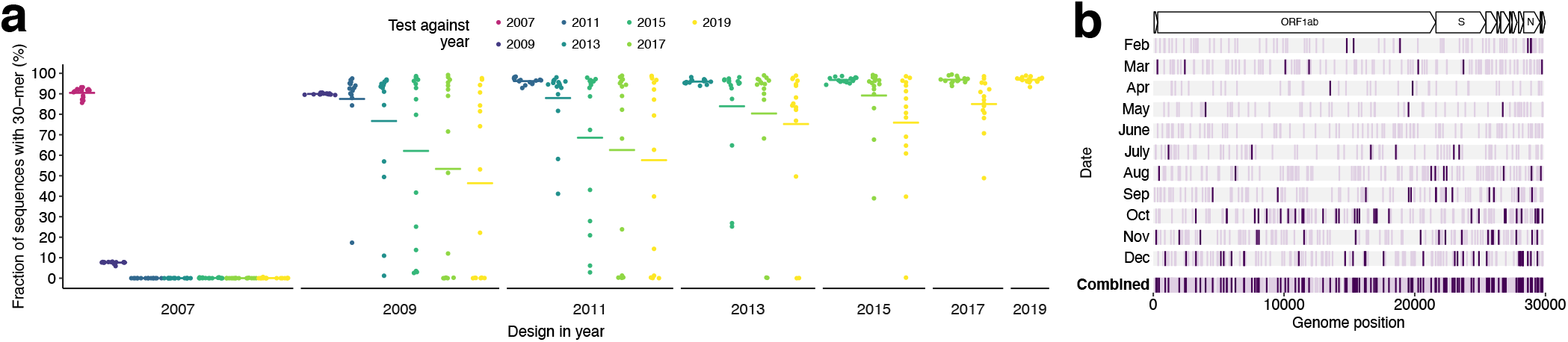
Emerging viral variation over time and the effect on diagnostic performance. **(a)** Diagnostic performance for influenza A virus subtyping may degrade over time, even considering conserved sites. At each year, we select the 15 most conserved 30-mers from recent sequences for segment 6 (N) for all N1 subtypes; each point is a 30-mer. Plotted value is the fraction of sequences in subsequent years (colored) that contain the 30-mer; bars are the mean. To aid visualization, only odd years are shown. 2007 N1 30-mers are absent following 2007 owing to antigenic shift during the 2009 H1N1 pandemic. **(b)** Variation along the SARS-CoV-2 genome emerging over time. Bottom row (‘Combined’) shows all 1,131 single nucleotide polymorphisms, among 361,460 genomes, that crossed 0.1% or 1% frequency between February 1 and the end of 2020—i.e., (i) at <0.1% frequency in genomes collected before February 1 and 0.1–1% frequency by December 31 (light purple); or (ii) at <1% frequency before February 1 and at ≥1% frequency by December 31 (dark). Other rows show the month in which each polymorphism crosses the frequency threshold.

We applied our model-based optimization approach to design maximally sensitive, species-specific diagnostics for the 1,933 viral species known to infect vertebrates. We experimentally show their potential on SARS-CoV-2 and other viruses. Using synthetic targets designed to represent known variation, we validate our approach against target variation more extensively than is typical for diagnostics: we evaluate 69 designs, each against up to 15 targets that encompass one virus’s variation (290 different targets in total). The results demonstrate that our approach provides designs with comprehensive detection across genomic variation and lineage-level specificity. Our designs also outperform the outputs of standard design techniques that are based on conventional CRISPR diagnostic design heuristics and sequence conservation.

## Results

### Predicting efficiency of a CRISPR-based diagnostic

We first consider the problem of predicting the efficiency of a diagnostic in detecting a viral target. We sought to advance beyond the current rule-based assay design paradigm—in which methods use thermodynamic criteria and simple heuristics^9–15^—by building a machine learning model that predicts efficiency. To accomplish that, we constructed a dataset of fluorescence over time, which corresponds to diagnostic sensitivity, using CARMEN^8^. CARMEN is a multiplexed droplet-based platform that performs a massive number of detection reactions in parallel; here, we ran the detection reactions to assess a library of assay designs. We then developed a model of efficiency informed by the reaction kinetics, which we applied to our fluorescence measurements, allowing us to train our tailored machine learning model to accurately predict efficiency from nucleotide sequence alone. Such an approach is more measurement-driven than contemporary assay design techniques and offers quantitative evaluations of a design’s efficacy, including against viral variants.

We focus on CRISPR-Cas13 technologies^1, 2^, in which Cas13 enzymes use guide RNAs to detect a target sequence and then exhibit collateral RNase activity that cleaves fluorescent reporters, leading to a fluorescent diagnostic readout. Collateral activity is a key property distinguishing Cas13 and other CRISPR enzymes that enable nucleic acid detection from the more widely-studied uses involving on-target cleavage^16–18^, which can be measured by sequencing. Prior studies have characterized sequence requirements for the RNA reporters^2, 34^ and have established Cas13 guide design principles—such as the importance of a protospacer flanking site (PFS), RNA secondary structure, and a mismatch-sensitive “seed” region^35–37^—but have not measured the collateral activity of guide-targets pairs in high-throughput nor modeled it.

To enable a model, we first designed a library of 19,209 unique LwaCas13a guide-target pairs (Fig. 2a and Supplementary Fig. 3a). The library has a sequence composition representative of viral genomes (Methods), an average of 2.9 mismatches between each guide and target (Supplementary Fig. 3b), and a representation of the different PFS alleles (Supplementary Fig. 3c). The reporter presence during each pair’s reaction falls over time following an exponential decay, and thus we use the negative of the decay to model fluorescence over time (Fig. 2a; Methods). The fluorescence growth rate is a function of the target concentration and enzymatic efficiency of a guide-target-Cas13a complex, so we hold the former constant to evaluate the latter (Supplementary Fig. 4a,b); we define activity as the logarithm of this growth rate, serving as our measurement of enzymatic efficiency.

**Figure 2.**
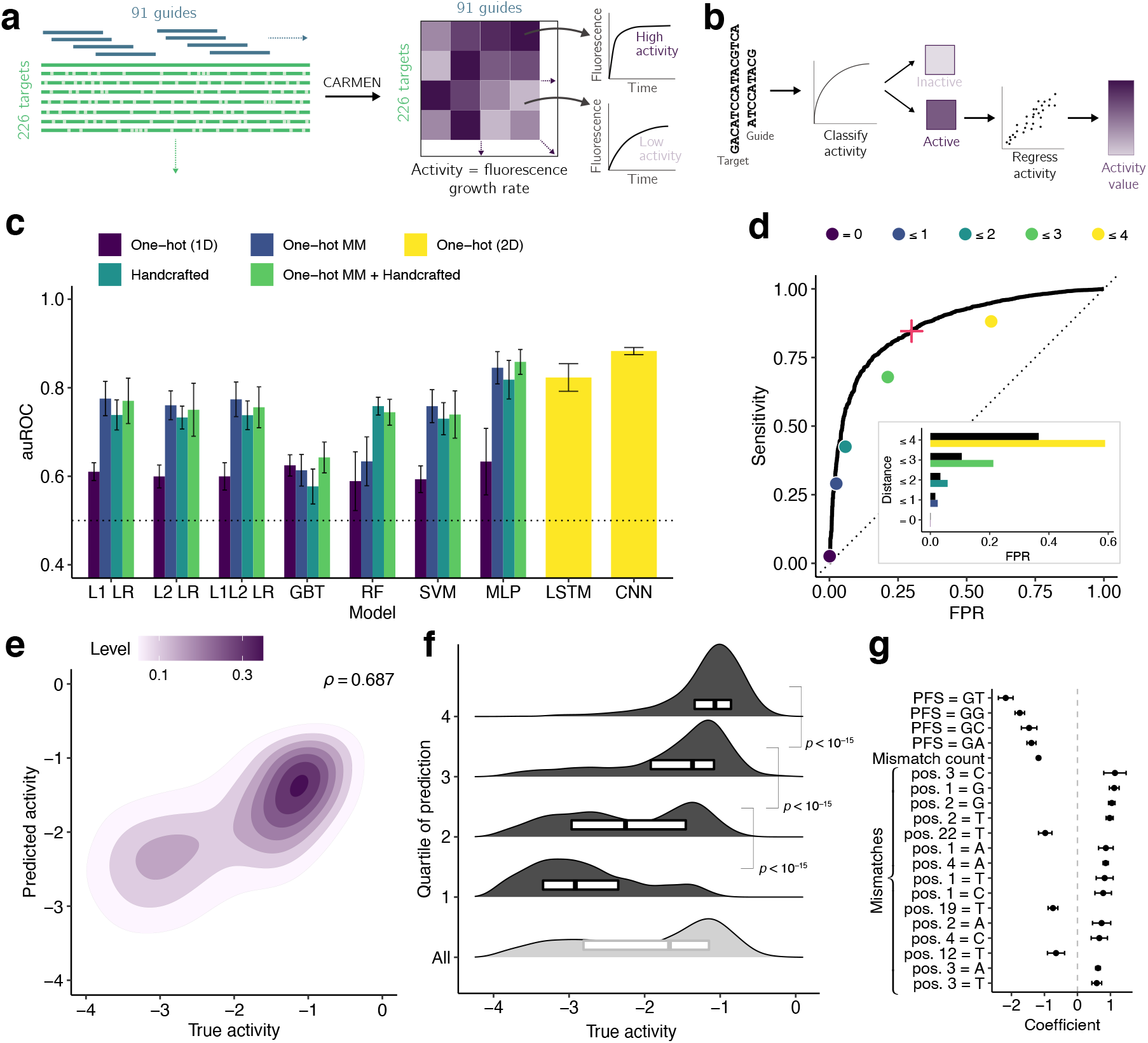
Predicting CRISPR-Cas13a activity. **(a)** The library consists of a 865 nt long wildtype target sequence and 91 guide RNAs complementary to it, along with 225 unique targets containing mismatches and varying PFS alleles relative to the wildtype (19,209 unique guide-target pairs). We measure fluorescence every ~20 minutes for each pair and use the growth rate to quantify activity. **(b)** We predict activity for a guide-target pair in two parts: training a classifier on all pairs and a regression model on the active pairs. **(c)** Results of model selection, for classification, with nested cross-validation. For each model and input type (color) on five outer folds, we performed a five-fold cross-validated hyperparameter search. Plotted value is the mean auROC of the five models, and error bar is the 95% confidence interval. L1 LR and L2 LR, logistic regression; L1L2 LR, elastic net; GBT, gradient-boosted classification tree; RF, random forest; SVM, support vector machine; MLP, multilayer perceptron; LSTM, long short-term memory recurrent neural network; CNN, neural network with parallel convolutional filters and a locally-connected layer. One-hot (1D), one-hot encoding of target and guide sequence independently, i.e., without encoding a pairing of nucleotides between the two; One-hot MM, one-hot encoding of target sequence nucleotides and of mismatches in guides relative to the target; Handcrafted, curated features of hypothesized importance (details in Methods); One-hot (2D), one-hot encoding of target and guide sequence with encoded guide-target pairing. Supplementary Fig. 6 shows auPR and regression models. **(d)** ROC curve, on a held-out test set, of a CNN classifying pairs as inactive or active. Points indicate sensitivity and false positive rate (FPR) for baseline heuristic classifiers: choosing a guide-target pair to be active if it has a non-G PFS and the guide-target Hamming distance is less than a specified threshold (color). Inset plot shows comparison of FPR between CNN (black) and baseline classifiers at equivalent sensitivity. Red ‘+’ indicates decision threshold in ADAPT. **(e)** Regression results, on the held-out test set, of a CNN predicting activity values of active guide-target pairs. Contour color, point density. *ρ*, Spearman correlation. Supplementary Fig. 11 shows regression using full dataset. **(f)** Same data as (e). Each row contains one quartile of pairs divided by predicted activity (top row is predicted most active), with the gray row showing all active pairs. Smoothed density estimates and interquartile ranges show the distribution of true activity for the pairs in each quartile. *P*-values are from Mann-Whitney *U* tests (one-sided). **(g)** Top 20 feature coefficients, by absolute value, in L1 logistic regression model classifying activity with ‘One-hot MM + Handcrafted’ features. Dot is the mean over training on five folds and error bar is the 95% confidence interval. Mismatch features indicate a mismatch with the base being the complement of the spacer nucleotide; positions are relative to the target (pos. 28 is 5’ end of spacer).

We measured the fluorescence arising from the library’s guide-target pairs roughly every 20 minutes with CARMEN, and from these measurements we calculated the activity for each pair. The data include, on average, about 15 replicate activity measurements per pair (Supplementary Fig. 5a–d) and exhibit activity differences, spanning several orders of magnitude in fluorescence growth rate, between different guide sequences and across the different target variants each guide detects (Supplementary Fig. 5e–g). Our measurements are consistent with expected covariates of activity, including the PFS and number of mismatches.

Using our dataset, we developed a machine-learned model to predict CRISPR-Cas13a activity of a guide-target pair. We use a two-step hurdle model: classifying a pair as inactive or active, and then regressing activity for active pairs (Fig. 2b and Supplementary Fig. 4b; 86.8% of the full dataset is labeled active). For classifying guide-target pairs as inactive or active, we performed nested crossvalidation—fitting models multiple independent times, on separate splits of the training data—to evaluate our fitting procedure and compare nine models potentially suited to this task using different inputs, including one-hot encodings (representing nucleotide sequences directly with binary vectors) and handcrafted features. We found that a deep convolutional neural network (CNN), trained from nucleotide sequence alone, outperforms the other models (Fig. 2c and Supplementary Fig. 6a). For regression on active pairs, we also found that a CNN outperforms other models, albeit with a less noticeable improvement over the simpler models than for classification (Supplementary Fig. 6b,c). CNNs have performed well when regressing Cas12a guide RNA on-target editing activity on matching target sequences^17^, likely because the convolutional layers detect motifs in the sequences; here, such layers may also detect patterns of mismatches. In all model training, we accounted for measurement error (Methods).

We proceeded with further evaluating CNN models to use for designing CRISPR-based diagnostics. During model selection, our space of CNN models allows for multiple parallel convolutional and locally connected filters of different widths (Supplementary Fig. 7). The latter use separate (“local”) filters for different regions of the guide-target sequence. In addition to convolutional layers, which are common in genomics applications, our model search also prefers to incorporate locally connected layers (Supplementary Fig. 8 and 9). We hypothesize they help the model uncover fixed spatial dependencies in this data, such as a mismatch-sensitive seed region, that would be missed by convolutional layers and potentially difficult for fully connected layers to learn.

We evaluated our models’ performance on a held-out test set of CRISPR-Cas13a guide-target pairs (Methods) and against methods that use standard Cas13a design rules. Our classification CNN performs well on the held-out data (auROC = 0.866; auPR = 0.972 with 85.6% of the test set being true positive; Fig. 2d and Supplementary Fig. 10a). When the guide and target are not identical, it yields a lower false-positive rate and higher precision than a heuristic classifying activity according to the PFS and guide-target divergence (Fig. 2d and Supplementary Fig. 10b). Our regression CNN also predicts the activity of active guide-target pairs well (Spearman’s *ρ* = 0.687; Fig. 2e) and accurately separates pairs into quartiles based on predicted activity (Fig. 2f). A regression on the complete dataset would also predict the activity of guide-target pairs well (Spearman’s *ρ* = 0.774; Supplementary Fig. 11), but is less suited to the problem than our two-step hurdle model.

Further exploring our CNN models’ performance, we considered that some critical features, such as the PFS and number of mismatches, could explain much of the models’ performance. Yet the models retain predictive ability when evaluated individually on different choices of PFS (Supplementary Fig. 12a,b and 13a,b) and mismatch thresholds (Supplementary Fig. 12c,d and 13c,d), albeit often with reduced performance compared to the full dataset. For example, among only the perfectly matching guide-target pairs in the test set, we find auROC = 0.869 for classifying activity and Spearman’s *ρ* = 0.472 for regression on the active pairs. A learning curve shows that additional data similar to our current dataset would not improve model performance (Supplementary Fig. 14).

Precision matters greatly in our application because we would like confidence that designs determined to be active are indeed active. To use these models as part of our design process, we chose a decision threshold for the classifier that yields a precision of 0.975 (Methods; Fig. 2d and Supplementary Fig. 10a). We apply a two-step hurdle model (Fig. 2b) by defining a piecewise activity function that is 0 if a guide-target pair is classified as inactive, and the regressed activity if active.

We tested our model’s performance on datasets from two independent studies, starting with ref. 37. Measurements from this study use a different ortholog (LbuCas13a) than the one used for training our model (LwaCas13a): we mitigated the effects of LbuCas13a’s greater overall cleavage activity^34^ and adjusted for different guide lengths (Methods). Following these changes (Supplementary Fig. 15a), we found that our model’s predictions of LwaCas13a activity correlate well with the independently-measured LbuCas13a cleavage rates (Spearman’s *ρ* = 0.816 with *ρ* < 10^-4^; Supplementary Fig. 15b), providing validation of our model on an independent dataset. Considering data on LbuCas13a—RNA binding affinity from this same study, we found that pairs with high predicted collateral activity rarely exhibit low binding affinity, whereas pairs with high binding affinity can have low predicted collateral activity (Supplementary Fig. 15c–f). This relationship is consistent with binding being necessary, but not sufficient, to achieve collateral activity.

We also tested our model on data from ref. 36, which assesses on-target (*cis*-) cleavage by measuring RNA knockdown using LwaCas13a. Our model’s predictions correlate highly with Lwa-Cas13a knockdown levels (Spearman’s *ρ* = −0.826 with *ρ* < 10^-10^; Supplementary Fig. 15g). The roughly linear relationship between on-target and predicted collateral cleavage activity (Pearson’s *r* = −0.868 with *ρ* < 10^-12^) suggests that our model could predict Cas13a on-target cleavage and that high-throughput on-target assays could be valuable for modeling Cas13a collateral activity. These results provide another independent validation of our model’s performance and show its generalizability.

Beyond modeling CRISPR diagnostic activity, we examined our dataset to understand LwaCas13a preferences, starting with the PFS. Prior studies have seen weaker detection activity with a `G` PFS for LwaCas13a^1^ and characterized the PFS preference, if any, for other orthologs^35,37^ and in other settings^36^. We also observe reduced activity with a `G` PFS for LwaCas13a and, extending to another position, we find that `GA`, `GC`, and `GG` provide higher activity than `GT` (Supplementary Fig. 16a), suggesting 3’ `HV` may be a more stringent preference. Mechanistically, the reduced activity of a `G` PFS results from base pairing with the crRNA’s complementary `C` in the direct repeat sequence immediately 5’ of the spacer, which prevents separation of the guide-target duplex^38^; the direct repeat’s subsequent base is `A`, so the additional complementarity of `GT` may explain this further reduction.

The breadth of our data also illuminates effects of mismatches (Supplementary Fig. 16b) on Lwa-Cas13a activity. U-g mismatches (`U` in target, `G` in guide RNA spacer) rescue activity in our dataset, though we do not observe this same effect for G-u mismatches (Supplementary Fig. 16c); while both wobble pairings might be tolerated for binding, the asymmetry could stem from how the pairings affect nuclease activation. The data also clarifies, for LwaCas13a, the mismatch-sensitive seed region that had previously been identified, using a limited number of guide-target pairs, for Lbu-Cas13a^37^ and LshCas13a^35^. The worst-performing guide-target pairs are relatively likely to contain mismatches in positions 6–11 of the guide RNA spacer (Supplementary Fig. 16d), concordant with the known region. We also find nearly full tolerance for multiple mismatches on the 3’ end of the spacer (5’ end of the target binding site; Supplementary Fig. 16e), a property that our predictive model learns and leverages when positioning CRISPR guides in variable regions. Coefficients from the regularized linear models included in our model comparison are consistent with these findings on LwaCas13a preferences. In particular, the coefficients signal that the PFS and number of mismatches dominate activity (Supplementary Fig. 17a,b), that there exist position-specific nucleotide preferences along the target (Supplementary Fig. 17c), and that the effect of mismatched nucleotides varies along the guide-target complex (Supplementary Fig. 17d). The greater magnitude of coefficients on mismatches compared to those on nucleotide composition suggests that, in general, the prescence of a mismatch at a position will more considerably affect activity than the choice of matching nucleotide (Supplementary Fig. 17c,d).

While our dataset and model focus on the CRISPR-Cas13a detection technology, a similar measurement-driven approach could be applied to and benefit other nucleic acid technologies for viral diagnostics; the remainder of our work is agnostic to the model.

### Designing maximally efficient assays across variation

An advantage of our model-based approach is that it provides quantitative predictions that can then be used within an optimization framework to obtain highly effective viral diagnostic assays. With a model in hand to accurately predict the detection efficiency of a guide-target pair, we sought an algorithm to design assays that are maximally efficient in detecting a virus’s variation. This formulation more explicitly optimizes sensitivity than the often-used design approaches^9–15^ of targeting conserved regions or treating the detection of sequence variation as a constraint.

We aim to formulate the problem of designing optimal diagnostic probe sequences over known variation. The result is broadly applicable to many detection technologies given a learned model. We rely on all known sequences within a genomic region, represented by *S*, and an activity function that quantifies detection efficiency between a probe and a targeted sequence. In the case of Cas13a-based diagnostics, probe sequences are CRISPR guide RNAs and the activity function is our predictive model of guide-target detection activity; later, we address how to identify optimal regions, including amplification primers. We initially construct a ground set of possible probes by finding representative subsequences in *S*, through rapidly clustering subsequences using locality-sensitive hashing^39^. Our goal is to find the set *P* of probes, a subset of the ground set, that maximizes the expected activity between *P* and *S* (Fig. 3a). In this objective function, the expectation is over the sequences in *S* and the probability for each sequence reflects a prior on applying the probes toward that sequence (in practice, we usually use a uniform prior over the sequences). Larger numbers of probes would require more detection reactions or, if they are multiplexed in a single reaction, may interfere^40^, with the resulting kinetic impact harming sensitivity^41^. For these reasons, we impose a penalty in our objective function and a hard constraint on the number of probes (details in Supplementary Note 1a).

**Figure 3.**
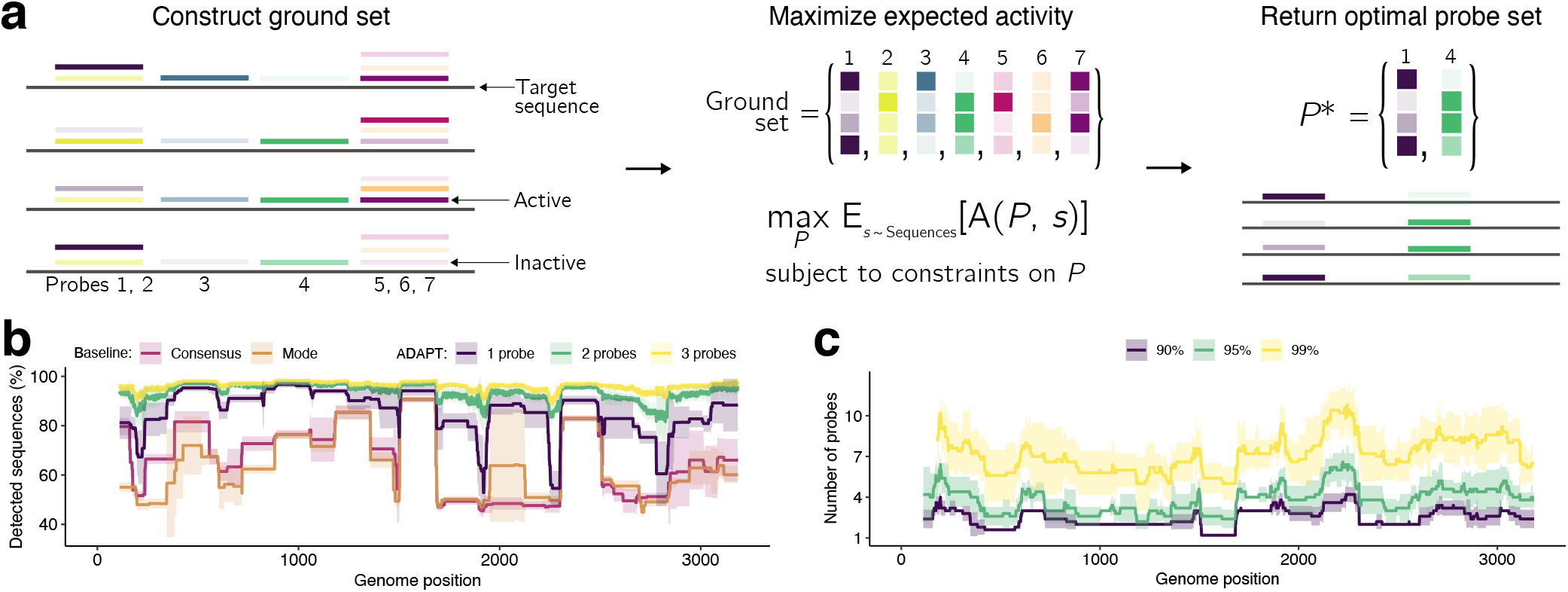
Maximizing detection efficiency across genomic variation. **(a)** Our approach for designing maximally active probe sets. We (1) find representative subsequences in a genomic region, which are possible probes (colored); (2) compute an activity (shaded) between each probe and each target sequence *s*, forming the ground set of probes; (3) find a probe set *P*, a subset of the ground set, maximizing the expected activity A(P, s) between P and s, subject to soft and hard constraints on P (including on |P|, the number of probes). **(b)** Fraction of Lassa virus (LASV; segment S) genomes detected, with different design strategies in a 200 nt sliding window, using a model in which 30 nt probes detect a target if they are within 1 mismatch, counting G-U pairs at matches. ‘Consensus’, probe-length consensus subsequence that detects the most number of genomes in the window; ‘Mode’, most common probe-length subsequence within the window. Our approach (ADAPT) uses hard constraints of 1–3 probes and maximizes activity, **(c)** Number of probes when solving a dual objective: minimizing the number of probes to detect >90%, >95%, and >99% of LASV genomes using the model in (b). In (b) and (c), lines show the mean and shaded regions around them are 95% pointwise confidence bands across genomes sampled for LASV calculated by bootstrapping, i.e., randomly sampling genomes to be input to the design process.

Having formulated an objective function, we then developed an approach to maximize it using combinatorial optimization. Our objective function is non-monotone submodular and, under activity constraints on which probes are allowed in the ground set, is non-negative. That the function is nonmonotone means that while initially its value will increase as the number of probes increases, the value may eventually decrease; this is an expected property because, while adding more probes may initially improve detection, our function penalizes the number of them. Submodularity corresponds to the property of “diminishing returns”—that is, as the number of probes increases, the improvement in performance from each additional probe diminishes, which is also an expected property. We apply a fast randomized combinatorial algorithm^42^ for maximizing a non-negative and nonmonotone submodular function under a hard constraint on the number of probes, which provides a probe set whose objective value is within a factor 1*/e* of the optimal. Supplementary Note 1a contains the proofs and algorithmic details. We also evaluated the canonical greedy algorithm^43^ for constrained submodular maximization; it returns similar results in practice (Supplementary Fig. 18), though does not offer provable guarantees in our case because it requires monotonicity whereas our function is non-monotone.

We benchmarked our approach’s comprehensiveness across sequence variation. To make this benchmarking interpretable, here we chose a detection activity function that equals 1 if a probe is within 1 mismatch of a target (detected) and 0 otherwise (not detected); this function has the property that expected activity is equivalent to the fraction of genomes detected. Two simple but often-used strategies for constructing probes—using the consensus or most common sequence in a region—fail to capture diversity for Lassa virus and other highly diverse viruses (Fig. 3b and Supplementary Fig. 19a).

Our approach, to maximize expected activity, yields designs with greater comprehensiveness than the two simple strategies on these same viruses. The designs from our approach detect more variation—even when constrained to a single probe—across the genome, and the amount detected increases as we permit more probes (Fig. 3b and Supplementary Fig. 19a). And if we compare against more than one of the most common subsequences—generalizing the baseline strategy that selects the single most common, to allow it to use multiple probes—our approach at an equivalent number of probes still detects more variation across the genomes of diverse viruses (Supplementary Fig. 19b). This result is expected because our approach explicitly maximizes detection over the variable sequences. Using a related objective function that minimizes the number of probes subject to constraints on comprehensiveness (Supplementary Note 1b), we find it is possible to reach near-complete comprehensiveness across the genome with few probes (Fig. 3c and Supplementary Fig. 19c). On species with less extensive diversity, such as Zika virus, the simple strategies perform well (Supplementary Fig. 19a,b), suggesting that our more involved approach is not always necessary. Nevertheless, having options to comprehensively target many regions of a genome enables us to integrate genuine activity predictions and enforce viral taxon-specificity, both of which will constrain the designs.

In many diagnostic and routine surveillance applications, assays must distinguish between viral species or strains that are genetically similar. In patient diagnostics, closely-related viruses can cause clinically similar symptoms, and a highly specific assay—whether run in a singleplex or multiplex panel—helps to identify the underlying infection or rule out possibilities. And with routine surveillance uses that test for a large number of viruses, taxon-specific assays can better pinpoint sample contents. When solving our combinatorial optimization problem, we enforce taxon-specificity by constraining the ground set of probes to only ones deemed specific. To enable that constraint, we wish to query probes in the ground set against an index of off-target taxa (Supplementary Note 2a). Determining whether a probe is specific to a viral taxon faces two computational challenges: it should tolerate multiple mismatches between a probe and potential off-targets and, when the probes and targets are RNA (as with Cas13-based diagnostics), account for G-U wobble base pairs. Accommodating the latter is critical: we found that ignoring G-U pairing when designing viral species-specific probes can result in missing nearly all off-target hits and deciding many probes to be specific to a viral species when they likely are not (Supplementary Fig. 20). Similar challenges arise in other RNA applications, such as the design of small interfering RNA, but most prior approaches simply ignore the G-U problem and, if they do address it, they use a solution inadequate for viral diagnostic goals (details in Supplementary Note 2b).

We experimented with two algorithms, which address these challenges, to decide a probe’s specificity to a viral taxon. Since near-neighbor searches can efficiently locate similar nucleotide sequences, we first tested a probabilistic near-neighbor query algorithm (i.e., one that may miss hits) with a tunable reporting probability for identifying off-target hits (Supplementary Note 2c). When amplified across many designs, the technique misses off-target hits and thus permits non-specific guides; a sufficiently high reporting probability would make this outcome unlikely, but we found it makes queries too slow to be practical in our case. Thus, we instead developed a data structure and query algorithm—more tailored to our problem—that are fully sensitive to high divergence and G-U wobble base pairing. The data structure indexes *k*-mers from all input taxa, in which the *k*-mers are split across many small tries according to a hash function (Supplementary Fig. 21 and Supplementary Note 2d). The approach is exact—it provably identifies all off-target hits—and therefore, in theory, guarantees high specificity of our designs. When designing species-specific viral diagnostics, the approach runs 10–100 times faster than a simple data structure with the same capability (Supplementary Fig. 22).

### Designing comprehensive diagnostics at scale

To accommodate design in the face of extensive and ever-growing viral genomic data, we built ADAPT. ADAPT fully-automatically designs and outputs the best assay options using our machine learning-based optimization approach, which includes transparently interfacing with viral genomic databases so that designs always reflect the latest available data (Fig. 4a).

**Figure 4.**
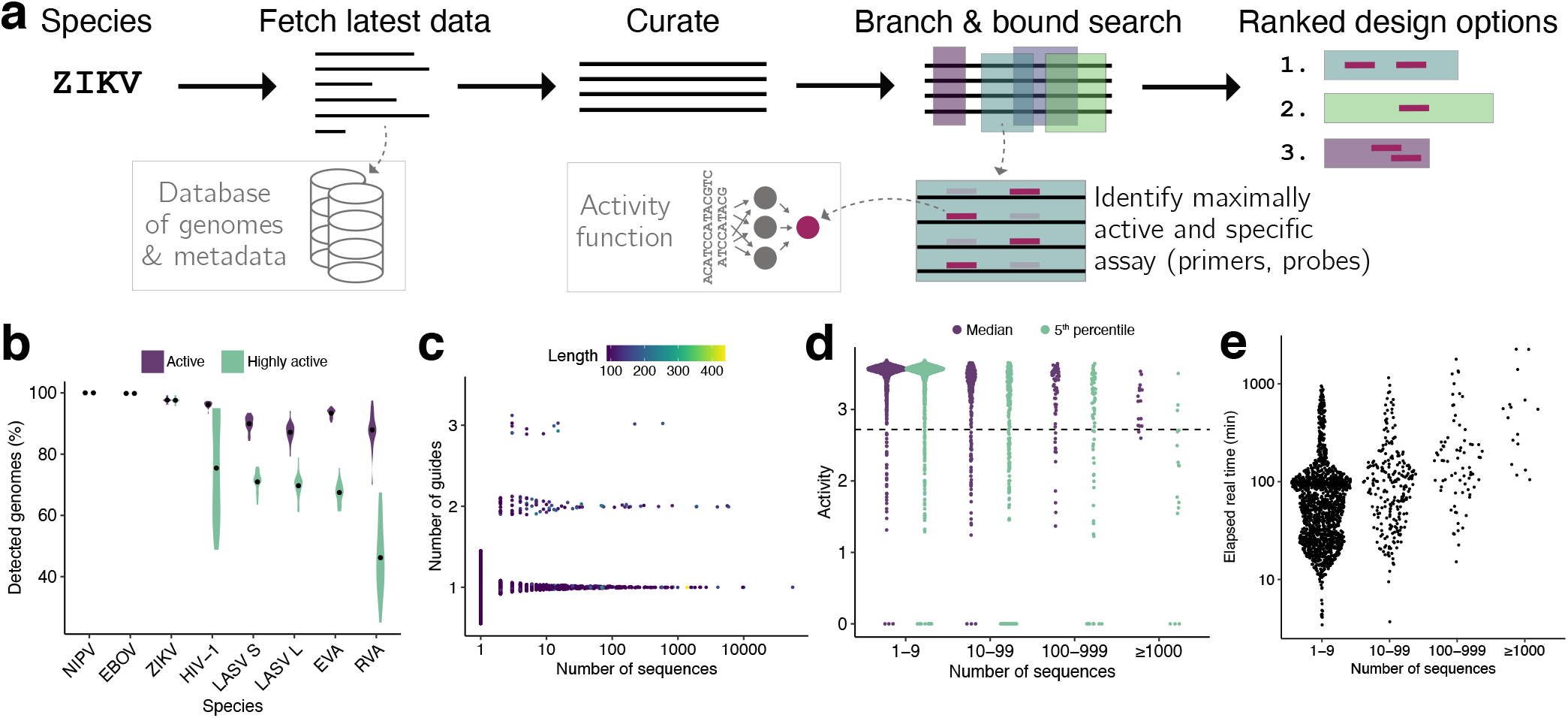
End-to-end design with ADAPT. **(a)** Sketch of ADAPT’s steps. ADAPT accepts taxonomy identifiers and fetches and curates their sequences from NCBI’s viral genome databases. It performs a branch and bound search to find genomic regions—each is an amplicon with primers—that contain a maximally active and taxon-specific probe set. ADAPT outputs the top design options, ranked by an objective function. **(b)** Cross-validated evaluation of detection. For each species, we ran ADAPT on 80% of available genomes and estimated performance, averaged over the top 5 design options, on the remaining 20%. Distributions are across 20 random splits of these genomes and dots indicate mean. Purple, fraction of genomes detected by primers and for which Cas13a guides are classified as active. Green, same except Cas13a guides also have an activity, predicted by the regression model, in the top 25% of our dataset (deemed “highly active”). NIPV, Nipah virus; EBOV, Zaire ebolavirus; ZIKV, Zika virus; LASV S/L, Lassa virus segment S/L; EVA, Enterovirus A; RVA, Rhinovirus A. **(c)** Number of Cas13a guides in the top-ranked design option for each species in the vertebrate-infecting virus designs. Color is the length of the targeted region, i.e., the amplicon, in nt. **(d)** Activity of each guide set with two summary statistics of its performance across known genome sequences: median and the 5^th^-percentile taken across each species’s sequences. For the latter, value *a* indicates that 95% of sequences are detected with activity ≥ a. Dashed line indicates the “high activity” threshold from (b). Sequences at 0 activity are predicted to not be detected by the classifier. Activities shown here are shifted up by 4 compared to the model output from Fig. 2. **(e)** End-to-end elapsed real time running ADAPT. In (c–e), each point is a vertebrate-infecting viral species.

ADAPT searches across a viral genome to identify regions to target for detection. These regions include amplification primers designed to achieve a desired coverage of sequence diversity, and ADAPT scores the regions according to their amplification potential and the activity of an optimal probe set within the region. ADAPT’s genome-wide search follows the branch and bound paradigm and identifies the best *N* diverse design options; users preset *N*, for which smaller values speed the search. Diverse design options enable assays that target multiple distinct regions of a genome. ADAPT memoizes repeated computations during its search, which decreases its runtime by 99.71– 99.96% for species we tested (Supplementary Fig. 23). Details of primer design are in Supplementary Note 3a and details of the search algorithm are in Supplementary Note 3b. ADAPT downloads and curates all near-complete or complete genomes from NCBI databases^31^ for a specified virus taxonomy, as well as for building the index that enforces specificity (Supplementary Note 3c). Fully-automated, scalable design greatly simplifies the design process and helps assays keep pace with evolution, especially for viruses with a large and growing number of public genome sequences. To accommodate additional applications, ADAPT supports many customizations in its predictive modeling and optimization components.

For some viruses there are few available genome sequences, especially early in an outbreak, and therefore little informative data on how it varies. We developed a scheme that uses the GTR nucleotide substitution model^44^ to forecast relatively likely combinations of genome substitutions in the region a probe detects, allowing us to estimate a probability that a probe’s activity will degrade over time (Supplementary Note 4 and Supplementary Fig. 24a). Applied to SARS-CoV-2, we found that, for some Cas13a designs, there is a low probability (~10%) their predicted activity will degrade over a 5 year period (Supplementary Fig. 24b,c). The decay results from mutations at mismatch-sensitive sites or at sites in the context of the Cas13a protospacers, which would impact some designs more than others. This forecasting may be helpful in risk-averse situations, but it has only a minor effect overall on ADAPT’s design option rankings (Supplementary Fig. 24d).

We computationally evaluated designs output by ADAPT, accounting together for their primers and probes, on seven RNA viruses spanning a range of genomic diversity. We used our CRISPR-Cas13a activity function, so the probes are Cas13a guides. Algorithmic randomness and the particular distribution of known sequences may introduce variability; we found that ADAPT’s designs are rarely exactly the same across runs (Supplementary Fig. 25a,b), but that they do often target similar genomic regions (Supplementary Fig. 25c,d). Cross-validation confirms the designs’ generalization: designs are predicted to be active in detecting >85% of held-out genomes for all seven species and have, in all but one species, “high activity” (defined as top 25% of measurements in our dataset) in detecting the majority of held-out genomes (Fig. 4b). Relaxed design choices, which permit more complex assays (Methods), can achieve a higher sensitivity, with designs predicted to be active in detecting >96% of held-out genomes for all seven species and >98% for all but one (Supplementary Fig. 26). The results show that ADAPT’s outputs are robust across different viruses.

We then applied ADAPT to design species-specific detection assays, including amplification primers and Cas13a guides, for the 1,933 viral species known to infect vertebrates. The designs have short amplicons (generally <200 nt) and use 3 or fewer guides for all species, including ones with thousands of genome sequences (Fig. 4c and Supplementary Fig. 27a). Thus, the assays are practical. For 95% of species, the guides detect the majority of known genomes with high predicted activity (Fig. 4d; for 88% of species, they detect >95% with high activity). If we were to instead optimize a reformulation of our maximization objective—minimize the number of guides subject to detecting >98% of genomes with high activity—we would also obtain few guides for most species but 40 species would require more than 3 guides (Supplementary Fig. 27b) and, in one extreme case, as many as 73 (Enterovirus B).

Along with detecting known viruses, our assays—designed to comprehensively detect species-level diversity—could detect novel viruses that are nested within known species, as is commonly the case. We simulated the design of assays in 2018 for detecting the SARS-related coronavirus species and then evaluated their detection of SARS-CoV-2, which is a part of the species but did not emerge until a year later. ADAPT’s second-highest–ranked assay is predicted to detect SARS-CoV-2 well and two other designs in the top five are predicted to exhibit low activity for SARS-CoV-2 (Supplementary Fig. 28a,b). This result is facilitated by bat SARS-like viral genomes that were available in 2018 and had homology to SARS-CoV-2. Nevertheless, heavy sampling biases hinder the efficacy of ADAPT’s designs: in 2018, SARS-CoV-1 was overrepresented in the species (85%) relative to bat SARS-like viruses owing to SARS outbreak sequencing. After down-weighing ADAPT’s consideration to SARS-CoV-1 (Methods), four of ADAPT’s five highest-ranked 2018 assays are predicted to detect SARS-CoV-2 well (Supplementary Fig. 28c,d). Such broadly-effective assays promise to benefit diagnostics and routine surveillance by constituting a proactively developed toolkit.

We also examined the computational requirements of designing assays for 1,933 viral species. End-to-end design with ADAPT completed in under 2 hours for 80% of species, under 24 hours for all but 3 species (human cytomegalovirus, SARS-related coronavirus, and FLUAV), and under 38 hours for all; runtime depended in part on the number of genome sequences (Fig. 4e). ADAPT required about 1 to 100 GB of memory per species (Supplementary Fig. 27c), which may necessitate using cloud computing or similar services. After curating available sequences, ADAPT considers all or almost all genome sequences for most species (Supplementary Fig. 27d–f), including the ones with >1,000 sequences; it does retain fewer than half of sequences for 38 species, which might be appropriate but prompts further investigation. Enforcing species-specificity imposes a considerable computational burden, adding to ADAPT’s runtime and memory usage (Supplementary Fig. 29a,b) while decreasing the solution’s activity and objective value because of the constraints, as expected (Supplementary Fig. 29c,d); relaxing the stringency of these specificity constraints can greatly decrease the required computational resources, which could be helpful for some users. Overall, the fast runtime implies it is possible to rapidly produce new, optimal designs that reflect evolving genomic variation or to detect a novel virus.

### Experimental evaluation of ADAPT’s designs

We experimentally benchmarked the diagnostic utility of our approach. We first focused on the US CDC’s SARS-CoV-2 qPCR assay amplicons, a target both of qPCR and CRISPR-based diagnostic assays, which allows us to benchmark our approach against previously-used design strategies for CRISPR-based SARS-CoV-2 diagnostics^45, 46^. As baselines in the N1 amplicon, we selected a Cas13a guide at the site of the qPCR probe and 10 random guides in the amplicon, all having an active (non-G) PFS; selecting guides according to the PFS is the canonical design strategy, and the distribution of their activity in this amplicon is a benchmark that mirrors the previously-used design strategies. The guide designed by our approach exhibits higher fluorescence at low target concentrations than all 11 of the baseline guides (Fig. 5a). Its fluorescence also grows at a faster rate than all 11 (Fig. 5b and Supplementary Fig. 30a). We observed similar results when repeating the comparison in the N2 amplicon (Supplementary Fig. 30b,c). Background activity does not impact these comparisons because the background (no-template) fluorescence is generally similar across guides with different performances and low compared to fluorescence against the intended target (Supplementary Fig. 31). These findings indicate that our model-based design permits better sensitivity, against a known target sequence, than the canonical rule-based approach focused on the PFS.

**Figure 5.**
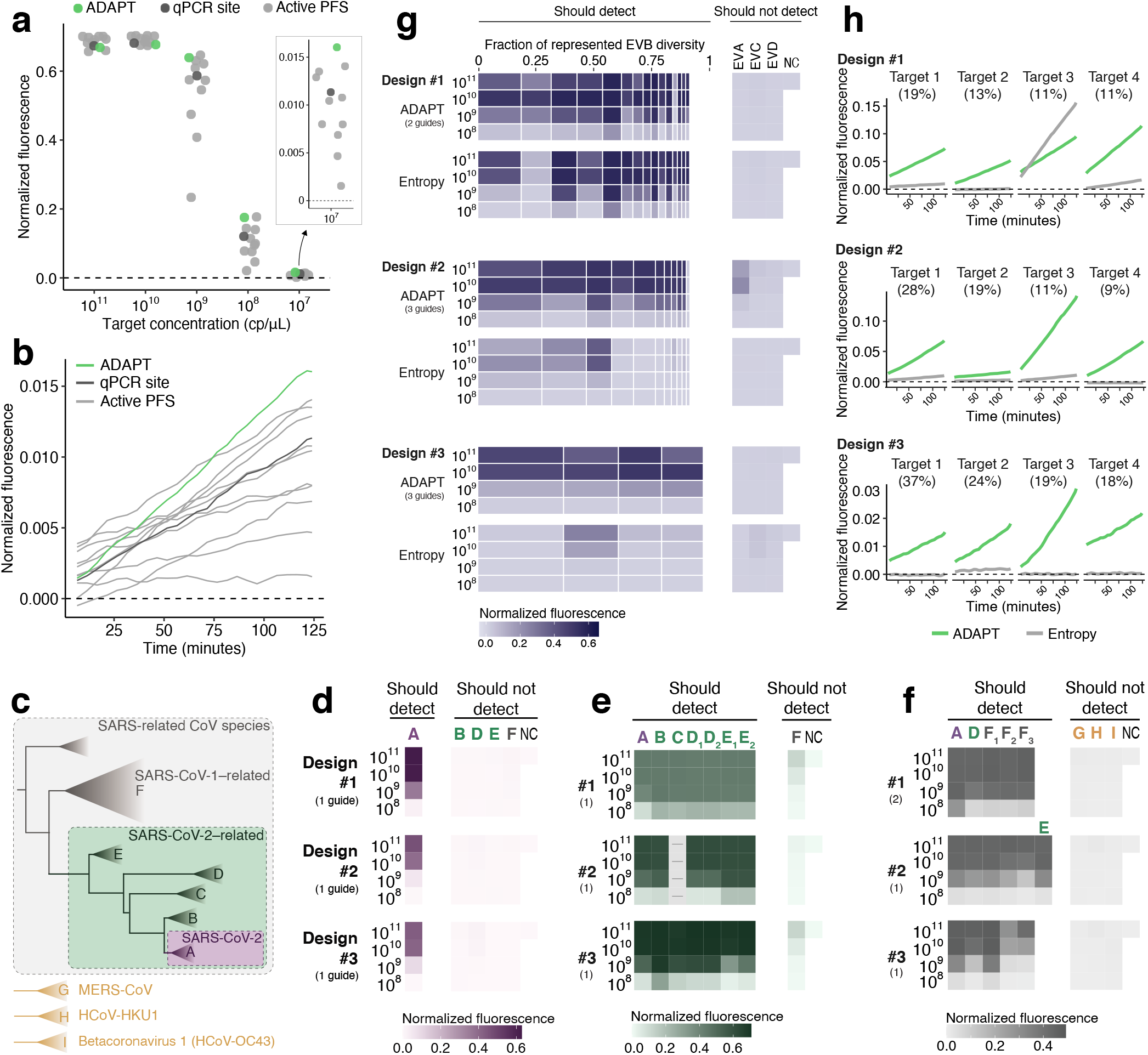
Experimental evaluation of designs. **(a)** Fluorescence at varying target concentrations in the US CDC’s SARS-CoV-2 N1 qPCR amplicon. Compared Cas13 guides are from our approach using ADAPT (green); a guide with an active (non-G) PFS at the site of the qPCR probe from the N1 assay (dark gray); and 10 randomly selected guides with an active PFS (light gray). All were constrained to the amplicon. Each point is one guide. **(b)** Fluorescence over time for guides from (a) at target concentration of 10^7^ cp/μL. (a) shows final time point (124 minutes). **(c)** Phylogeny within SARS-related CoV species based on ref. 47, and 3 related betacoronaviruses. **(d)** Fluorescence for ADAPT’s top-ranked SARS-CoV-2 designs in detecting representative targets of the clades in (c). Rankings, from 1 to 3, are by ADAPT’s predicted performance. Label to the left of each row indicates target concentration (cp/μL). NC, no template control. In (d–f), colors refer to (c). In (d–g), parenthetical numbers are the number of Cas13 guides in ADAPT’s design. **(e)** Same as (d), but for SARS-CoV-2–related taxon. Empty column with dashes indicates sequence in the design’s amplicon has high ambiguity and was not tested. Clades D and E sometimes have two representative targets in these amplicons; when two target labels were tested with one representative sequence, the same value is plotted for each. **(f)** Same as (d), but for SARS-related CoV species. Clade F_1_ is SARS-CoV-1, and F_2_ and F_3_ are related bat SARS-like CoVs. E requires a separate representative target in only one amplicon. G, H, and I are defined in (c). **(g)** Fluorescence for ADAPT’s top-ranked Enterovirus B (EVB) designs in detecting EVB and representative targets for Enterovirus A/C/D (EVA/C/D). Each band is an EVB target having width proportional to the fraction of EVB genomic diversity represented, within the amplicon of ADAPT’s design. Under each ADAPT design is one baseline guide (“Entropy”) from the site in the amplicon with an active PFS and minimal Shannon entropy. **(h)** Fluorescence over time at target concentration of 10^8^ cp/μL, for the 4 EVB targets encompassing the largest fraction of EVB genomic diversity. (g) shows middle time point (59 minutes). In all panels, fluorescence is reference-normalized and background-subtracted (Methods).

Next, we validated the comprehensiveness and specificity offered by our approach by considering multiple taxa comprising the SARS-related coronavirus (SARS-related CoV) species (Fig. 5c). We tested Cas13a-based designs against representative targets within each taxon, which we identified systematically according to sequence composition (Methods). Our testing directly measures the fluorescent signal yielded by the Cas13a guides at varying target concentrations. We started with precise targeting of SARS-CoV-2. Using ADAPT, we generated assays for detecting SARS-CoV-2, with lineage-level specificity, that should not detect any known bat or pangolin SARS-like coronaviruses, including the RaTG13 genome (96% identity to SARS-CoV-2^47^), nor SARS-CoV-1 and other coronaviruses (Fig. 5c). All three of our approach’s best design options (ranked according to predicted activity) detect SARS-CoV-2 with complete specificity—we observe no fluorescent signal for the closely related lineages (Fig. 5d).

We then broadened the targeted space within SARS-related CoV. We designed assays for the SARS-CoV-2–related lineage^48^, which consists of SARS-CoV-2 and its closely related bat and pangolin CoVs. Our approach’s three top-ranked designs sensitively and specifically detect all representative targets within SARS-CoV-2–related (Fig. 5e). Unlike the SARS-CoV-2 designs, we observed low off-target SARS-CoV-1 signal with the SARS-CoV-2–related designs because their added comprehensiveness antagonizes specificity; however, this is unlikely to affect diagnostic results that use an adequate signal threshold for detection. We also designed species-specific assays for the full SARS-related CoV species, and ADAPT’s top-ranked designs detect all representative targets within the species without any signal for three other betacoronaviruses (Fig. 5f).

Overall, all of our top-ranked designs—generated fully automatically and without any human input or experimental refinement—perform as desired across the SARS-related CoV taxa. For each taxon, ADAPT also generated additional design options (25 total) and we generally observe the desired activity; designs with poorer sensitivity or specificity are in the lower half of rankings according to predicted activity (Supplementary Fig. 32a–c). Four of the designs use two Cas13a guides and, in these cases, our measurements show that our combinatorial optimization algorithm selects them to detect distinct lineage groupings in order to maximize their collective sensitivity for the taxon (Supplementary Fig. 33).

We also evaluated limits of detection across extensive genomic variation. We focused on enteroviruses, which are estimated to cause millions of symptomatic infections yearly and frequent outbreaks^49^. They have over 100 types and likely many undiscovered types that cause human disease^50^. Testing increasingly relies on pan-enterovirus qPCR by targeting a highly conserved region, which has clinical value but limited surveillance utility^51^; an assay that provides more resolution than pan-enterovirus qPCR or serology, albeit less than sequencing, would aid surveillance.

We used ADAPT to design species-specific assays for Enterovirus B (EVB), which is widespread^52^ and exceptionally diverse, with 63 known types^50^. We found all three of ADAPT’s top-ranked designs detect the spectrum of genomic variation with specificity for EVB, as desired (Fig. 5g). To benchmark the efficacy of ADAPT’s approach, we targeted conserved sites by designing a guide within each amplicon at the site with minimal Shannon entropy and an active PFS (Methods). Targeting conserved sites is a standard, widely-used strategy for managing sequence diversity: conserved sites are a common target of CRISPR-based diagnostics^3,53^ for diverse viruses and, in particular, entropy commonly steers the design of qPCR assays^12, 54, 55^. The entropy-based strategy fails to detect many targets representative of EVB’s genomic diversity (Fig. 5g). By contrast, our approach in ADAPT provides a higher fluorescent signal in nearly all representative targets, enabling a lower limit of detection in about half of them (Fig. 5h and Supplementary Fig. 34a–c). In many design options we tested beyond the top three, the entropy-based strategy is more sensitive than ADAPT’s approach; however, in these cases the entropy-based strategy lacks species-specificity and ADAPT’s designs are ranked lower according to their predicted activities (Supplementary Fig. 35). Though ADAPT’s designs incorporate 1–3 guides and the entropy-based strategy uses one, we tested multiple entropy-based guides in five designs and found they exhibit similar activity at low target concentrations (Supplementary Fig. 34d–f and 36). Our results indicate that model-based optimization enables diagnostics that sensitively detect vast genomic diversity, including with improved sensitivity over a standard conservation-based strategy.

To further evaluate specificity in clinically relevant conditions, we performed an in silico comparison of all experimentally-valiated designs with the human transcriptome and 11 common bacterial pathogen genomes (Methods). All ADAPT-designed guides are at least 5 mismatches different from human transcripts and these bacterial genomes, indicating they are unlikely to produce detectable signal (Supplementary Fig. 16b). The same is true of all guides in our testing, except for one from the entropy-based strategy for benchmarking EVB detection (Design #8; 4 mismatches from a human transcript). The results demonstrate that, in addition to our designs being highly specific against related viruses, they are unlikely to cross-react with other non-targeted RNA in samples.

## Discussion

We developed an approach, combining a deep learning model with combinatorial optimization, to design viral diagnostics. We applied our approach using CRISPR-based diagnostics, for which we generated the largest dataset available on diagnostic signal and learned a model that quantitatively predicts detection efficiency. Critically, our approach directly integrates viral variation into an objective function to generate designs that are maximally active across viral variants. We built ADAPT, which runs our approach rapidly at scale, across thousands of viruses, and automatically integrates public viral genome data into the design process.

We applied our machine learning-based design approach and performed a rigorous experimental validation across extensive target variation, testing 69 diagnostic designs against a total of 290 different targets. The results show that ADAPT’s designs (i) exhibit significantly higher fluorescence for SARS-CoV-2 at low target concentrations than designs based on previously-used strategies; (ii) are sensitive and specific down to the lineage level across multiple closely-related taxa; and (iii) delineate a diverse species, Enterovirus B, with a lower limit of detection, across the spectrum of its genomic diversity, than a state-of-the-art design strategy focused on sequence conservation. While we tested extensively across viral variation, our results come from synthetic targets and we did not include clinical validation as part of our study. Validation on patient and environmental samples would be important before laboratories or field sites deploy ADAPT-designed assays in practice, although several prior studies using ADAPT-designed assays^8, 53, 56^ have demonstrated that they work well on such samples.

While this paper is the first to publish our approach, it has been helpful for prior design needs. In January, 2020, days after SARS-CoV-2 was first sequenced, we applied an early version of ADAPT to design a CRISPR-based SARS-CoV-2 diagnostic assay, and shortly after assays for 66 other viruses relevant to the COVID-19 pandemic^57^. Though designed from only the 20 genomes available at the time^58^, we predict this SARS-CoV-2 assay to detect 99.8% of the ~616,000 genomes sequenced through early March, 2021. Early versions of ADAPT were also applied to design diagnostic assays for 169 human viruses and influenza subtyping^8^, and separately for Lassa virus^53^.

We envision running ADAPT regularly for thousands of viruses, so that designs continually reflect their latest known variation. This will provide a resource of broadly-effective designs in advance of an outbreak, and assays for many strains could be proactively validated. Our results show that ADAPT’s designs perform well for known viruses, and can even be useful for novel viruses not yet known at the time of the design. For the latter case, however, our results also show that severe genome sampling biases can hinder ADAPT’s assays. The challenge is acute when a novel virus is distant from its nearest highly-sampled lineage, as with SARS-CoV-2. To mitigate that issue, it can help to synthesize multiple highly ranked assays rather than one, having them ready to compare on a novel virus. Indeed, our simulations of this application—in which we designed assays for the SARS-related coronavirus species prior to the discovery of SARS-CoV-2—showed that ADAPT’s second-highest–ranked design would outperform the highest-ranked one in detecting that novel virus.

There is room for methodological improvements in ADAPT. Developing and integrating into ADAPT a learned model for amplification primers, rather than using conventional primer heuristics, has the potential to improve the amplification steps of CRISPR-based diagnostic assays. However, such a model would also require expensive, time-consuming steps to construct a suitable training dataset for a particular amplification method, and recent developments in amplification-free CRISPR-based detection^46, 59, 60^ may negate the motivation for such work. Another area for progress is algorithmic. Improved design algorithms could solve particular, well-defined objectives that arise in viral contexts—such as optimally differentiating highly-homologous viral variants—better than our current algorithms; we are currently exploring the role of generative models for several objectives, and they fit well into ADAPT’s framework. And, rather than maximizing detection over a uniform distribution of known genomes, a principled approach that weighs genomes in the design process could help to correct for genome sampling biases and improve the chances that ADAPT’s designs sensitively detect under- or non-sampled emerging and novel viruses.

Our dataset and modeling of Cas13a collateral activity could be useful for understanding the biology of Cas13a. While we analyzed linear models to extract design considerations using a limited number of predefined features, a more thorough modeling of such features—like that performed for Cas13d^18^ using predicted RNA secondary structures among other crRNA and target RNA features—may reveal principles for Cas13a guide design beyond ones we describe. Though harder to interpret, our CNN models may learn novel features: ones we cannot predefine or that exhibit complex interactions missed by other models. Techniques for model interpretation, which have been applied to CNNs in genomics^61^, can elucidate important elements of the input sequences that promote high activity predictions; such learned features could form the basis of new design principles.

Our platform can be broadly applicable to detecting and responding to pathogens. Though we trained and applied a deep neural network for CRISPR-Cas13a, ADAPT flexibly accommodates machine-learned models that could be used with other nucleic acid diagnostic technologies. One example is qPCR, which remains widely used for viral diagnostics; COVID-19 qPCR assays exhibit high variability in their reported sensitivities^62^ and many target regions that have acquired mutations^32, 33, 63^, which suggests room for design improvement. ADAPT could design highly specific qPCR assays, including primers and probes, with maximal amplification efficiency over genomic variation, both for broad diagnostics and lineage typing. The design would be accomplished most effectively with a learned model for qPCR, but would also be possible using a model proxy based on existing, conventional hybridization criteria.

A web frontend for running ADAPT is available at https://adapt.guide. This resource, showing annotated visualizations of ADAPT’s output, also provides pre-designed assays against > 1,500 viruses, ready for experimental validation. Using ADAPT, those proactive designs can be continually updated to reflect the latest known variation across thousands of viruses. ADAPT is available as a software package at https://github.com/broadinstitute/adapt, and is written flexibly to support multiple applications.

Beyond viral diagnostics, our approach can benefit other efforts that require designing maximally active sequences from viral genome data. For example, genomic variation impacts the efficacy of sequence-based siRNA^64^ and antibody^65^ therapeutics. CRISPR-based antiviral development also requires deep consideration of sequence diversity, guide activity, and taxon-specificity^66^. These uses fit directly into our approach in ADAPT. ADAPT’s approach could also benefit sequence-based vaccine selection^67, 68^. Comparative genomic analyses demonstrate a considerable potential to improve vaccine antigen candidates for pathogens with high strain diversity, and comparative-genomics-informed design provides putative vaccine antigens with greater coverage than existing ones^69^. With appropriate models, model-based optimization may further improve upon comparative-genomics-informed design by rapidly designing antigens that yield maximal predicted antibody titers and T cell responses over viral strain diversity.

Our approach, together with the introduction of ADAPT, improves the development speed and efficacy of viral diagnostics, and has the potential to do so for other viral sequence-based technologies.

## Acknowledgements

We thank Yaron Singer, Mitchell O’Connell, Remy Tuyeras, and Daniel Kassler for fruitful discussions and pointers. This project was made possible by DARPA grant D18AC00006; HHMI; the AWS Diagnostic Development Initiative; Flu Lab; and a cohort of generous donors through TED’s Audacious Project, including the ELMA Foundation, MacKenzie Scott, the Skoll Foundation, and Open Philanthropy. H.C.M. was supported by NIH/NIAID grant K01 AI163498. N.J.H. was funded by the Landry Cancer Biology Consortium Fellowship and NIH/NIGMS grant T32 GM008313. M.M. was funded by NSF grants CCF-1535795 and CCF-1563710.

## Author contributions

H.C.M. initiated and led the study, and developed the algorithms and models in ADAPT, with advice from C.M., M.M., and P.C.S. H.C.M. and P.P.P. implemented ADAPT. N.J.H. designed the Cas13a guide-target library, tested it experimentally with C.M.A. and C.M., and analyzed its data with H.C.M. N.L.W., H.C.M., Y.B.Z., and P.P.P designed experiments evaluating ADAPT’s designs and analyzed results. N.L.W. performed those experiments with insight from P.C.B. and J.W. L.R., S.M., and D.K.Y. contributed to computational methods development and analyses. H.C.M. wrote the paper with contributions from N.L.W., C.M., M.M., and P.C.S., and feedback from all authors.

## Methods

### ADAPT

Supplementary Notes describe ADAPT’s algorithms, data structures, and implementation details. Supplementary Note 1 defines and solves objective functions. Supplementary Note 2 describes how ADAPT enforces specificity. Supplementary Note 3 describes how ADAPT searches for genomic regions to target and links with sequence databases.

### Introductory analyses

To illustrate viral database growth, we charted the growth in the number of viral genomes and their unique 31-mers over time (Supplementary Fig. 1). We first curated a list of viral species from NCBI^70^ known to infect humans (November 2019). For each, we took all NCBI genome neighbors^31^ (influenza sequences from the Influenza Virus resource^71^), which represent near-complete or complete genomes. To assign a date for each, we used the GenBank entry creation date rather than sample collection date for several reasons, including that this date more directly represents our focus in the analysis (when the sequence becomes present in the database) and that every entry on GenBank contains a value for this field. To control for some viruses having multiple segments (and thus sequences), we only used counts for one segment for each species, namely the segment that has the most number of sequences.

We used influenza A virus subtyping as an example to demonstrate the effect of evolution on diagnostics (Fig. 1a and Supplementary Fig. 2). We selected the most conserved *k*-mers—representing probe or guide sequences—from the sequences available at different years. Here, for simplicity, we ignored all other constraints, such as detection activity and specificity (the latter of which is critical for subtyping), which would further degrade the temporal performance of the selected *k*-mers. In particular, for each design year *Y*, we selected the 15 non-overlapping 30-mers found in the largest number of sequences taken from the two most recent years (*Y* – 1 and *Y*). We then measured the fraction of sequences in subsequent test years (*Y, Y* + 1, …) that exactly contain each of these *k*-mers. We performed the design strategy over 10 resamplings of the sequences and use the mean fraction. We repeated this 4 times: for segment 4 (HA) sequences of H1 and H3 subtypes, and segment 6 (NA) sequences of N1 and N2 subtypes.

To visualize mutations accumulating on a genome during the course of an outbreak (Fig. 1b), we used complete SARS-CoV-2 genomes from GISAID^58^. We called variants in all genomes, through 2020, against the reference genome ‘`hCoV-19/Wuhan/IVDC-HB-01/2019`’ (GISAID accession ‘`EPI - ISL_402119`’). For every date *d* between February 1, 2020 and January 1, 2021, spaced apart by one month, at every position we calculated the fraction of all genomes collected up to *d* that have a variant against the reference. We called all variants present between 0.1% and 1% frequency on some *d* as “low” frequency and variants at ≥ 1% frequency on some *d* as “high” frequency. We ignored all variants present at ≥ 1% frequency on the initial *d* (ancestral) or that were both low frequency on the initial *d* and stayed low frequency by the final *d*—i..e, we kept the variants that transitioned to low or high frequency by the final *d*. We show the *d* when the variant first becomes called as low (light purple) or high (dark purple) frequency. If a variant transitions both to low and then to high frequency by the final *d*, we only show it for the *d* when it becomes high frequency.

### Cas13a library design and testing

We designed a collection of CRISPR-Cas13a crRNA guides and target molecules to evaluate guide-target activity, focusing on assessing likely-active guide-target pairs. First, we designed a target (the *wildtype* target) that is 865 nt long (design details for the wildtype target are in the subsequent paragraph). We then created 94 guides (namely, the 28 nt spacers) tiling this wildtype target (Fig. 2a and Supplementary Fig. 3a). In the tiling scheme there are 30 nt blocks, each having 4 overlapping guides, in which the starts of the 3 guides, from the start of the most 5’ guide, are 4 nt, 13 nt, and 23 nt. Of the 94 guides, 87 are experimental, 3 are negative controls, and 4 are positive controls. We created 229 unique target sequences: 1 of them is the wildtype sequence (guides should exhibit activity against this target), 225 are experimental (mismatches and varying PFS alleles against the guides), and 3 are negative controls. All guides exactly match the wildtype target and should detect this, except the 3 negative control guides, which are not intended to detect any targets except one of the 3 negative control targets each. The 4 positive control guides target 4 30-nt regions with a perfectly complementary sequence and non-`G` PFS that are held constant across all targets, with the exception of the 3 negative control targets. Across the experimental targets, the mismatches profile varying choices of positions and alleles against the guide. For the experimental targets, we generated single mismatches evenly spaced every 30 nt along the experimental region such that every guide targeting this region has either a single mismatch or an altered PFS at +1 or +2 nt from the protospacer; we created a total of 45 (3 × 15) such targets to probe all 3 possible mismatch alleles and 15/30 of the possible phasings. In the remainder of the experimental targets, we generated targets with 2, 3, or 4 mismatches per 30 nt block with respect to the guide RNA in phase with the block. For these targets, we randomly selected mismatch positions to uniformly sample (or, when possible, exhaustively enumerate) average mismatch spacing and average mismatch distance to the center of the spacer, and randomly selected mismatch alleles. The 87 experimental guides may detect up to 226 unique target sequences (the wildtype and 225 experimental targets), providing 19,662 experimental guide-target pairs.

To construct the wildtype target sequence, we aimed to produce a composition spanning viral genomic sequence diversity. In particular, we started with a previously-described dataset of genomes from human-infecting viral species^72^, constructed a vector of the dinucleotide frequencies for each species, and performed principal component analysis of the species from these vectors. For each 30 nt block of the wildtype target, we selected a point from the space of the first 3 principal components (uniformly at random), reconstructed a corresponding vector of dinucleotide frequencies (i.e., transformed the point back to the original space), and then iteratively selected every next nucleotide in the block according to the distribution of dinucleotides. A goal of this scheme is for dinucleotides that are variable across viral species to also vary in frequency across the wildtype target: a dinucleotide that explains considerable variance across viral species (e.g., is rich in some viral species and poor in others) ought to be rich in some blocks of the wildtype target and poor in other blocks, whereas a dinucleotide that explains little variance across species ought to have similar frequency along the target. In positions that would serve as a PFS site for a guide, we disallowed `G`, and proportionately adjusted upwards the probability of choosing a `G` in non-PFS positions to maintain the total dinucleotide frequency in accordance with the randomly selected distribution (mismatches in experimental targets can still introduce a `G` PFS).

We synthesized the targets as DNA, in vitro transcribed them to RNA, and synthesized the crRNAs as RNA. On all crRNAs, we used the same direct repeat (GAUUUAGACUACCCCAAAAACGAAGGGGACUAAAAC). To determine a reasonable concentration for measuring fluorescence over time points, we tested 8 concentrations of 2 targets and 2 guides in a pilot experiment (Supplementary Fig. 4a) and proceeded with 6.25 × 10^9^ cp/μL. We tested the library using CARMEN, a droplet-based Cas13a system; we followed the methodology described in ref. 8, which also contains the protocol. Briefly, a guide-target pair is enclosed in a droplet, together with the Cas13a enzyme, that may result in a detection reaction and thus fluorescence. We took an image of each location on each chip roughly every 20 minutes to measure this fluorescence. To alleviate the presence of microdroplets in this experiment (i.e., an irregular pairing of target and guide; about 1/3 of the droplets), we trained and applied a convolutional neural network on hand-labeled data to identify and remove these.

### Quantifying activity

In our Cas13a detection experiments, a fluorescent reporter is cleaved over time and its cleavage follows first-order kinetics:

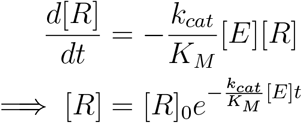

where [*R*] is the concentration of the not-yet-cleaved reporter, [*E*] is the concentration of the Cas13a guide-target complex, 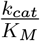 is the catalytic efficiency of the particular guide-target complex, and *t* is time. The fluorescence measurements that we make, *y,* are proportional to the quantity of cleaved reporter at some time point:

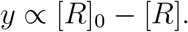

Therefore, for each guide-target complex we fit a curve of the form

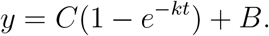

Here, *C* and *B* represent the saturation point and background fluorescence, respectively. *k* represents the rate at which the reporter is cleaved, and it is proportional to the catalytic efficiency of the particular guide-target complex:

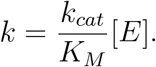

This relationship is validated by the linear relationship between *k* and [*E*] (Supplementary Fig. 4a) when we vary the concentration of target (the limiting component of the complex). In producing our dataset, we held [*E*] constant. We used log_10_(*k*) as our measurement of guide activity (Fig. 2 and Supplementary Fig. 4a,b). Intuitively, each step-increase in log_10_(*k*) corresponds to a fold-decrease in the half-life of the reporter in the reaction.

Our experimental data incorporates multiple droplets for each guide-target pair (Supplementary Fig. 5a). Each droplet represents one technical replicate of a particular guide-target pair. Thus, we have fluorescence values for each replicate at different time points, and in practice we compute the activity log_10_(*k*) for each replicate.

We curated the data to obtain a final dataset. Namely, we discarded data from two guides that showed no activity between them and any targets, owing to low concentrations in their synthesis. We also did not use data from positive or negative control guides, nor from the negative control target. Our final dataset contains 19,209 unique guide-target pairs (Supplementary Fig. 3b,c), counting 20 nt of sequence context around each protospacer in the target (18,253 unique pairs when not counting context).

Most guide-target pairs show activity (Supplementary Fig. 5d), as expected. At small values of *k* on a limited time scale (*t* up to ~120 minutes), we do not observe reporter activation (Supplementary Fig. 4b). Moreover, the curve becomes approximately linear (first order Maclaurin expansion: *y* ≈ *Ckt* + *B*). At such values of *k,* we cannot estimate both *C* and *k* together; intuitively, this is because there is too little detectable signal. Therefore, there is a cutoff at which we can estimate *k*; we labeled activities at log(*k*) > −4 as active, and the others as inactive. This phenomenon also implies that at smaller values of *k*, including ones we label as active, activity estimates might be less reliable.

### Predicting detection activity

#### Measurement error

To account for measurement error, we sampled, with replacement, 10 technical replicate measurements of activity for each guide-target pair (Supplementary Fig. 5a). We used this strategy to ensure that, although there are differing numbers of replicates per guide-target pair, each pair would be represented in the dataset with the same number of replicates. There are 19,209 × 10 = 192,090 points in total in our dataset that we use for training and testing. When plotting regression results on guide-target pairs in the test set (Fig. 2e, Supplementary Fig. 11a, Supplementary Fig. 13), we set the true activity of a pair to be the mean of the measured activities across the technical replicates for the pair.

#### Model and input descriptions

We approached prediction using a two-step hurdle model, reasoning that (i) separate processes govern whether a guide-target pair is active compared to the level of its activity; and (ii) we could better predict the activity of active pairs if we exclude the inactive pairs from a regression. We developed a classifier to decide whether a pair is inactive or active, and a regression model to predict the activity of only active pairs.

We explored multiple models for classification (Fig. 2c and Supplementary Fig. 6a), each with a space of hyperparameters:

- L1 logistic regression: regularization strength (logarithmic in [10^-4^,10^4^])
- L2 logistic regression: regularization strength (logarithmic in [10^-4^,10^4^])
- L1+L2 logistic regression (elastic net): regularization strength (logarithmic in [10^-4^,10^4^]), L1/L2 mixing ratio (1.0 – *2^x^* + 2^-5^ for *x* uniform in [−5, 0])
- Gradient-boosted trees (GBT): learning rate (logarithmic in [10^-2^, 1]), number of trees (logarithmic in [1, 2^8^], integral), minimum number of samples for splitting a node (logarithmic in [2,2^3^], integral), minimum number of samples at a leaf node (logarithmic in [1,2^2^], integral), maximum depth of a tree (logarithmic in [2, 2^3^], integral), number of features to consider when splitting a node (for *n* features, chosen uniform among considering all, 0.1*n*, 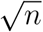, and log_2_ *n*)
- Random forest (RF): number of trees (logarithmic in [1, 2^8^], integral), minimum number of samples for splitting a node (logarithmic in [2, 2^3^], integral), minimum number of samples at a leaf node (logarithmic in [1, 2^2^], integral), maximum depth of a tree (chosen uniformly among not restricting the depth or restricting the depth to a value picked logarithmically from [2, 2^4^] and made integral), number of features to consider when splitting a node (for *n* features, chosen uniform among considering all, 0.1 *n*, 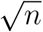, and log_2_ *n*)
- Support vector machine (SVM; linear): regularization strength (logarithmic in [10^-8^, 10^8^]), penalty type (chosen uniformly among L1 and L2)
- Multilayer perceptron (MLP): number of layers excluding the output layer (uniform in [1, 3]), dimensionality of each layer excluding the output layer (each chosen uniformly in [4, 127]), dropout rate in front of each layer (uniform in [0, 0.5]), activation function (chosen uniformly among ReLU and ELU), batch size always 16
- Long short-term memory recurrent neural network (LSTM): dimensionality of the output vector (logarithmic in [2, 2^8^], integral), whether to be bidirectional (chosen uniformly among unidirectional and bidirectional), dropout rate in front of the final layer (uniform in [0, 0.5]), whether to perform an embedding of the one-hot encoded nucleotides and the dimensionality if so (chosen with 1/3 chance to not perform an embedding, and with 2/3 chance to perform an embedding with dimensionality chosen uniformly in [1, 8]), batch size is always 16
- Convolutional neural network (CNN): number of parallel convolutional filters and their widths (chosen uniformly among not having a convolutional layer, 1 filter of width 1, 1 filter of width 2, 1 filter of width 3, 1 filter of width 4, 2 filters of widths {1, 2}, 3 filters of widths {1, 2, 3}, and 4 filters of widths {1, 2, 3, 4}), convolutional dimension (uniform in [10, 249]), pooling layer width (uniform in [1, 3]), pooling layer computation (chosen uniformly among maximum, average, and both), number of parallel locally connected layers and their widths (chosen uniformly among not having a locally connected layer, 1 filter of width 1, 1 filter of width 2, and 2 filters of widths {1, 2}), locally connected filter dimension (uniform in [1, 4]), number of fully connected layers and their dimensions (chosen uniformly among 1 layer with dimension uniform in [25, 74] and 2 layers each with dimension uniform in [25, 74]), whether to perform batch normalization in between the convolutional and pooling layers (uniform among yes and no), activation function (chosen uniformly among ReLU and ELU), dropout rate in front of the fully connected layers (uniform in [0, 0.5]), L2 regularization coefficient (lognormal with mean *μ* = −13, *σ* = 4), batch size (uniform in [32, 255]), learning rate (logarithmic in [10^-6^, 10^-1^])

Similarly, for regression we explored multiple models (Supplementary Fig. 6b,c), each with a space of hyperparameters:

- L1 linear regression: regularization strength (logarithmic in [10^-8^, 10^8^])
- L2 linear regression: regularization strength (logarithmic in [10^-8^, 10^8^])
- L1+L2 linear regression (elastic net): regularization strength (logarithmic in [10^-8^, 10^8^]), L1/L2 mixing ratio (1.0 – 2^*x*^ + 2^-5^ for *x* uniform in [−5, 0])
- Gradient-boosted trees (GBT): same hyperparameter space as for classification
- Random forest (RF): same hyperparameter space as for classification
- Multilayer perceptron (MLP): same hyperparameter space as for classification
- Long short-term memory recurrent neural network (LSTM): same hyperparameter space as for classification
- Convolutional neural network (CNN): same hyperparameter space as for classification

Model selection and evaluation describes the search process.

When training and testing the models, we used 28 nt guide and target sequence, and include 10 nt of context in the target sequence on each side of the protospacer. We tested the following different inputs:

- ‘One-hot (1D)’: vector containing 4 bits to encode the nucleotide at each target position and 4 bits similarly for each guide position; with a 28 nt guide and 10 nt of context in the target around the protospacer, there are (10 + 28 + 10 + 28) × 4 = 304 bits
- ‘One-hot MM’: similar to ‘One-hot (1D)’ except explicitly encoding mismatches between the guide and target—i.e., vector containing 4 bits to encode the nucleotide at each target position and 4 bits, at each guide position, encoding whether there is a mismatch (if not, all 0) and, if so, the guide allele; same length as ‘One-hot (1D)’
- ‘Handcrafted’: features are count of each nucleotide in the guide, count of each dinucleotide in the guide, GC count in the guide, total number of mismatches between the guide and target sequence, and a one-hot encoding of the 2-nt PFS (coupling the 2 nucleotides); the number of features are 4 + 16 + 1 + 1 + 16 = 38
- ‘One-hot MM + Handcrafted’: concatenation of features from ‘One-hot MM’ and ‘Handcrafted’, except removing from ‘One-hot MM’ the bits encoding the 2-nt PFS because these are included in ‘Handcrafted’

We used these inputs for all models except the LSTM and CNN. For these two models, which can capture and extract spatial relationships in the input, we used an alternative input (labeled ‘One-hot (2D)’ in figures). Here, the input dimensionality is (48, 8) and consists of a concatenated one-hot encoding of the target and guide sequence. Namely, each element *x_i_* (*i* ∈ {1 … 48}) is a vector [*x_i,t_, x_i,g_*]. Target context corresponds to *i* ∈ {1 … 10} (5’ end) and *i* ∈ {39 … 48} (3’ end); for these *i, x_i,t_* is a one-hot encoding of the target sequence and *x_i,g_* is all 0. The guide binds to the target at *i* ∈ {11 … 38} and, for these *i, x_i,t_* is a one-hot encoding of the target sequence protospacer at position *i* – 10 of where the guide is designed to bind, while *x_i,g_* is a one-hot encoding of the guide at position *i* – 10.

We evaluated all models, except the MLP, LSTM, and CNN, in scikit-learn 0.22^73^. We implemented and evaluated the MLP, LSTM, and CNN models in TensorFlow 2.1.0^74^.

For the MLP, LSTM, and CNN models, we used binary cross-entropy as the loss function for classification and mean squared error for regression. For these 3 models, we used the Adam optimizer^75^ and performed early stopping during training (maximum of 1,000 epochs) with a held-out portion of the training data. Additionally, for the CNN we regularized the weights (L2). When training all classification models, we weighted the active and inactive classes equally.

#### Data splits and test set

In evaluating our models, we must determine folds of the data and pick a held-out test set. One challenge is that, in our design, guides overlap according the position against which they were designed along the wildtype target. Although effects on activity might be position-dependent within the guide, this overlap can cause guides to have similar sequence composition or to be in regions of the target sequence with similar structure. To remove this possibility of leakage between a data split, after making a split of *X* into *X*_train_ and *X*_test_, we remove all guide-target pairs from *X*_test_ for which the guide has any overlap, in target sequence they are designed to detect, with a guide in *X*_train_. We perform this strategy during all cross-validated analyses. We also use it to choose a test set that we hold out from all analyses and use only for evaluating the final CNNs. This test set consists of 30% of all guides (counted before removing overlaps between *X*_train_ and *X*_test_), taken from the far 3’ end of the target.

#### Model selection and evaluation

We performed nested cross-validation to select models—both for classification and regression—and evaluate our selection of them (Fig. 2c and Supplementary Fig. 6). We used 5 outer folds of the data. For each outer fold, we searched for hyperparameters using a cross-validated (5 inner folds) random search over the space defined in Model and input descriptions; we scored using the mean auROC (classification) or Spearman correlation (regression) over the inner folds. In each random search, we used 100 hyperparameter choices for all models, except for the LSTM and CNN models (50), which we found slower to train.

The CNN models outperformed others in the above analysis, so we selected a final CNN model for classification and another for regression. For each of classification and regression, we performed a random search across 5 folds of the data using 200 random samples. We selected the model with the highest auROC (classification) or Spearman correlation (regression) averaged over the folds. Our evaluations of these two models used the held-out test set.

#### Incorporating into ADAPT

We integrated the CNN models into ADAPT. First, we set the decision threshold on the classifier’s output to be 0.577467. We chose the threshold, via cross-validation, to achieve a desired precision of 0.975. In particular, we took 5 folds of our data (excluding test data) and, for each fold, we calculated the threshold that achieves a precision of 0.975 on the validation data. Our decision threshold is the mean across the folds.

We then defined a piecewise function, incorporating the classification and regression models, as:

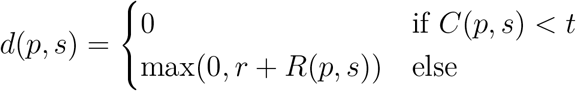

where *d*(*p, s*) is the predicted detection activity between a probe *p* and target sequence *s*(*s* includes 10 nt of context). *C*(*p, s*) is the output of the classifier, *t* is the classification decision threshold, and *R*(*p, s*) is the output of the regression model. *r* is a shift that we add to regression outputs to ensure *d*(*p, s*) is non-negative; though a nice property, it is not strictly needed as long as we constrain the ground set as described in Supplementary Note 1a. The choice of *r* should depend on the range of activity values in the dataset; here, *r* = 4.

#### Comparison of predictions with independent Cas13a datasets

We compared our model’s predictions with data from ref. 37. Although our model was trained for LwaCas13a whereas the data in ref. 37 is from LbuCas13a, the comparison enables validation on an independent dataset and the opportunity to assess whether RNA binding activity correlates with collateral (or *trans*-) cleavage activity. We padded the 20-nt spacer sequences used in the data with 8 randomly determined bases because our model requires 28-nt spacer sequences; in particular, we added 5’-AAAATCTG-3’ to the 3’ end of the crRNA spacer sequences, with matching complementary bases in the target sequences. Additionally, we padded the target sequences with random bases to obtain 10-nt on each side of the protospacer, as required by our model. In the RNA binding data: for crRNA X (‘X’ and ‘Y’ are as in ref. 37), we added `5’-AAGCATG-3’` to the 5’ end of the corresponding target sequence and `5’-TGCCATA-3’` to the 3’ end; for crRNA Y, we added `5’-GATTCAA-3’` to the 5’ end of the corresponding target sequence and 5’-CAGCATA-3’ to the 3’ end. In the collateral cleavage activity data: we added 5’-CATAT-3’ to the 3’ end of crRNA X’s corresponding target sequence (the 5’ end has a sufficient number of context bases for our model, and crRNA Y was not a part of this experiment).

Along with padding the data from ref. 37 for input to our model, we made several other choices when handling the data from ref. 37 to allow for comparisons. For LbuCas13a–RNA binding measurements, we normalized the data as done in ref. 37. That is, we used fold-changes (ratio of read counts relative to apo-Cas13a, i.e., protein with no guide), and then normalized them by subtracting average fold-change for the off-target (for crRNA X, crRNA Y’s target; for crRNA Y, crRNA X’s target) and then dividing by the average fold-change for the on-target (for crRNA X, crRNA X’s target, and likewise for crRNA Y). In our comparison, we skipped data points where the fold-change could not be computed (reported as ‘nan’; possibly, the control (apo-Cas13a) had 0 reads). To correct for substantial differences in the numbers of mismatches across data points, we randomly downsampled to 150 data points for each number of mismatches (the number of mismatches is a strong predictor of activity, and not downsampling would cause data points with fewer than 3 mismatches to be a small component of the comparison); we chose 150 because there are 156 data points with 1 mismatch, and more than 150 for each choice of ≥ 2 mismatches. For collateral cleavage measurements, we used 0 as the value from from ref. 37 (the normalized cleavage rate relative to no mismatches) for two data points labeled ‘ND’.

When comparing measured LbuCas13a cleavage rates from ref. 37 to our model’s predictions of LwaCas13a collateral activity, we also subsetted the data. On the full cleavage rate data (1 guide, 26 targets), our model’s predictions show a weak and non-significant correlation with the measurements (Spearman’s *ρ* = 0.360 with *p* = 0.07; Supplementary Fig. 15a). LbuCas13a’s greater overall cleavage activity^34^, compared to LwaCas13a, could explain the discrepancy. Thus, we also considered only the guide-target pairs where mismatches harm activity, removing the 8 mismatched targets where LbuCas13a exhibits even greater activity than against a matching target (Supplementary Fig. 15b).

We also compared our model’s predictions with measurements (mean across replicates) of on-target (*cis*-) cleavage, namely LwaCas13a knockdown, from Figure 3a–c in ref. 36. For the target sequence of the Gluc guide 1 (defined in ref. 36), we used sequence from GenBank accession MF882921; for the *CXCR4* guide, we used sequence from [NM_001008540](https://www.ncbi.nlm.nih.gov/nuccore/MF882921); and for the Gluc guide 2, we used sequence from MF882921. To insert mismatches into spacers, we randomly selected an allele for the mismatch at the positions given in the data.

For all comparisons, when making predictions on these data with our model, we used the hurdle model (piecewise function) described in Incorporating into ADAPT with the same high-precision decision threshold on the classifier that we use in ADAPT. When reporting *p*-values for Spearman’s test and Pearson’s test, the alternative hypothesis is that the true correlation is not 0 (Pearson’s test uses a *t*-distribution).

### ADAPT analyses

#### Comparing algorithms for submodular maximization

To compare the canonical greedy algorithm for constrained monotone submodular maximization^43^ with the fast randomized combinatorial algorithm^42^ (Supplementary Fig. 18), we ran ADAPT 5 times under each choice of parameter settings and species and, for each run, considered the mean final objective value taken across the best 5 design options. We used the arguments ‘`-pm 3 -pp 0.9 --primer-gc-content-bounds 0.3 0.7 --max-primers- at-site 10 -gl 28 --max-target-len 250`’with our Cas13a activity model. We used the default objective function in ADAPT: 4 + *A* – 0.5*P* – 0.25*L*, where *A* is the expected activity of the guide set, *P* is the number of primers, and *L* is the target length.

#### Benchmarking comprehensiveness

To benchmark comprehensiveness (Fig. 3b,c and Supplementary Fig. 19), we ran ADAPT with three approaches. In all approaches, we decided that a probe detects a target sequence if and only if they are within 1 mismatch, counting G-U wobble pairs as matches; and used a sliding window of 200 nt and a probe length of 30 nt. We used bootstrapping to estimate uncertainty around plotted values owing to viral genome sampling: 5 times, we randomly sampled with replacement from all NCBI genome neighbors^31^ for each species (if there are *N* neighbors, we randomly sampled *N* with replacement) and used each of these resamplings as input to 5 runs. In the first approach (baselines), we used ADAPT’s `design_naively.py` program to select probes within each window via three strategies: (1) the consensus probe, computed at every site within the window, that detects the most number of genome sequences (‘consensus’); (2) the most common probe sequence, determined at every site within the window, that detects the most number of genome sequences (‘mode’); and (3) the *n* most common subsequences, with all *n* determined at each site in the window, choosing the *n* from the site where they collectively detect the most number of genome sequences (doing this separately for *n* ranging from 1 to 10). In the second approach, we maximized expected activity using ADAPT across the target sequences with different numbers of probes (hard constraints) using a penalty strength of 0 (i.e., no soft constraint). Here, we defined the activity to be binary: 1 for detection, and 0 otherwise; this has the property that expected activity is equivalent to the fraction of sequences detected. In the third approach, we use the objective function in ADAPT that minimizes the number of probes subject to constraints on the fraction of sequences detected (specified via ‘`-gp`’; 0.9, 0.95, and 0.99).

#### Evaluating dispersion and generalization

We evaluated the dispersion, owing to randomness and sampling, in ADAPT’s designs (Supplementary Fig. 25). In all cases, we used all NCBI genome neighbors^31^ for each species and used the following arguments with ADAPT: ‘--obj maximize-activity --soft-guide-constraint 1 --hard-guide-constraint 5 --penalty-strength 0.25 -gl 28 -pl 30 -pm 3 -pp 0.98 --primer-gc-content-bounds 0.35 0.65 --max-primers-at-site 10 --max-target-length 500 --obj-fn-weights 0.50 0.25’, with a cluster threshold such that there is only 1 cluster, and used our Cas13a activity model. We ran ADAPT in two ways: 20 times without changing the input (output differences are owing to randomness) and 20 times with resampled input (output differences are owing both to randomness and sampling of the input sequences). Then, we measured dispersion by treating the 5 highest-ranked design options from each run as a set and computing pairwise Jaccard similarities across the 20 runs. This computation requires us to evaluate overlap between two sets: in one comparison, we consider a design option *x* to be in another set if *x* is present exactly in that other set (same primers and probes) and, in the other comparison, we consider a design option *x* to be in another set if that other set has some design option with both endpoints within 40 nt of *x*’s endpoints.

To evaluate the generalization of ADAPT’s designs (Fig. 4b), we performed cross-validation via repeated random subsampling. For each species, we took all NCBI genome neighbors^31^ and, 20 times, randomly selected 80% of them to use as input for design and the remaining 20% to test against. For each split, we used the same arguments with ADAPT as when evaluating dispersion: ‘--obj maximize-activity --soft-guide-constraint 1 --hard-guide-constraint 5 --penalty-strength 0.25 -gl 28 -pl 30 -pm 3 -pp 0.98 --primer-gc-content-bounds 0.35 0.65 --max-primers-at-site 10 --max-target-length 500 --obj-fn-weights 0.50 0.25’, with a cluster threshold such that there is only 1 cluster, and used our Cas13a activity model. When computing the fraction of sequences in the test set that are detected, we required the sequence to be detected by a primer on the 5’ and 3’ ends of a region (within 3 mismatches) and a probe (here, guide) to detect the region; we used the `analyze_coverage.py` program in ADAPT for this computation. We labeled detection of a sequence as “active” if a guide in the guide set is decided by our Cas13a classification model to be active against the target. We labeled the detection as “highly active” if a guide in the guide set is both decided to be active by the Cas13a classification model and its predicted activity, according to the Cas13a regression model, is ≥ 2.7198637 (4 added to the output of the model, −1.2801363). This threshold corresponds to the top 25% of predicted values on the subset of our held-out test set that is classified as active.

Using the same cross-validation strategy, we also evaluated generalization except with relaxed settings on constraints for the number of guides and more stringent settings on primer coverage (Supplementary Fig. 26): ‘--obj maximize-activity --soft-guide-constraint 3 `--hard-guide-constraint 10 --penalty-strength 0.05 -gl 28 -pl 30 -pm 3 -pp 0.995 --primer-gc-content-bounds 0.20 0.80 --max-primers-at-site 15 --max-target-length 1000 --obj-fn-weights 0.30 0.05`’. These settings allow for more complex assay designs (e.g., more guides and primers) to enable a higher sensitivity. Additionally, when deciding detection of the held-out genomes in this analysis, we adjusted thresholds to allow a higher sensitivity with lower precision: we allowed 4 mismatches for primers (instead of 3) and lowered the decision threshold of our Cas13a classification model to 0.3 (instead of 0.577467).

#### Benchmarking trie-based specificity queries

We benchmarked the approach described in Supplementary Note 2d against a single, large trie (Supplementary Fig. 22). For this, we sampled 1.28% of all 28-mers from 570 viral species (~78.7 million 28-mers in total), and built data structures indexing these. We then randomly selected 100 species (here, counting each segment of a segmented genome as a separate species), and queried 100 randomly selected 28-mers from each of these for hits against the other 569 species. We performed this for varying choices of mismatches. We used the same approach to generate results in Supplementary Fig. 20, there comparing queries with and without tolerance of G-U base pairing.

#### Benchmarking runtime improvement with memoization

We benchmarked the effect on runtime of memoizing repeated computations (Supplementary Fig. 23), as described in Supplementary Note 3b. We used all genome neighbors from NCBI’s viral genomes resource^31^ as input for each of the three species tested. To run ADAPT while memoizing computations, we used the arguments: ‘--obj maximize-activity --soft-guide-constraint 1 --hard-guide-constraint 5 --penalty-strength 0.25 --maximization-algorithm random-greedy -pm 3 -pp 0.9 --primer-gc-content-bounds 0.3 0.7 --max-primers-at-site 10 -gl 28 --max-target-len 250 --best-n-targets 10 --id-m 4 --id-frac 0.01 --id-method shard’. We also used our Cas13a predictive model and enforced specificity against all other species within each species’s family. To perform runs without memoizing computations, we did the same except added the argument ‘`--do-not-memoize-guide-computations`’, which skips all memoization steps during ADAPT’s search (except for calls to the predictive model).

#### Design of broadly-effective SARS-related coronavirus assays in 2018 and their evaluation

To evaluate the efficacy of species-level assays on a novel virus (Supplementary Fig. 28), we focused on SARS-related coronavirus. We simulated the 2018 design of assays for detecting the SARS-related CoV species, roughly a year before the initial detection of SARS-CoV-2. In particular, we used as input all genome neighbors from NCBI’s viral genomes resource^31^ for SARS-related CoV that were released on or before December 31, 2018 (there are 311 genomes). For ADAPT’s designs, we used the same parameters used for the vertebrate-infecting viral species designs (Designs across vertebrate-infecting species), except tolerating up to 1 mismatch between primer and target sequences; the specificity criteria were also the same as in those designs.

In 2018, SARS-related CoV was biased toward SARS-CoV-1 genomes (owing to SARS outbreak sequencing) relative to viruses sampled from animals. To alleviate this overrepresentation, we also produced designs using ADAPT in which the input downsampled SARS-CoV-1 to a single genome (Supplementary Fig. 28c,d). We used the RefSeq, GenBank accession AY274119, as that genome.

To determine the performance of these designs on SARS-CoV-2, we used the 184,197 complete genomes (low-quality removed) available on GISAID^58^ as of November 12, 2020. For an assay to be predicted to detect a target sequence (Supplementary Fig. 28b,d), we require that (i) primers on both ends are within 3 mismatches of the target sequence; and (ii) a guide in the guide set is classified by our Cas13a predictive model as active. We used this criteria both for evaluating detection of SARS-CoV-2 and of the design’s input.

### Designs across vertebrate-infecting species

We found all viral species in NCBI’s viral genomes resource^31^ that have a vertebrate as a host, as of April, 2020. These are species ratified by the International Committee on Taxonomy of Viruses (ICTV)^76^. We added to this list others that may have been incorrectly labeled, as well as influenza viruses, which are separate from the resource. There were 1,933 species in total and we used ADAPT to design primers and Cas13a guides to detect them. As input, we used all genome neighbors from NCBI’s viral genomes resource^31^ (influenza database for influenza species^71^). We ran ADAPT in May–June, 2020, and thus the input incorporates sequences available through those dates.

We constrained primers to have a length and GC content that are recommended for use with RPA^77^, and thus are suitable for use with SHERLOCK1 detection. We enforced specificity at the specieslevel within each family. That is, we required that the guides for each species not have off-target hits to sequence from any other species in its same family. Restricting our specificity queries to one family at a time reduces ADAPT’s memory usage and runtime.

We used the following arguments when running ADAPT to maximize expected activity:

- Initial clustering: clustered with a maximum distance of 30% (`--cluster-threshold 0.3`)
- Primers and amplicons: primer length of 30, primers must have GC content between 35% and 65%, at most 10 primers at a site^1^, up to 3 mismatches between primers and target sequence for hybridization, primers must hybridize to ≥ 98% of sequences, and length of a targeted genome region (amplicon) must be ≤ 250 nt (-`pl 30 --primer-gc-content-bounds 0.35 0.65 --max-primers-at-site 10 -pm 3 -pp 0.98 --max-target-length 250`)
- Guides: Cas13a guide length of 28 nt, together with our Cas13a predictive model (`-gl 28 --predict-activity-model-path models/classify/model-51373185models/regress/model-f8b6fd5d`)
- Guide activity objective: soft constraint of 1 guide, hard constraint of 5 guides, guide penalty (*λ*) of 0.25, using the randomized greedy algorithm (`--obj maximize-activity --soft-guide-constraint 1 --hard-guide-constraint 5 --penalty-strength 0.25 --maximization-algorithm random-greedy`)
- Specificity: query up to 4 mismatches counting G-U pairs as matches, calling a guide nonspecific if it hits ≥ 1% of sequences in another taxon (`--id-method shard --id-m 4 --id- frac 0.01`)
- Objective function and search: weights *λ_A_* = 0.5 and *λ_L_* = 0.25 in the objective function (defined in Supplementary Note 3b) and finding the best 20 design options (`--obj-fn-weights 0.5 0.25 --best-n-targets 20`)

We made some species-specific adjustments. For influenza A and dengue viruses, two especially diverse species, we decreased the number of tolerated primer mismatches to 2 and allowed at most 5 primers at a site (`-pm 2 --max-primers-at-site 5`); while these further constrain the design, they decrease runtime. For Norwalk virus and Rhinovirus C, we relaxed the number of primers at a site and the maximum region length to identify designs (`--max-primers-at-site 20 --maxtarget-length 500`). For Cervid alphaherpesvirus 2, which has a short genome, we changed the GC-content bounds on primers to be 20%–80% (`--primer-gc-content-bounds 0.2 0.8`) to allow more potential amplicons. For 42 species, we relaxed specificity constraints to identify designs (list and details in code).

Of the 1,933 species, 7 could not produce a design while maximizing activity and enforcing specificity, even with species-specific adjustments. They are: Bat mastadenovirus, Bovine associated cyclovirus 1, Chiropteran bocaparvovirus 4, Cyclovirus PKgoat21/PAK/2009, Finkel-Biskis-Jinkins murine sarcoma virus, Panine gammaherpesvirus 1, and Squirrel fibroma virus. Each of these 7 species has just one genome sequence and ADAPT could not identify a guide set satisfying specificity constraints; it is possible they are misclassified or have very high genetic similarity to other species. When showing results for this objective, we report on 1,926 species.

In addition to using the above settings, which maximizes activity and enforces specificity, we ran ADAPT with 3 other approaches: We minimized the number of guides while enforcing specificity, requiring that guides be predicted to be highly active (as defined in Evaluating dispersion and generalization) in detecting 98% of sequences. We also ran the objectives to maximize activity and minimize guides without enforcing specificity. 67 of the 1,933 species did not yield a design when minimizing the number of guides and enforcing specificity, owing to the constraints with this objective: ADAPT could not identify a guide set that is predicted to be highly active and achieves the desired coverage and specificity.

For species with segmented genomes, we ran ADAPT and produced designs separately for each segment. We then selected the segment whose highest-ranked design option has the best objective value (if multiple clusters, according to the largest cluster). We expect the selected segment to generally be the most conserved one.

In all analyses showing results of the designs (e.g., number of guides, guide activity, and target length), we used the highest-ranked design option output by ADAPT. For the species with more than one cluster, we report the mean across clusters from the highest-ranked design option in each cluster.

For producing designs across vertebrate-infecting viral species, we ran ADAPT on Amazon Web Services using the `x1.16xlarge` instance type. We ran ADAPT in parallel across multiple species to fully utilize the instance’s resources. We evaluated ADAPT’s computational requirements, namely the species-specific runtime and memory usage, as part of these runs on that instance type.

### Designs for evaluating sensitivity and specificity

#### ADAPT design parameters

To generate designs with ADAPT for experimental testing, we used the following arguments unless otherwise noted:

- Initial clustering: force a single cluster (`--cluster-threshold 1.0`)
- Primers and amplicons: primer length of 30, primers must have GC content between 35% and 65%, at most 5 primers at a site, up to 1 mismatch between primers and target sequence for hybridization, primers must hybridize to ≥ 98% of sequences, and length of a targeted genome region (amplicon) must be ≤ 250 nt (`-pl 30 --primer-gc-content-bounds 0.35 0.65 --max-primers-at-site 5 -pm 1 -pp 0.98 --max-target-length 250`)
- Guides: Cas13a guide length of 28 nt, together with our Cas13a predictive model (`-gl 28 --predict-activity-model-path models/classify/model-51373185models/regress/model-f8b6fd5d`)
- Guide activity objective: soft constraint of 1 guide, hard constraint of 5 guides, guide penalty (*λ*) of 0.25, using the randomized greedy algorithm (`--obj maximize-activity --soft-guide-constraint 1 --hard-guide-constraint 5 --penalty-strength 0.25 --maximization-algorithm random-greedy`)
- Specificity: query up to 4 mismatches counting G-U pairs as matches, calling a guide nonspecific if it hits ≥ 1% of sequences in another taxon (`--id-method shard --id-m 4 --id- frac 0.01`)
- Objective function and search: weights *λ_A_* = 0.5 and *λ_L_* = 0.25 in the objective function (defined in Supplementary Note 3b) (`--obj-fn-weights 0.5 0.25`)

For SARS-CoV-2 input sequences, we used the 9,054 complete genomes available on GISAID^58^ as of April 28, 2020. We also used genomes from GISAID for pangolin SARS-like CoV input sequences (isolates from Guangxi, China and Guandong, China). For all other input sequences—SARS-like CoV isolates RaTG13, ZC45, and ZXC21; other SARS-like CoVs; SARS-CoV-1 (also referred to as SARS-CoV); and other Coronaviridae species—we used all genome neighbors from NCBI from each species^31^.

#### Generating test target sequences

Experimentally testing design options output by ADAPT also requires generating representative target sequences. We found representative sequences for a design option, using a collection of genomes spanning diversity of a taxon, as follows: (1) We extracted the amplicon (according to provided positions, e.g., from primer sequences), while extending outward to achieve a minimum length (usually 500 nt). (2) We removed sequences that are too short, e.g., owing to gaps in the alignment. (3) We computed pairwise Mash distances^78^ and performed hierarchical clustering (average linkage) to achieve a desired number of clusters or a maximum inter-cluster distance. (4)

To avoid outliers, we greedily selected (in order of descending size) clusters that include a desired total fraction of sequences, a particular number of targets, or ones representing particular taxa (specifics below). (5) We computed the medoid of each cluster—i.e., the sequence with minimal total distance to all other sequences in the cluster. (6) We used the medoids of each clusters as representative target sequences. The `pick_test_targets.py` program in ADAPT implements the procedure and we used this program.

#### Baseline distribution of activity

We established a baseline distribution of activity using Cas13a guides, to detect SARS-CoV-2, selected from the genomic regions targeted by the United States Centers for Disease Control and Prevention (US CDC) RT-qPCR assays^79^. In particular, we picked 10 random 28-mers from the US CDC N1 amplicon that have a non-G PFS and used these as Cas13a guides, according to the ‘`hCoV-19/Wuhan/IVDC-HB-01/2019`’ genome^58^. We also chose another Cas13a guide at the site of the TaqMan probe with a non-G PFS. We did the same from the US CDC N2 amplicon. In addition, in the N1 and N2 amplicons, we used ADAPT to design a single guide with maximal activity (ignoring specificity) from within the amplicon. This provides 24 guides in total.

#### Experimental designs with ADAPT

To evaluate the activity and lineage-level specificity of SARS-CoV-2 designs, we used ADAPT to produce 10 design options for detecting SARS-CoV-2. We increased the specificity in ADAPT to call a guide non-specific if it hits any sequence outside SARS-CoV-2 and also use the greedy maximization to obtain more intuitive outputs because, in this case, we expect only a single Cas13a guide for each design option (`--id-frac 0 --maximization-algorithm greedy`). We enforced specificity to not detect any sequences outside of SARS-CoV-2 from the SARS-related CoV species (including related bat and pangolin coronavirus isolates) and also to not detect sequences from the other 43 species in the Coronaviridae family. Owing to experimental constraints, we tested the highest-ranked 5. We generated targets for each design option against which to test, using the ones representative of SARS-CoV-2; pangolin SARS-like CoVs (isolates from Guangxi, China); bat SARS-like CoV isolates ZC45 and RaTG13; and SARS-CoV-1.

To further evaluate activity and subspecies-comprehensiveness, we used ADAPT to produce 10 design options for detecting the SARS-CoV-2–related taxon. In referring to SARS-CoV-2–related, we use the definition given in Fig. 1b of ref. 48; it encompasses SARS-CoV-2 and several related bat and pangolin SARS-like coronaviruses. To correct for sampling biases, we used 10 sampled SARS-CoV-2 genomes as input so that they make up roughly half of sequences in the SARS-CoV-2–related taxon. We used the same adjusted arguments in ADAPT as used for the SARS-CoV-2 designs (`--id-frac 0 --maximization-algorithm greedy`). We enforced specificity to not detect any sequences outside of SARS-CoV-2–related from the SARS-related CoV species (including other bat SARS-like coronaviruses) and also to not detect sequences from the other 43 species in the Coronaviridae family. For each design option, we generated targets, and used the ones representative of SARS-CoV-2; pangolin SARS-like CoVs (isolates from Guangxi, China and Guangdong, China); bat SARS-like CoV isolates ZC45, ZXC21, and RaTG13; and SARS-CoV-1. For this experiment, the SARS-CoV-1 target allows us to evaluate specificity, while the others allow us to evaluate activity and subspecies-comprehensiveness.

We used ADAPT to produce 10 design options to detect the SARS-related CoV species, and we used these to evaluate activity, species-comprehensiveness, and specificity. To correct for sampling biases, we used 300 sampled SARS-CoV-2 genomes as input so that they make up roughly half of sequences in the species. We enforced specificity to not detect sequences from the other 43 species in the Coronaviridae family. For each design option, we generated representative targets that encompass SARS-CoV-2, SARS-CoV-1, bat SARS-like CoVs, pangolin SARS-like CoVs, MERS-CoV, Human coronavirus OC43, and Human coronavirus HKU1. There were 8 or 9 representative targets in total for each design option.

To evaluate species-comprehensiveness, we focused on Enterovirus B and used ADAPT to produce 10 design options. Owing to its extensive diversity, we made several adjustments to arguments, which help to increase the space of potential design options (`--primer-gc-content-bounds 0.30 0.70 -pm 4 -pp 0.80 --max-primers-at-site 10 --id-frac 0.10 --penalty-strength 0.15`). We enforced specificity to not detect the 18 other species in the Enterovirus genus. For each design option, we generated representative targets from clusters that encompass at least 90% of all sequences. There were between 1 and 15 targets for each design option (the precise number depends heavily on the location of the design option amplicon in the genome). We additionally tested specificity within the Enterovirus genus by generating a single representative target for each of Enterovirus A, Enterovirus C, and Enterovirus D.

To benchmark ADAPT’s designs for Enterovirus B, we created baseline Cas13a guides using an entropy-based approach that identifies conserved sites. For each of ADAPT’s design options, we considered the amplicon it targets. Then, we computed the information-theoretic (Shannon) entropy, over alleles, at every site in the amplicon. (We counted an ambiguous base fractionally and a gap as a “base”.) We define the average entropy of a 28 nt site to be the mean entropy across its 28 positions. The approach finds the site in the amplicon that has the minimal average entropy and an active (`non-G`) PFS in GenBank accession MK800120. Our entropy-based baseline guide is the sequence from GenBank accession MK800120 at this site. We performed this process in the amplicon from each of ADAPT’s designs to generate and test one baseline guide; for 5 of the 10 designs, we generated and tested two baseline guides, where the second was from the site with the second lowest entropy and an active PFS. The approach is implemented in ADAPT’s `design_naively.py` program.

We built a positive control into each target. In particular, we added the sequence 5’-CACTATAGGGGCTCTAGCGACTTCTTTAAATAGTGGCTTAAAATAAC-3’ to the 5’ end of each target and included in our tests of every target a guide with protospacer sequence 5’-GCTCTAGCGACTTCTTTAAATAGTGGCT-3’.

### Experiments evaluating sensitivity and specificity

#### Experimental procedure

We largely followed the CARMEN-Cas13 platform^8^ for experimentally validating ADAPT’s designs, with some key differences. DNA targets were ordered from Integrated DNA Technologies and in vitro transcribed using the HiScribe T7 High Yield RNA Synthesis Kit (New England Biolabs). Transcriptions were performed according to the manufacturer’s recommendations with a reaction volume of 20 μL that was incubated overnight at 37 °C. The transcribed RNA products were purified using RNAClean XP beads (Beckman Coulter) and quantified using NanoDrop One (Thermo Scientific). The RNA was serially diluted from 10^11^ to 10^4^ cp/μL and used as input into the detection reaction. crRNAs were synthesized by Integrated DNA Technologies, resuspended in nuclease-free water, and diluted to 1 μM for input into the detection reaction. The Cas13 detection reactions were made into two separate mixes for loading onto a 192.24 Dynamic Array IFC for Gene Expression (Fluidigm). The assay mix contained 42.5nM LwaCas13a, 42.5nM crRNA, 2× Assay Loading Reagent (Fluidigm), and nuclease-free water. The sample mix contained 1 μL RNAse Inhibitor (New England Biolabs), 1× ROX Reference Dye (Invitrogen), 1× GE Sample Loading Reagent (Fluidigm), 1.95 nM quenched synthetic fluorescent RNA reporter (`FAM/rUrUrUrUrUrUrU/3IABkFQ/`, Integrated DNA Technologies), 9nM MgCl_2_ in a nuclease assay buffer (40mM Tris-HCl, 1 mM DTT pH 7.5). Syringe, Actuation Fluid, Pressure Fluid (Fluidigm), and 4μL of assay and sample mixtures were loaded into their respective locations on a 192.24 IFC according to the manufacturer’s instructions. The IFC was loaded onto the IFC Controller RX (Fluidigm) where the ‘Load Mix’ script was run. After proper IFC loading, images over a two-hour period were collected using a custom protocol on Fluidigm’s Biomark HD.

#### Displaying experimental results

We plotted reference-normalized background-subtracted fluorescence for guide-target pairs. For a guide-target pair (at some time point *t* and target concentration), we first computed the reference-normalized value as

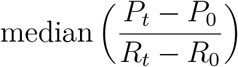

where *P_t_* is the guide signal (FAM) at the time point, *P*_0_ is its background measurement before the reaction, *R_t_* is the reference signal (ROX) at the time point, *R*_0_ is its background measurement, and the median is taken across Fluidigm’s replicates. We performed the same calculation for the no-template (water) control of the guide, providing a background fluorescence value for the guide at *t* (when there were multiple technical replicates of such controls, we took the mean value across them). The reference-normalized background-subtracted fluorescence for a guide-target pair is the difference between these two values. Note that, by definition, plotted values greater than 0 represent fluorescence that exceeds background and the no-template control (‘NC’ in figures) has value of 0. When plotting the no-template control separately (Supplementary Fig. 31), we show reference-normalized values without background-subtracting. In heatmaps showing fluorescence at a fixed time point, we used the middle time point (59 minutes). In kinetic curves that show fluorescence over time (for example, Fig. 5b), we smoothed the value by taking the rolling mean within a window of 2 time points.

When displaying the top-ranked design options from ADAPT (for example, in Fig. 5d–h), we ordered them according to the predicted activity of the Cas13a guides in expectation across the input genomes. ADAPT’s ranking incorporates additional factors (Supplementary Note 3b) that reflect amplification potential, and we used ADAPT’s objective function to identify the top *N* design options to test. But we ordered them according to only predicted fluorescent activity because our experimental testing did not involve amplification. When plotting fluorescence for a design that uses more than one guide, we plot the maximum fluorescence across the guides (computed separately at each target, target concentration, and measurement time point). This is analogous to ADAPT’s model for measuring a probe set’s activity (Supplementary Note 1a), in which its activity in detecting a target sequence equals that of the best probe in the set for detecting that sequence.

#### Evaluating specificity against non-viral taxa

We performed an in silico comparison of all experimentally-tested guides with human transcripts and bacterial pathogens to determine if there is potential cross-reactivity. We first built an index consisting of human transcript sequences from GENCODE v38^80^ and NCBI reference genome sequences for 11 bacterial pathogens (Bordetella pertussis (NC_018518.1); Chlamydia pneumoniae ([NC_-005043.1](https://www.ncbi.nlm.nih.gov/nuccore/NC_005043.1)); Haemophilus influenzae (NZ_CP009610.1); Legionella pneumophila (NZ_CP013742.1); Mycobacterium tuberculosis (NC_000962.3); Mycoplasma pneumoniae (NZ_CP010546.1); Pseudomonas aeruginosa (NC_002516.2); Staphylococcus epidermidis (NZ_CP035288.1); Streptococcus pneumoniae (NZ_CP046357.1); Streptococcus pyogenes (NZ_CP010450.1); Streptococcus salivarius (NZ_CP066093.1)). We also included, as positive controls for the analysis, NCBI reference sequence genomes for SARS-CoV-1 (NC_004718.3) and SARS-CoV-2 (NC_045512.2).

We sought to query guide sequences against this index while tolerating multiple mismatches over a short query length (i.e., the guide length of 28 nt). To enable this, we used Bowtie 281 to align guide sequences to the index with the parameters `-a --end-to-end -N 1 -L 7 -i S,1,1 --ma 0 -- mp 1,1 --rdg 100,1 --rfg 100,1 --score-min L,-4,0`. These settings permit us to identify all alignments of guides, against our index, having 4 or fewer mismatches across the length of the guide without tolerating gaps. Such alignments represent potential non-specificity of the guides.

## Data availability

Data is available in several repositories:

- The CRISPR-Cas13a library and activity dataset is available at: https://github.com/broadinstitute/adapt-seq-design/tree/82db28/data
- Serialized trained models are available at: https://github.com/broadinstitute/adapt-seq-design/tree/82db28/models/cas13
- Experimentally tested designs are available at: https://github.com/broadinstitute/adapt-designs/tree/783ac9

## Code availability

Code is available in several repositories :

- ADAPT is freely available under the MIT license at: https://github.com/broadinstitute/adapt
- Code to replicate the predictive modeling and analyses is available at: https://github.com/broadinstitute/adapt-seq-design
- Code to replicate the designs across the vertebrate-infecting viral species is available at: https://github.com/broadinstitute/adapt-designs-continuous
- Code to replicate the other analyses in this paper is available at: https://github.com/broadinstitute/adapt-analysis

## Supplementary Note 1

This note describes two formulations for objective functions implemented in ADAPT. Unless otherwise noted, for designs and analyses in the paper, we use the formulation in Design formulation #1: maximizing expected activity.

### 1a Design formulation #1: maximizing expected activity

#### Objective

Let *S* be an alignment of sequences from species *t* in a genomic region. We wish to find a set *P* of probes that maximizes detection activity over these sequences. This objective bears some similarity to the problem of designing PCR primers that cover a maximum number of sequences^82^, though solutions to that problem binarize decisions about detection rather than accommodate continuous predictions. As described in Methods, we have a model to predict a measurement of detection activity between one probe *p* and one sequence *s* ∈ *S*, which we represent by *d*(*p, s*). (In Fig. 3a, we use *A*(*P, s*) to represent this quantity.) While more than one probe in the set *P* may be able to detect *s*, we use the best probe against *s* to measure *P*’s detection activity for *s*; that is, the predicted detection activity is

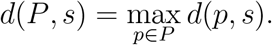

One way to consider why taking the maximum is reasonable is that, in practice, we could apply each *p* ∈ *P* in parallel reactions to a sample even if only one *p* works well for the particular target in that sample. We also define *d*(*P, s*) when *P* is the empty set to be the lowest value in the range of *d*(*p, s*), indicating no detection (in practice, 0).

We represent the predicted detection activity by *P*, against all sequences in *S*, with the function *F*(*P*). *F*(*P*) is the expected value of *d*(*P, s*) taken over all the *s* ∈ *S*; the weights *w_s_* for each *s* can reflect a prior probability on applying the detection in practice to targeting a sequence like *s* (e.g., based on the genome’s date or geographic location). Thus, we define

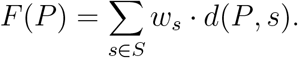

We currently set a uniform prior over targeting the genomes in *S*, with the effect being that all *w_s_* are equal.

We must introduce penalties for the number of probes. Striving for a small number of probes is important because (a) if used in separate reactions, this adds time and labor; (b) if multiplexed in one reaction, there is generally competition in binding to a target, and having more in a reaction may reduce the resulting detection signal; and (c) they require time and money to synthesize, and to experimentally validate. For this penalty, we use a soft constraint *m_p_* and hard constraint 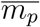 on the number of probes, with 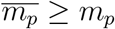. We wish to solve

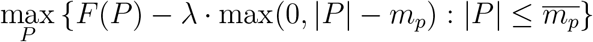

where *λ* > 0 serves as a weight on the penalty. *λ* reflects a tolerance for higher *F*(*P*) at the expense of more probes.

#### Submodularity of the objective

Let 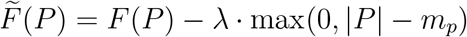. We want to prove that 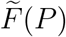 is submodular.

We start by showing first that *d*(*P, s*) is submodular. For ease of notation, we drop *s*, referring to *d*(*P, s*) as *d*(*P*) and *d*(*p, s*) as *d*(*p*). Consider probe sets *A* and *B*, with *A* ⊆ *B,* and some possible probe *x* ∉ *B*. Note that

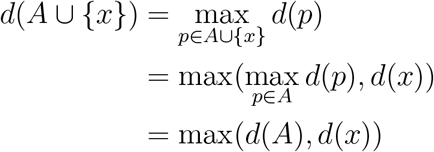

and therefore *d*(*A* ∪ {*x*}) – *d*(*A*) ≥ 0. Likewise, *d*(*B* ∪ {*x*}) = max(*d*(*B*), *d*(*x*)), and *d*(*B* ∪ {*x*}) – *d*(*B*) ≥ 0. Also, since *A* ⊆ *B*, we have

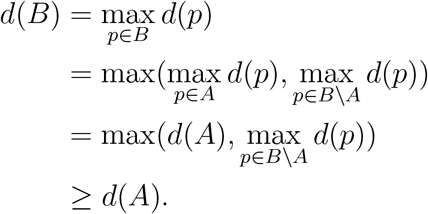

We now consider two cases:

- Assume *d*(*B*) ≥ *d*(*x*). Then, *d*(*B* ∪ {*x*}) – *d*(*B*) = max(*d*(*B*), *d*(*x*)) – *d*(*B*) = 0. Therefore, *d*(*A* ∪ {*x*}) – *d*(*A*) ≥ *d*(*B* ∪ {*x*}) – *d*(*B*).
- Assume *d*(*B*) < *d*(*x*). It follows from *d*(*B*) ≥ *d*(*A*) that *d*(*x*) > *d*(*A*) and that *d*(*x*) – *d*(*A*) ≥ *d*(*x*) – *d*(*B*). Since max(*d*(*A*), *d*(*x*)) = *d*(*x*) and max(*d*(*B*), *d*(*x*)) = *d*(*x*), we have max(*d*(*A*), *d*(*x*)) – *d*(*A*) ≥ max(*d*(*B*), *d*(*x*)) – *d*(*B*). Therefore, in this case as well, *d*(*A* ∪ {*x*}) – *d*(*A*) ≥ *d*(*B* ∪ {*x*}) – *d*(*B*).

Hence, *d*(*P, s*) is submodular. Since *F*(*P*) is a non-negative linear combination of *d*(*P, s*), it follows that *F*(*P*) is submodular.

Using the above result, we show that 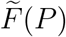 is submodular. Again, consider probe sets *A* and *B*, with *A* ⊆ *B*, and some possible probe *x* ∉ *B*. We want to show that 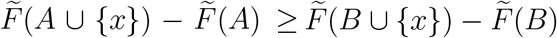. We have that

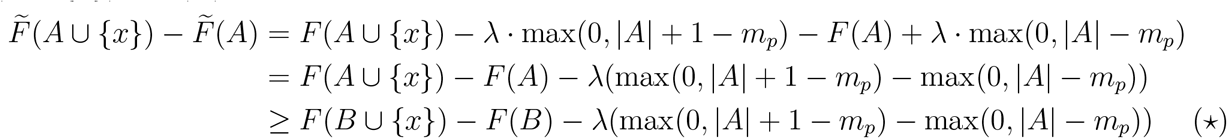

where the last step follows from submodularity of *F*(*P*). We now consider two cases:

- Assume |*A*| ≥ *m_p_*. Since *A* ⊆ *B*, |*B*| ≥ *m_p_*. Continuing from (*), we have

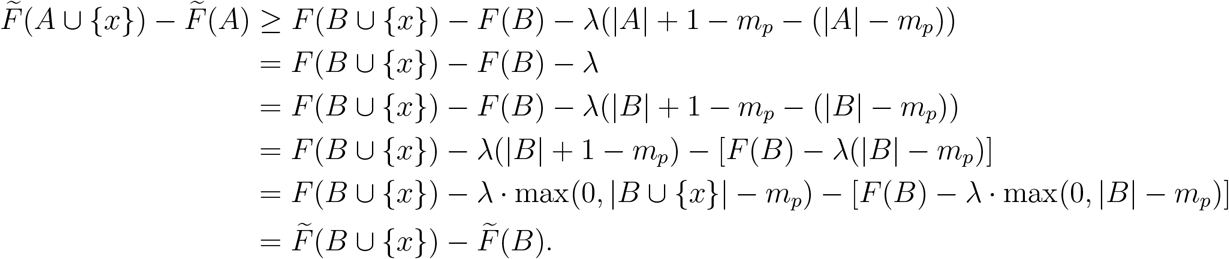
- Assume |*A*| < *m_p_*. Continuing from (*) in this case, we now have

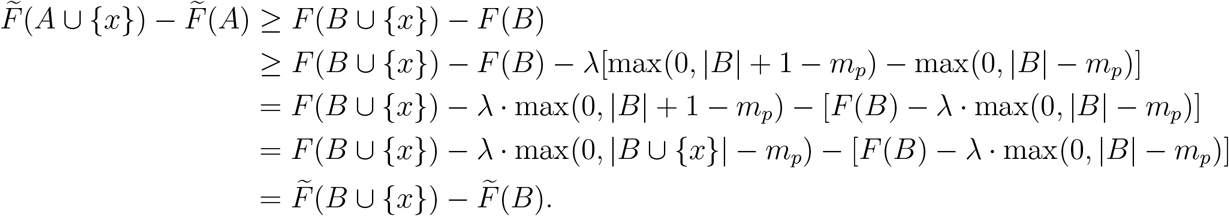

Hence, 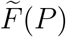 is submodular.

Note that 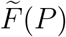 is non-monotone owing to the penalty term.

#### Non-negativity of the objective

Now we show how to ensure that 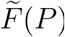 is non-negative. Let *P* only contain probes *p* such that 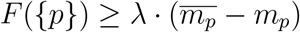. Since *F* is monotonically increasing, 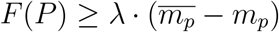. Thus,

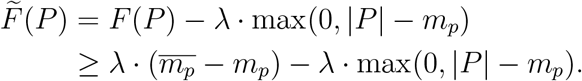

If |*P*| ≤ *m_p_*, then

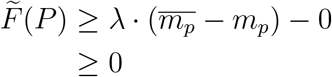

where the last inequality follows from 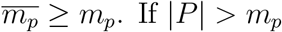. If |*P*| > *m_p_*, then

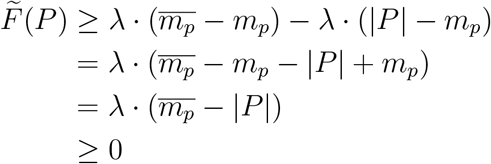

where the last inequality follows from 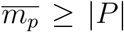. If *P* is the empty set, 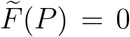 according to our definition of *d*(*P, s*). Therefore, 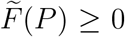 always. Let our ground set *Q* be the set of probes from which we select *P*—i.e., *P* ⊆ *Q*. To enforce non-negativity, we restrict *Q* to only contain probes *p* such that 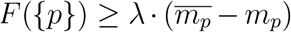. In other words, every probe has to be sufficiently good. In practice, given our activity function, *λ* ∈ [0.1, 0.5] is a reasonable choice and the constraint on *F*({*p*}) is generally met; for example, with *λ* = 0.25, 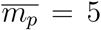, and *m_p_* = 1, then we require *F*({*p*}) ≥ 1.

#### Solving for *P*

Recall we want to solve

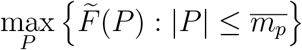

where 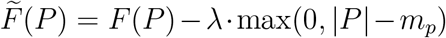. We need to maximize a non-negative and non-monotone submodular function subject to a cardinality constraint. The classical discrete greedy algorithm^43^ may provide poor results, and would not offer theoretical guarantees, because it assumes a monotone function. We apply the recently-developed discrete randomized greedy algorithms in ref. 42, namely Algorithm 1, which provides a 1/*e*-approximation for non-monotone functions. (Algorithm 5, which provides a better approximation ratio, is likely to not be much better in our case because the constraint 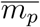 is small compared to the size of the ground set.)

**Algorithm 1.**
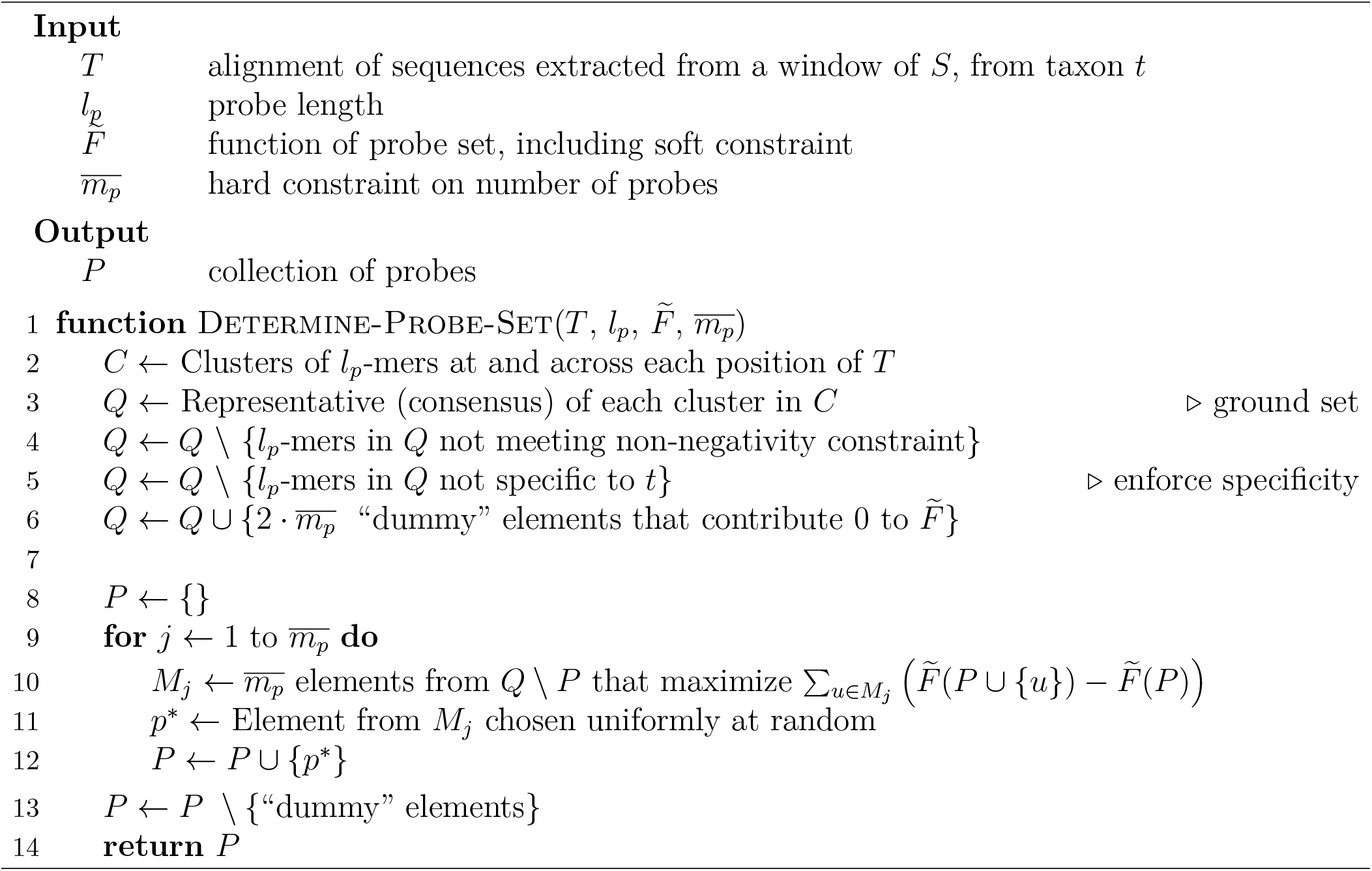
Construct set of probes *P* to maximize 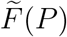 subject to hard constraint.

Based on the work in ref. 42, the function Determine-Probe-Set (Algorithm 1) shows how we compute *P* to detect a particular genomic window of an alignment *S*. We use locality-sensitive hashing to rapidly cluster potential probe sequences^2^ throughout the window, and their representatives form the ground set *Q* of probes (line 3). Then, we require that probes in *Q* be specific to the taxon to which *S* belongs, using the methods in Supplementary Note 2d). We add to the ground set “dummy” elements that provide a marginal contribution of 0 to any set input to 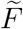 (line 6), as required by an assumption of the algorithm (Reduction 1 in ref. 42). Then, we greedily choose 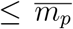 probes, at each iteration selecting one randomly from a set of not-yet-chosen probes that maximize marginal contributions to 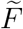 (lines 10–11).

The runtime to design probes is practical in the typical case. Here we ignore the runtime of evaluating specificity (line 5), which is given in Supplementary Note 2d. Let *L* be the window length and *n* be the number of sequences. There are *O*(*nL*) probes in the ground set in the worst-case, and they take *O*(*nL*) time to construct (line 3). Finding the 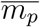 elements that maximize marginal contributions (line 10) takes *O*(*nL*) time, and we do this 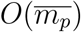 times. Thus, the runtime in the worst-case is 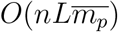. In a typical case, the number of clusters at a position in the window is a small constant (≪ *n*) owing to sequence homology in the alignment; thus, the size of the ground set is *O*(*L*), although it still takes *O*(*nL*) time to construct. Now, finding the 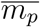 elements that maximize marginal contributions takes *O*(*L*) time, and we do this 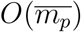 times. So the runtime in a typical case is 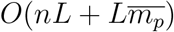. Note that, in general, 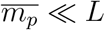 and 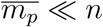.

### 1b Design formulation #2: minimizing the number of probes

#### Objective

As in the above objective, let *S* be an alignment of sequences from species *t* in a genomic region and let *d*(*p, s*) be a predicted detection activity between one probe *p* and one sequence *s* ∈ *S*. We wish to find a set *P* of probes with minimal |*P*| that satisfies constraints on detection activity across these sequences. In particular, we introduce a fixed detection activity *m_d_* and say that *p* is highly active in detecting *s* if *d*(*p, s*) ≥ *m_d_*. To define whether *P* detects a sequence with high activity, let

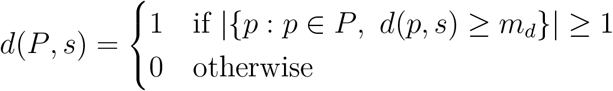

We additionally introduce a lower bound *f_S_* on the minimal fraction of sequences in *S* that must be detected with high activity. Then, we wish solve

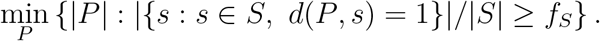

That is, we want to find the smallest probe set that detects, with high predicted activity, at least a fraction *f_S_* of all sequences.

#### Solving for *P*

To approximate the optimal *P*, ADAPT follows the canonical greedy solution to the set cover problem^83,84^ in which the universe consists of the sequences in *S* and each possible probe covers a subset of sequences in *S*. Similar approaches have been used for PCR primer selection^9, 14, 15, 85, 86^; in contrast to prior approaches, rather than starting with a collection of candidate probes (i.e., the sets), we construct them on-the-fly.

Iteratively, we approximate a probe that covers the most number of sequences that still need to be covered. Here, a probe *p covers* a sequence *s* if *d*(*p, s*) ≥ *m_d_.* Find-Optimal-Probe, shown in Algorithm 2, implements a heuristic. Briefly, at each position Find-Optimal-Probe rapidly clusters *l_p_*-mers in the input sequences (*l_p_* is the probe length) by sampling nucleotides—i.e., concatenating locality-sensitive hash functions drawn from a Hamming distance family—and uses each of these clusters to propose a probe. It iterates through the clusters in decreasing order of score, stopping early (line 11) if it is unlikely that remaining clusters will provide a probe that achieves more coverage than the current best. This procedure relies on two subroutines, Score-Cluster and Num-Detect, that are described below.

Using this procedure, it is straightforward to construct a set of probes in the window (region) that achieve the desired coverage by repeatedly calling Find-Optimal-Probe. This is shown by Determine-Probe-Set, in Algorithm 3. In other words, the output probes collectively detect, with high activity, the sequences in the region.

This approach, with on-the-fly construction of probes, is similar to a reduction to an instance of the set cover problem, the solution to which is essentially the best achievable approximation^87, 88^. In such a reduction, each set would represent one of the 4^*lp*^ possible probes, consisting of the sequences that it would detect with high activity. Then, each iteration would identify the probe that detects, with high activity, the most not-yet-covered sequences. Here, rather than starting with such a large space, we use a heuristic to approximate the probe at each iteration.

**Algorithm 2.**
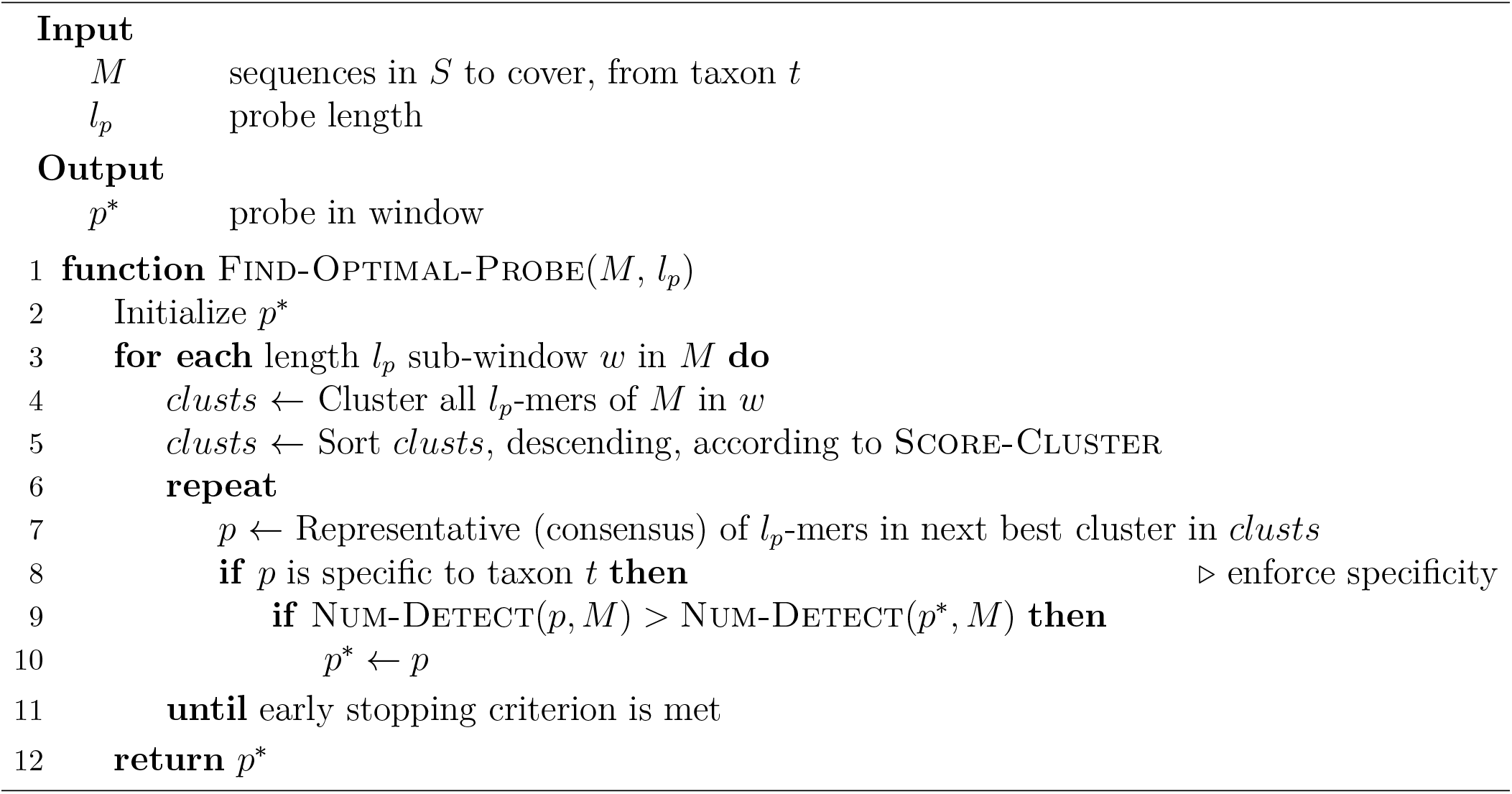
Construct probe *p*\* with highest coverage.

**Algorithm 3.**
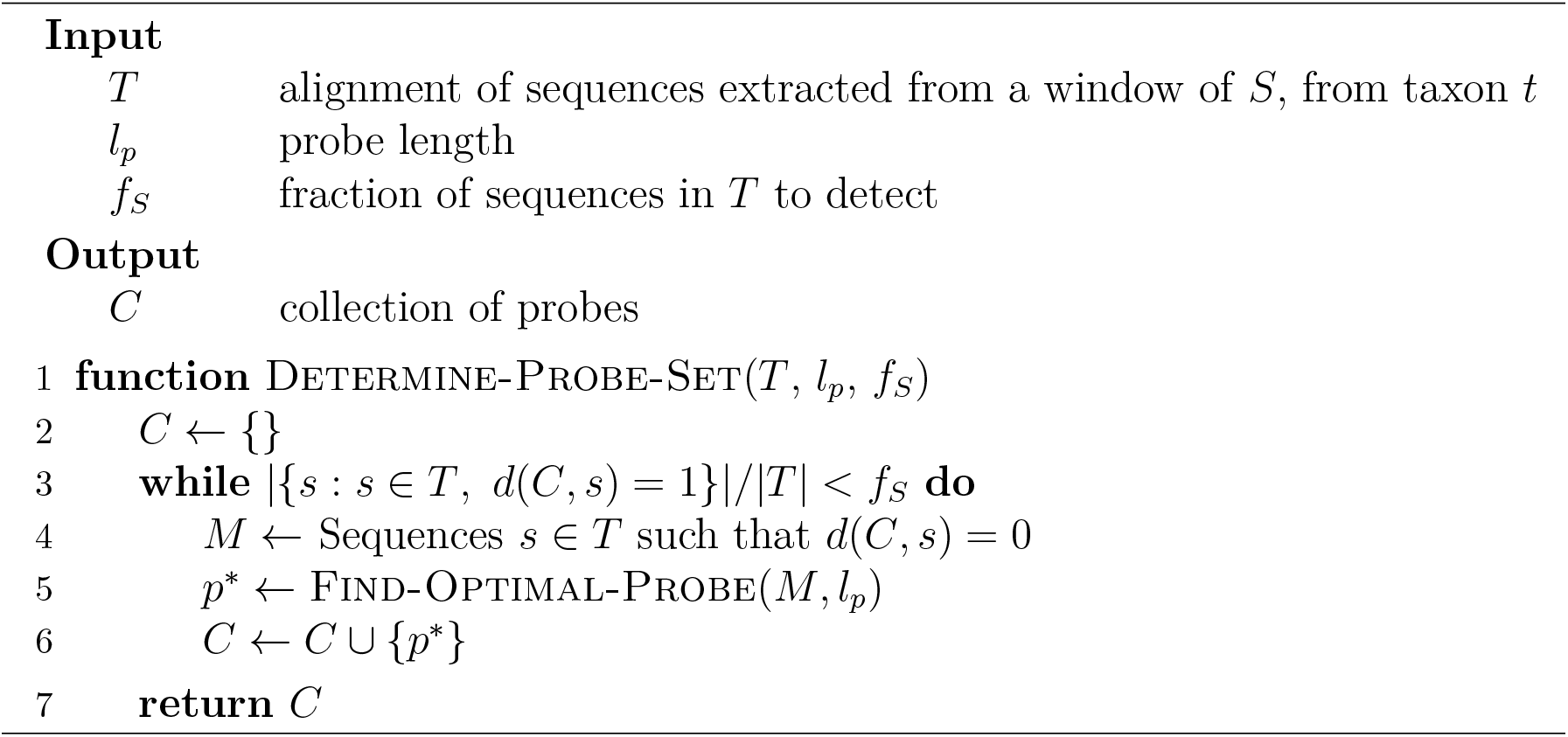
Construct minimal collection of probes in window that collectively achieve desired detection coverage.

The runtime to design probes in a window is poor in the worst-case but practical in the typical case. Let *n* be the number of sequences in the alignment and *L* be length of the window. In the worst-case, we choose *n* different probes in the window. Each choice requires iterating over *O*(*L*) positions, and at each one we iterate through *O*(*n*) clusters, taking *O*(*n*) time to evaluate the probe proposed by each cluster with Num-Detect. Thus, this is *O*(*n*^3^*L*) time. In a typical case, there is a small constant number of clusters owing to sequence homology across the alignment, and the number of probes needed to achieve the constraint is also a small constant. Selecting each probe requires iterating over *O*(*L*) positions, and at each one we consider *O*(1) clusters, taking *O*(*n*) time again to evaluate the probe proposed. So the runtime is *O*(*nL*) with these assumptions.

#### Scoring clusters and detection across sequences

Sequences from *S* can be *grouped* according to metadata such that each group receives a particular desired coverage (*f_S_g__*). For example, in ADAPT’s implementation they can be grouped according to year (each group contains sequences from one year), with a desired coverage that decays for each year going back in time, so that ADAPT weighs more recent sequences more heavily in the design.

There are two subroutines in Algorithm 2 that we consider here: scoring a cluster and computing the number of sequences detected by a probe. These must account for groupings. First, on line 5 of Find-Optimal-Probe, the function Score-Cluster(*clust*) computes the number of sequences *clust* ∈ *clusts* contains that are needed to achieve the desired coverage across all the groups. That is, it calculates

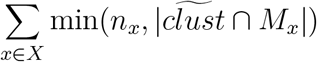

where *X* is the collection of sequence groups, *n_x_* is the number of sequences from group *x* that must still be covered to achieve *x*’s desired coverage, 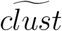 gives the sequences of *M* from which the *l_p_*-mers in *clust* originated, and *M_x_* consists of the sequences in *M* that are in group *x*. In essence, it computes a contribution of each cluster toward achieving the needed coverage of each group, summed over the groups. Similarly, on line 9 of Find-Optimal-Probe, the function Num-Detect(*p, M*) is the detection coverage provided by probe *p* across the groups. In particular, its value is

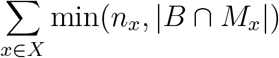

where *B* is the set of sequences in *M* that *p* covers—i.e., *B* = {*s* : *s* ∈ *M, d*(*p, s*) ≥ *m_d_*}.

These subroutines are intuitive in the case where sequences are not grouped. Equivalently, consider a single group *x*_0_. Here, Score-Cluster(*clust*) is 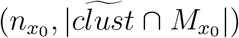. Since 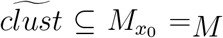, this is 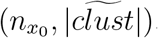. Thus, the score is simply the size of the cluster (larger clusters are preferred), or *n*_*x*_0__ for clusters large enough so as to provide more than sufficient coverage. Similarly, Num-Detect(*p, M*) is min(*n*_*x*0_, |*B* ⋂ *M*_*x*0_|). Because *B* ⊆ *M*_*x*0_ = *M*, this is min(*n*_*x*0_, |*B*|). So Num-Detect is effectively the number of sequences covered by *p* that must still be covered to achieve the coverage constraint.

Furthermore, if sequences are grouped, note that line 3 of Algorithm 3 instead iterates until achieving the desired coverage for each group.

A recent paper^89^ on submodular optimization looks at a similar problem; it refers to the groupings in this problem as ground sets and provides an approximation ratio given by the greedy algorithm.

## Supplementary Note 2

This note describes an overview of the challenge of evaluating specificity and two formulations, implemented in ADAPT, for doing so. Unless otherwise noted, for designs and analyses in the paper, we use the formulation in Exact trie-based search for probe near neighbors.

### 2a Overview

In applications where differentially identifying a taxonomy is important, ADAPT ensures that the probes it constructs are specific to the taxonomy they are designed to detect. In general, the probes directly perform detection; thus, their specificity is ADAPT’s focus, rather than other aspects of a design, such as primers.

The framework for this is as follows. Initially, ADAPT constructs an index of probes across all input taxonomies, which includes the taxonomies and particular sequences containing each probe. This index could also include background sequence to avoid, such as the human transcriptome, although we generally do not include non-viral background sequence. Then, when designing a probe for a taxonomy *t_i_* with genomes *S_i_*, ADAPT queries this index to determine its specificity against all sequences from any *S_j_* for *j* ≠ *i*—namely, to find hits within a specified number of mismatches of the query. The results inform whether the probe might detect some fraction of sequence diversity in *t_j_*. Typically, ADAPT deems a probe to be non-specific if the query yields hits in at least 1% of the sequences from another taxon. ADAPT performs this query while constructing the ground set, as described in Supplementary Note 1.

This problem is computationally challenging. When querying, we generally wish to tolerate a high divergence within a relatively short query to be conservative in finding potential non-specific hits— e.g., up to ~5 mismatches within 28 nt. Also, G-U wobble base pairing (described below) generalizes the usual alphabet of matching nucleotides. Together, these challenges mean that popular existing approaches, including seed/MEM techniques, are not fully adequate for performing queries.

### 2b G-U wobble base pairing

Some detection applications (e.g., CRISPR-Cas13) rely on RNA-RNA binding. That is, the probe we design is synthesized as RNA and the target is RNA as well. RNA-RNA base pairing allows for more pairing possibilities than with DNA-DNA. In particular, G may bind with U, forming a *G-U wobble base pair.* It has similar thermodynamic stability to the usual Watson-Crick base pairs^90^.

In our Cas13a dataset, we find that U-g mismatches (`U` in the target, `G` in the guide RNA spacer) preserves high activity across guide-target pairs (i.e., when 2 of 2 mismatches or 3 of 3 mismatches are U-g, activity is less likely to be reduced), but we do not observe this same effect for G-u mismatches (Supplementary Fig. 16c). Both U-g and G-u wobble pairings might be tolerated for binding, but the resulting geometries of the pairings could affect Cas13a nuclease activation in different ways. Nevertheless, treating both U-g and G-u pairs (collectively, G-U) as comparable to Watson-Crick base pairs would lead to designs at least as specific as if we were to only do so for U-g pairs. It also permits our algorithms to be tolerant of G-u pairs if other applications, with different enzymatic processes, tolerate those pairs well.

In ADAPT, we wish to treat G-U base pairs as matching when querying for a probe’s specificity. For simplicity, here we will use `T` instead of `U` (the RNA nucleobase `U` replaces the DNA nucleobase `T`), and thus we consider G-T base pairing. In particular, we consider a base *g*[*i*] in a probe to match a base *s*[*i*] in a target sequence if either (a) *g*[*i*] = *s*[*i*], (b) *g*[*i*] = `A` and *s*[*i*] = `G`, or (c) *g*[*i*] = `C` and *s*[*i*] = `T`^3^. Note that activity models in ADAPT that are trained for a particular detection technology could prune the query results if the effect is different in some application.

Tolerating G-U base pairing considerably complicates the problem for several reasons. The addition of G-U base pairing raises the probability of a matching hit between a 28-mer and an arbitrary target, thereby expanding the space of potential query results. It also means the Hamming distance between a query and valid hit (considered in the same frame) can often exceed 50% and be as high as 100%. Supplementary Fig. 20 illustrates the challenge in practice on viral genome data.

A similar challenge arises in determining off-target effects when designing small interfering RNA (siRNA)^91, 92^. It is common to ignore the problem (e.g., using BLAST to query for off-targets)^93–96^. Other approaches do address it. One is to treat G-U pairs like a mismatch, albeit not as heavily penalized as a Watson-Crick mismatch^97^; however, with this approach, searching for candidate hits may fail to find valid hits if the Hamming distance between the query and hit is sufficiently high owing to G-U pairs. Another approach uses the seed-and-extend technique where the seed is in a well-defined “seed region” that requires an exact match, tolerating G-U pairs in the seed^98^; although applicable to siRNA, a seed-based approach may fail to generalize if there is no seed region, if it is too short, or if it is not consistent or is tolerant of mismatches. For some RNA interference applications, G-U pairs may be detrimental to the activity of an enzyme complex^99^, and therefore it may not be necessary to fully account for it when determining specificity. None of these approaches are fully satisfying in ADAPT.

To approach the challenge of G-U wobble base pairing, at several points in the algorithms below we use a transformed sequence (Supplementary Fig. 21a). We transform a probe *g* into *g*′ by changing `A` to `G` and changing `C` to `T`; in *g*′, the only bases are `G` and `T`. Likewise, we do this for a target sequence *s*. This is useful because any G-T matching between *s* and the complement of *g* is not reflected by different letters between *g*′ and *s*′—i.e., if the reverse complement of *g* (what we synthesize) matches with *s* up to G-U base pairing, then *g*′ and *s*′ are equal strings.

### 2c Probabilistic search for probe near neighbors

To permit queries for specificity, we first experimented with performing an approximate near neighbor lookup similar to the description in ref. 100 for points under the Hamming distance. Here, we wish to find probes that are ≤ *m* mismatches from a query.

The approach precomputes a data structure *H* = {*H*_1_, *H*_2_,…, *H_L_*} where each *H*_i_ is a hash table that has a corresponding locality-sensitive hash function *h_i_*, which samples *b* positions of a probe. The *h_i_*s bear similarity to the concept of spaced seeds^101^. It chooses *L* to achieve a desired reporting probability *r*:

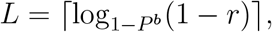

where *P^b^* = (1 – *m/k*)^*b*^ is a lower bound on the probability of collision (for a single *h_i_*) for nearby probes of length *k*. In ADAPT, we have used *r* = 0.95 and *b* = 22. For all probes *g* across all sequences in all taxa *t_j_*, each *H_i_*[*h_i_*(*g*′)] stores {(*g, j*)} where *j* is an identifier of a taxon from which *g* arises and *g*′ is *g* in the two-letter alphabet described above. Additionally, the data structure holds a hash table *G* where *G*[(*g, j*)] stores identifiers of the sequences in *j* that contain *g*. From these data structures, queries are straightforward. For a probe *q* to query, the query algorithm looks up *q*′ in each *H_i_* and check if *q* detects (is within *m* mismatches) each resulting *g*. For the ones that it does detect, *G* provides the fraction of sequences in each taxon containing *g* and therefore provides the fraction of sequences in each taxon that *q* detects. The algorithm deems *q* specific iff this fraction is sufficiently small. Note that, when designing probes for a taxon *t_j_*, it is straightforward to mask *j* from each *H_i_*; this is important for query runtime because most near neighbors would be from *j*.

This approach would be suitable if we were to not have to consider G-U base pairing, but this consideration makes it too slow for many applications. To accommodate G-U base pairs, it stores two-letter transformed probes (*g*′) and likewise queries transformed probes (*q*′). The dimensionality reduction enables finding hits within ≤ *m* mismatches of a query *q*, sensitive to G-U base pairs, but it also means that most results in each *H_i_*[*h_i_*(*q*′)] are far from *q*. As a result, the algorithm spends most of its time validating each of these results by comparing it to *q*. A higher choice of *b* can counteract this issue, but results in higher *L* and thus requires more memory. Also, the approach is probabilistic and may fail to detect non-specificity; while the reporting probability might be high per-taxon, if we use ADAPT to design across many taxa it becomes more likely to output a non-specific assay. Thus, below, we develop an alternative approach that is more tailored to the particular challenges we face.

### 2d Exact trie-based search for probe near neighbors

Here we describe a data structure and query algorithm that permits fully accurate queries for non-specific hits of a probe. Unlike the probabilistic approach above, this will always detect nonspecificity if present, and we show it is fast compared to a baseline. Having one trie containing all the indexed probes would satisfy the goal of being fully accurate because we could branch, during a query, for mismatches and G-U base pairs; however, the extensive branching involved means that query time would depend on the size of the trie and may be slow. To alleviate this, we place (or *shard*) the probes across many smaller tries.

Briefly, the data structure stores an index of all probes across the input sequences from all taxa. Let *k* be the probe length (e.g., 28). The data structure splits each probe into *p* partitions (without loss of generality, assume *p* divides *k*). Each partition maps to a 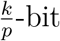 *signature* such that any two matching strings map to the same signature, tolerating G-U base pairing; each bit corresponds to a letter from the two-letter alphabet described in G-U wobble base pairing. There are *p* · 2^*k/p*^ tries in total, each associated with a signature and a partition, and every probe is inserted into *p* tries according to the signatures of its *p* partitions.

To query a probe *q*, the algorithm relies on the pigeonhole principle: tolerating up to *m* mismatches across all of *q*, there will be at least one partition with ≤ [*m/p*] mismatches against each valid hit. For each partition of *q*, the query algorithm produces all combinations of signatures within [*m/p*] mismatches—there are 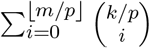 of them—and looks up *q* in the tries with these signatures for the partition. During each lookup, it branches to accommodate G-U base pairing and up to *m* mismatches. Note that the bit signature is sensitive to G-U base pairing—i.e., two positions have the same bit if they might be a match, including owing to G-U pairing—so the algorithm finds all hits, even if the query and hit strings diverge due to G-U pairing.

Supplementary Fig. 21 provides a visual depiction of building the data structure and performing queries, and Algorithms 4 and 5 provide pseudocode.

**Algorithm 4.**
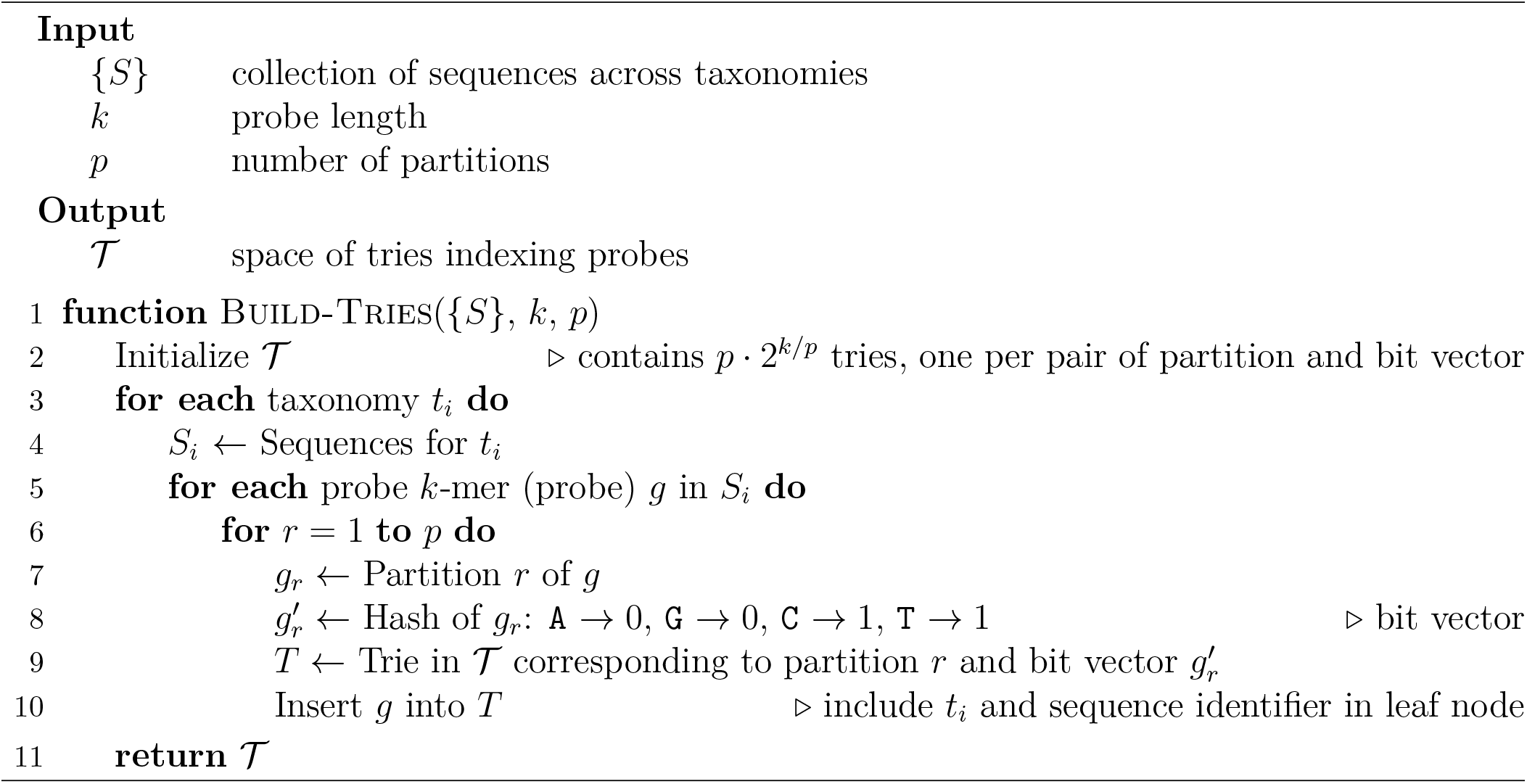
Build data structure of tries to support specificity queries.

**Algorithm 5.**
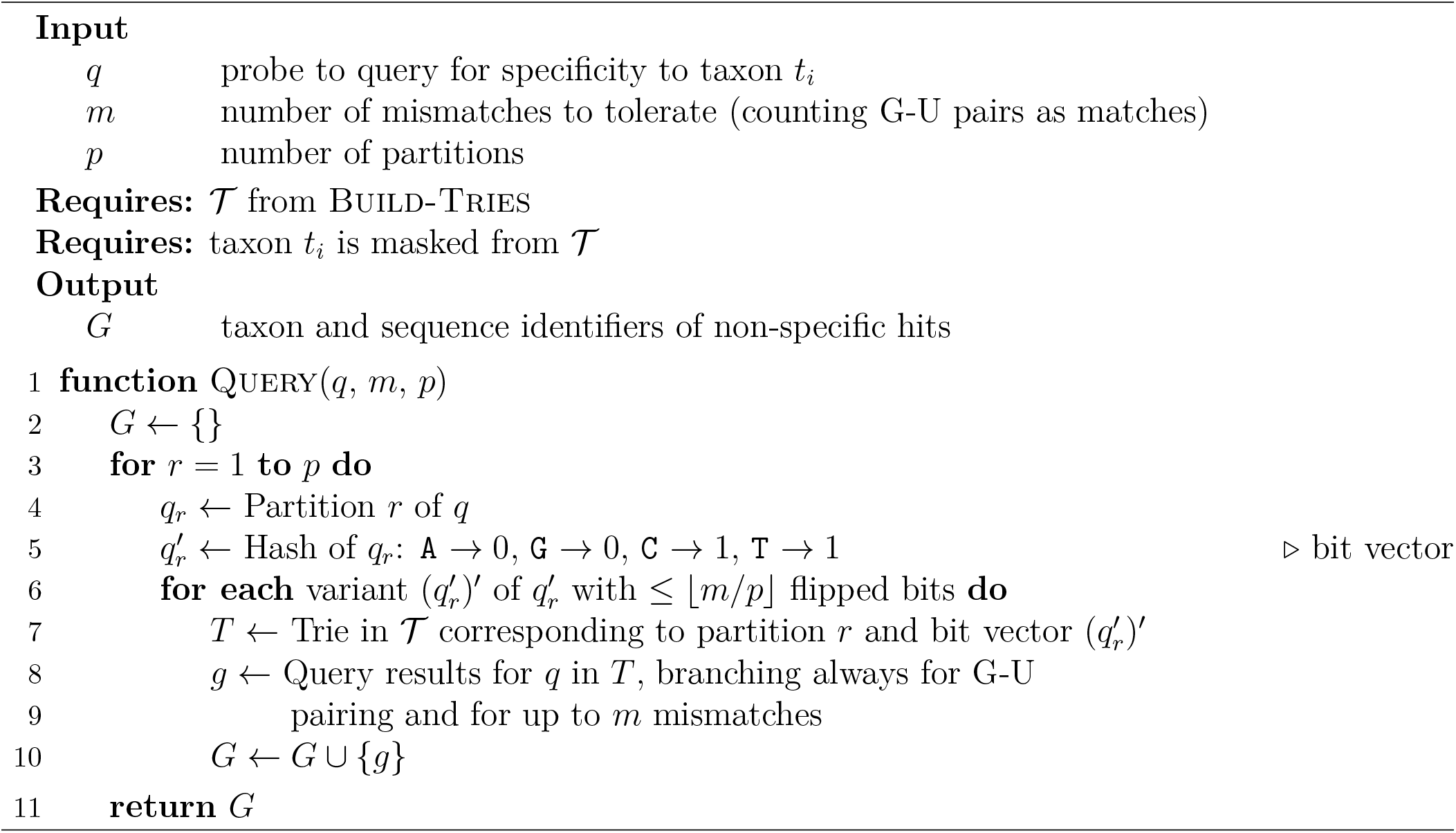
Query tries to find non-specific hits.

A loose bound on the runtime of a query is

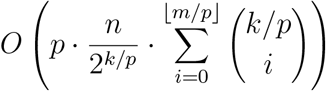

where *n* be the total number of probes indexed in the data structure. The query algorithm performs a search for *p* partitions of a query *q.* For each partition, it considers 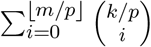 tries, one for each combination of [*m/p*] bit flips. The size of each trie is a loose upper bound on the query time within it; assuming uniform sharding, the size of each is 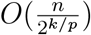. Multiplying the size of each trie by the number of them considered during a query provides the stated runtime. Adjusting *p*, a small constant, allows us to tune the runtime: higher choices reduce the number of bit flips, and thus the number of tries to search, but yield larger tries, and thus requires more time searching within each of them. The runtime does not scale well with our choice of *m*, but this is generally a small constant (up to ~5). The term providing the worst-case query time within each trie, 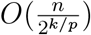, is likely to be a considerable overestimate in practice because queries usually do not need to fully explore a trie.

Because the data structure stores each probe in *p* separate tries, the required memory is *O*(*np*). Although this scales reasonably with *n*, it involves large constant factors and is memory-intensive in practice; one future direction is to compress the tries.

## Supplementary Note 3

This note describes how we link methods, from Supplementary Notes 1 and 2, in ADAPT to form an end-to-end system for designing assays. In particular, this involves identifying amplification primers, searching across genomic regions, and connecting with publicly available genome databases.

### 3a Identifying amplification primers

In many nucleic acid applications, we must amplify a genomic region to obtain enough material for detection. For example, the CRISPR-based detection platforms SHERLOCK^1^ and DETECTR^4^ use an isothermal approach, recombinase polymerase amplification (RPA), to amplify a target region; then, probes (in these applications, CRISPR guide RNAs) allow for target detection. Thus, the probes in a probe set *P* ought to be within a genomic region of the alignment that is bound by suitable primers to amplify the region (Supplementary Note Fig. 1).

In contrast to probes, our search for primers is related to a conventional approach that targets conserved regions and employs amplification-method–specific heuristics to filter primers. We identify suitable primers at every position of the alignment *S* by approximating a minimal set of primers that achieves a desired coverage over the input genomic variation. In particular, we run Algorithm 3 in Supplementary Note 1 at every site in the genome, except parameterized for primers. The parameters include primer length, number of tolerated mismatches, the fraction of genomes that must be covered, as well as bounds on GC content used as a filter; their values can be tuned based on a particular amplification method. See Methods for the particular parameter values that we use with ADAPT in practice, which we chose according to previously-published recommendations for RPA^77^.

### 3b Branch and bound search for genomic regions

In ADAPT, we perform a search for genomic regions to target (which may, optionally, include amplification primers) simultaneously with optimizing the probe objectives that are described in Supplementary Note 1. As with probes, we want to penalize the number of primers required to amplify a region because they can interfere with each other or require multiple reactions. Similarly, we wish to penalize the length of the region because longer regions are less efficient to amplify; penalizing the logarithm of length approximates the length-dependence of amplification efficiency. We first walk through the search using the objective that maximizes expected activity (Supplementary Note 1a). For this, we now perform a search for a genomic region *R* that encompasses the probe set *P* and solve

**Supplementary Note Figure 1.**
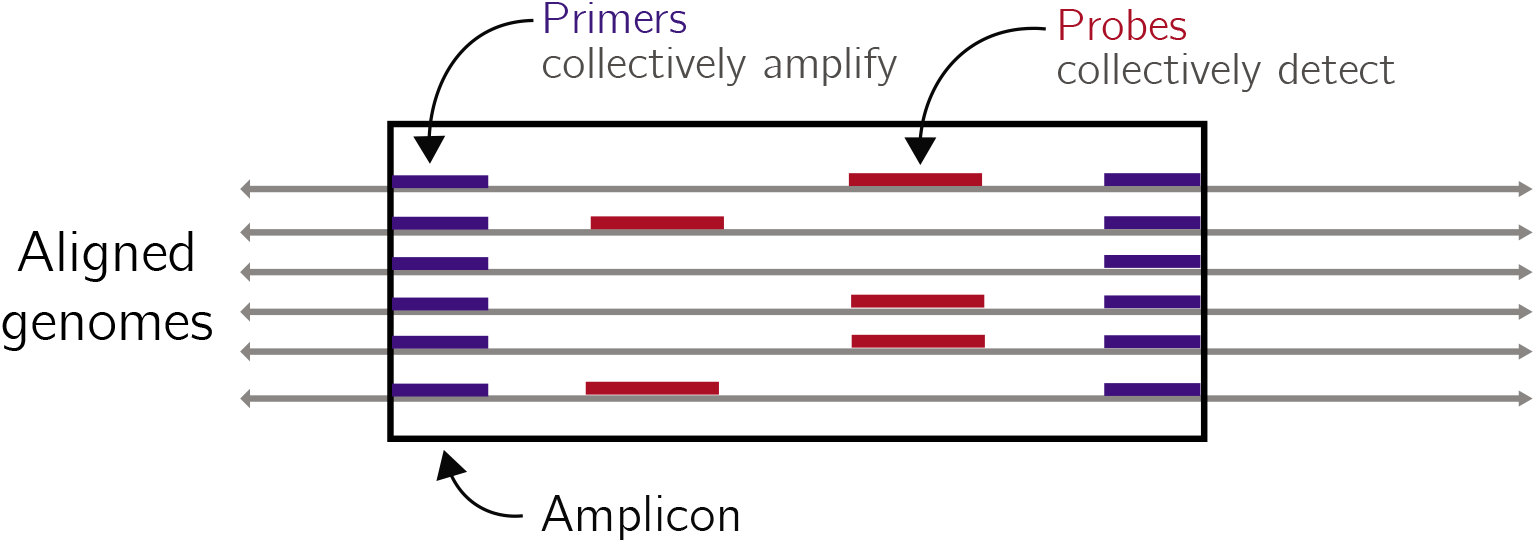
Searching for genomic regions. ADAPT searches for a region of the genome, bound by conserved sequence to use for primers, that contains probes that can collectively detect the region. The requirement that a region be bound by conserved sequence and represent an amplicon is optional.

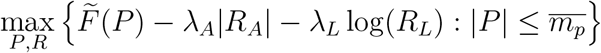

where 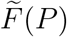 and 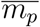 are defined in Supplementary Note 1a, *R_A_* gives the set of primers bounding the region, *R_L_* gives the nucleotide length of the region, and *λ_A_* and *λ_L_* give weights on the penalties^4^. Note that *λ_A_* and *λ_L_* can optionally be set to 0, removing the requirement that a region be bound by conserved sequence and represent an amplicon.

To solve this, we use an algorithm in which we search over options for *R* and prune unnecessary ones (Supplementary Note Fig. 2). Rather than finding a single maximum, we wish to compute the highest *N* solutions—i.e., *N* regions, each containing a probe set—to the objective. This is important so that multiple design options can be tested and compared experimentally; it also provides the option for an assay to target multiple regions in a genome. Note also that the range of 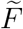 has an upper bound, which we call 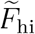, calculated from *F*(*P*)’s highest value (predicted activities are bounded) and |*P*| = 1.

We maintain a min heap *h* of the *N* designs with the highest value of the objective. First, we identify a primer set at every position in the alignment, as described in Identifying amplification primers. Then, we search over pairs of positions in the alignment, considering the regions that would be amplified by primers at each pair. Although the number of such regions is quadratic in the alignment length, we can effectively prune regions based on *R_A_*| and *R_L_*. We calculate, with these values, the objective value using 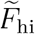 in place of 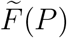; this value provides an upper bound on the solution. If this value falls below that of the minimum in *h*, the region cannot be in the top *N* and thus we do not need to compute *P*. For regions that could be in the best *N*, we compute the probe set *P* with the maximal 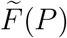 as described in Supplementary Note 1a (Solving for *P*). If the objective value for the design given by (*R, P*) is greater than the minimum in *h*, we pop from *h* and push the design to it. This search is guaranteed to identify the top *N* regions according to the objective, up to our approximation of 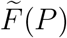.

This search follows the branch and bound paradigm in which the candidate solutions (*R, P*) make up a 3-level tree, excluding the root. The levels represent the (1) 5’ primers, (2) 3’ primers, and (3) probe set *P*. We can prune the choice of 3’ primers based on the length of the amplicon they would form. Exploring the final level in particular—determining *P*—is the slow step. Since we can easily construct an upper bound on the candidate solution for nodes in the final level, which we compare to the minimum in *h*, we can discard nodes and thus avoid having to compute *P* for many candidate solutions.

It is also important that the design options are diverse, i.e., reflect meaningfully different regions rather than being simple shifts of one another. To account for this, we implement the following: if a design to push to *h* has a region overlapping that of an existing design in *h*, it must replace that existing design (and only does so if the new one has a higher objective value).

Additionally, during our search many of the computations—particularly when computing probe sets—would be performed repeatedly from the same input, owing to overlap between different regions across the search. As a result, we memoize results of these probe set computations according to genome position. A branch and bound implementation, as described above, might start with all the 5’ primers (first level) and then select the best 3’ primers (second level), before advancing to computing probe sets. However, this could force each successive probe set computation to jump to a different region in the genome, as defined by the primer pairs: we would not be able to efficiently cleanup memoizations for these computations and memory would grow throughout the search. To avoid this issue, we scan linearly along the genome and, each time we advance the 5’ primer position, we determine probe set memoizations that we no longer need to store. Memoizing these computations provides a considerable improvement in runtime (Supplementary Fig. 23).

The above description applies to maximizing expected activity, but it is straightforward to adjust the strategy when minimizing the number of probes (Supplementary Note 1b). In this case, we change our objective to solve for

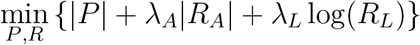

where we also impose the constraint on coverage described in Supplementary Note 1b. The search now stores a max heap *h* of the designs with the smallest values of the objective. For pruning, we compute a lower bound on the candidate solution by letting |*P*| = 1, and compare this bound to the maximum in *h*.

The search is embarrassingly parallel. One future direction is to parallelize the search across genomic regions, in which we perform it separately for contiguous parts of the genome and then merge the resulting heaps. In practice, the primary challenge is likely to be handling shared memory, in particular for the large index used to enforce specificity.

### 3c Fetching and curating sequences to target

ADAPT accepts a collection of taxonomies provided by a user: {*t*_1_, *t*_2_, … }. It can either design for one *t_i_* or for all *t_i_*, in either case ensuring designs are specific accounting for all *t_j_* where *j* ≠ *i*. Each *t_i_* generally represents a species, but can also be a subspecies taxon^5^. In NCBI’s databases, each taxonomy has a unique identifier^70^ and ADAPT accepts these identifiers. ADAPT then downloads all near-complete and complete genomes for each *t_i_* from NCBI’s genome neighbors database, but uses its Influenza Virus Resource database^71^ for influenza viruses. It also fetches metadata for these genomes (e.g., date of sample collection), which some downstream design tasks process.

We must then prepare these genomes for design. Briefly, for each *t_i_* we curate the genomes by aligning each one to one or more reference sequences^6^ for *t_i_* and remove genomes that align very poorly to all references, as measured by several heuristics: by default, we remove a genome that has < 50% identity to all references or that have < 60% identity to all references after collapsing consecutive gaps to a single gap. This process prunes genomes that are misclassified, have genes in an atypical sense, or are highly divergent for some other reason. Then, we cluster the genomes for *t_i_* with an alignment-free approach by computing a MinHash signature for each genome, rapidly estimating pairwise distances from these signatures (namely, the Mash distance^78^), and performing hierarchical clustering using the distance matrix. The default maximum inter-cluster distance (approximate average nucleotide dissimilarity) for clustering is 20%. In general, we obtain a single cluster for a species (Supplementary Fig. 27f). This provides another curation mechanism, because it can discard clusters that are too small (by default, just one sequence). Finally, ADAPT aligns the genomes within each cluster using MAFFT^102^. This yields a collection of alignments, where each is for a cluster of genomes from taxon *t_i_*.

**Supplementary Note Figure 2.**
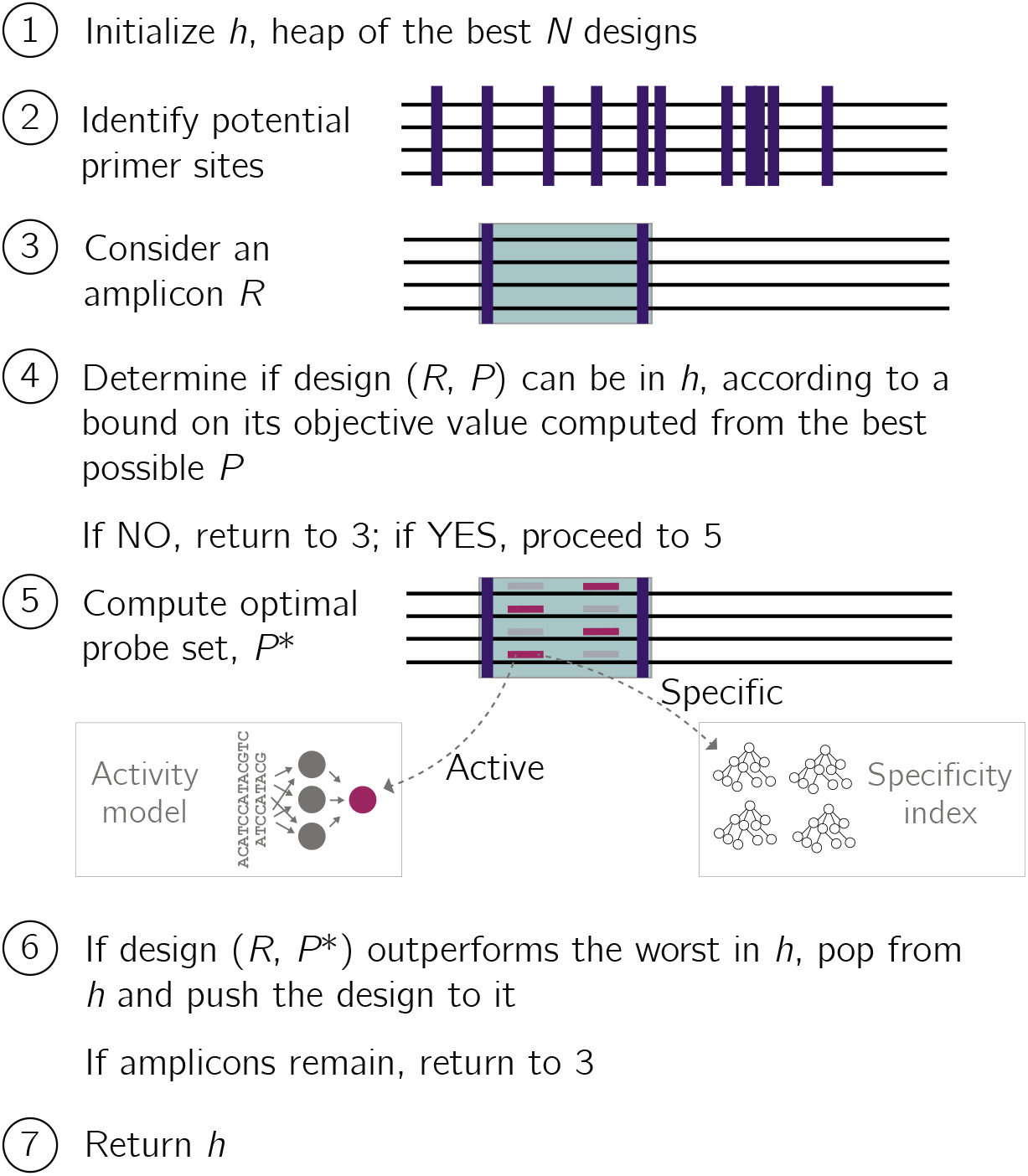
Branch and bound search for genomic regions. Sketch of the search for genomic regions (amplicons) and optimal probe sets within them. An amplicon *R* includes information about the primers used for amplifying it. Supplementary Note 1 describes the algorithms in Step 5. Step 5 makes use of the predictive activity model and the data structure (Supplementary Note 2) for evaluating specificity.

Many of these computations—such as curation, clustering, and alignment—are slow yet repeated on successive runs of ADAPT. ADAPT memoizes results of the above computations, to disk, to reuse on future runs when the input permits it. This memoization improves runtime for routine use of ADAPT.

## Supplementary Note 4

This note describes our approach to evaluate probe activity over time by simulating relatively likely substitutions. Results are in Supplementary Fig. 24.

### 4a Motivation

Nucleic acid diagnostics are susceptible to degraded performance as viral genomes accumulate substitutions. Extensive genomic data can inform where to design probes, for example, by identifying regions with less variability or regions where a probe (e.g., Cas13a guide) can maintain high activity across variation. However, finding such sites is difficult or impossible when there is little genomic data, as is the case early in an outbreak of a novel virus or for an understudied virus.

One option is to use a substitution model to simulate likely types of substitutions in the genome, and then to predict activity against these simulated sequences. Parameters of the model can be transferred from related viruses. The approach would account for the possibility that some potential target regions are more likely to accrue mutations that degrade activity than other regions. For example, consider a region rich in `T` nucleotides—complementary probes have `A`—and a virus with a high relative rate of `T` to `C` transitions. Also, consider a case where `A-C` probe-target mismatches harm activity, particularly in a specific location of the probe. Such substitutions in the genome would induce those mismatches; simulating these substitutions could inform ADAPT to avoid the regions or to position probes to avoid this type of mismatch.

### 4b Background on substitution model

We use the general time-reversible (GTR) model, which is based on a continuous-time Markov process that accounts for relative rates of substitutions. Ref. 103 contains further information, and this subsection summarizes the model. The parameters of this model are the equilibrium base frequencies, (*π_A_, π_C_, π_G_, π_T_*) and the rate parameters, *r_AC_, r_AG_, r_AT_, r_CG_, r_CT_, r_GT_*. Note that *r_ij_* = *r_ji_*. For convenience, denote the above parameters as *a, b, c, d, e, f* respectively.

The GTR model includes a rate matrix *Q* = {*q_ij_*}, giving the instantaneous rate at which a base *i* ∈ {A, C, G, T} changes to *j* ∈ {A, C, G, T} (i.e., the probability after an infinitesimal time):

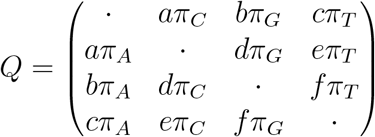

The diagonal entries are set so that each row sums to 0 and it is typical to normalize *Q* so that the average rate is 1 (i.e., 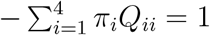).

The GTR model specifies a transition probability matrix *P*, computed numerically via matrix exponentiation:

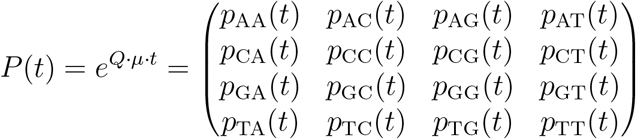

where *p_ij_* (*t*) is a probability that *i* transitions to *j* after elapsed time *t*. The overall substitution rate is *μ* and *μ* · *t* has units of expected number of substitutions per site. We use *P*, below, to simulate substitutions.

In analyses in Supplementary Fig. 24, we used *μ* = 10^-3^ substitutions/site/year and *t* = 5 years. We also empirically computed base frequencies from the SARS-CoV-2 genome used for those analyses and used the following relative rates (normalized to *r_GT_*), which we estimated: *r_AC_* = 1.3, *r_AG_* = 5.4, *r_AT_* = 1.8, *r_CG_* = 0.8, *r_CT_* = 9.5, and *r_GT_* = 1.0.

More sophisticated models—such as ones that accommodate rate variation among sites—may perform better for this task, but we have not experimented with them.

### 4c Evaluating probes against simulated sequences

Let *s* be the sequence of a virus at a given time, and *S_t_* be a discrete random variable representing the sequence of the virus after time *t* has elapsed. The above probability matrix *P* allows us to compute 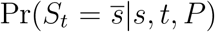, the likelihood of sequence 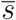 given the current sequence *s*. We use this model to construct a distribution of potential sequences after time *t*, and assess our probe’s activity against these sequences using our predictive model (namely, for Cas13a guides). For each pair of a probe *g* and original target sequence *s_i_*, we can obtain a sampling of activities,

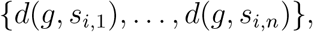

where each *s_i,j_* is one of *n* simulated sequences and *d*(*g, s_i,j_*) is a predicted detection activity.

**Algorithm 6.**
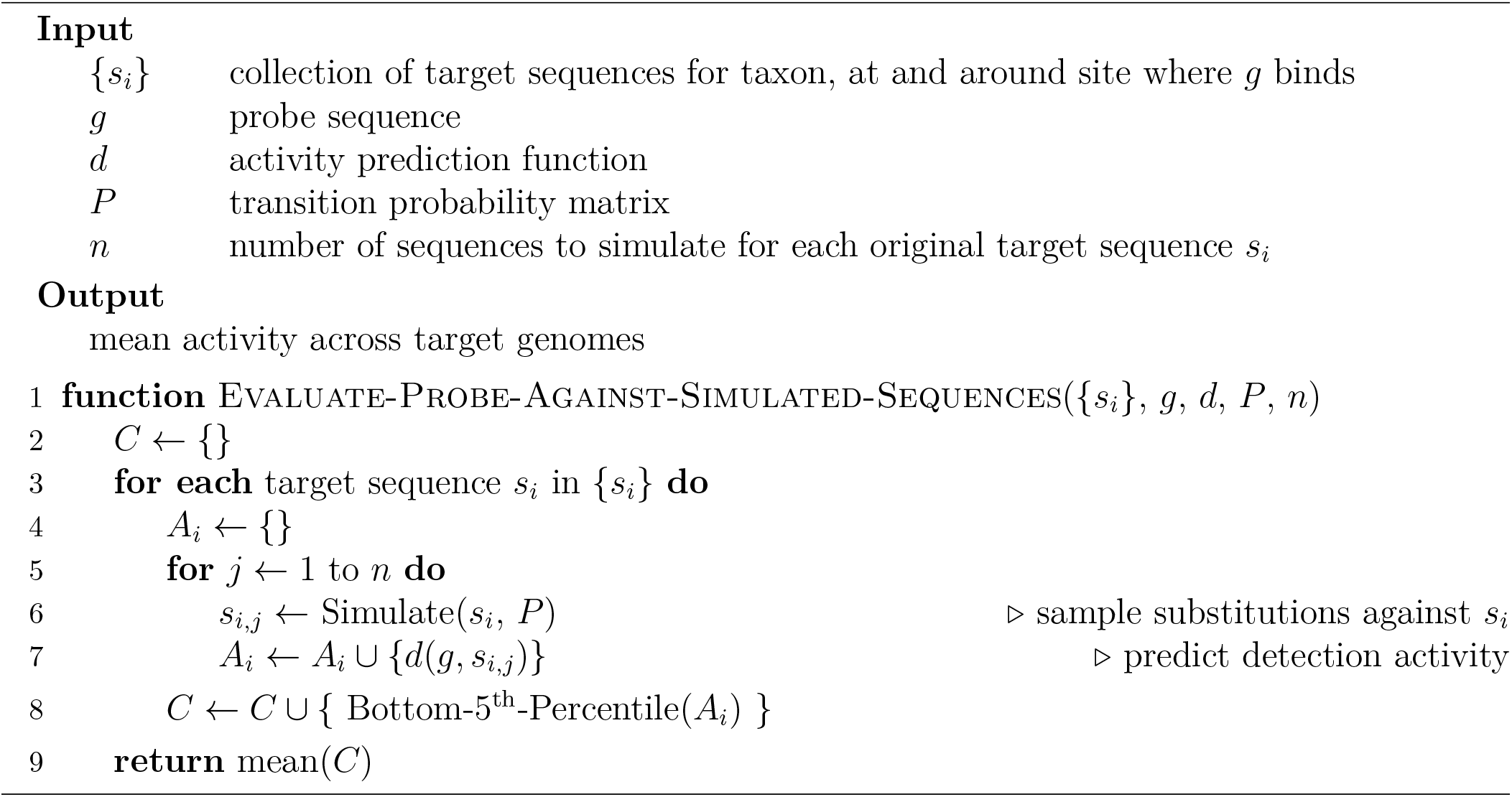
Estimate probe activity against simulated sequences.

We are often interested (Supplementary Fig. 24d) in whether a probe maintains activity against most of the potential substitutions—that is, we may want to be “risk-averse” and avoid a situation, even if unlikely, where we observe degraded activity owing to possible substitutions. For this goal, we use the bottom 5^th^ percentile of the different *d*(*g, s_i,j_*) to summarize the activity against simulated sequences originating with each *s_i_*. And then we summarize these across the different *s_i_* by taking the mean. Algorithm 6 shows pseudocode.

## Supplementary Figures

**Supplementary Figure 1.**
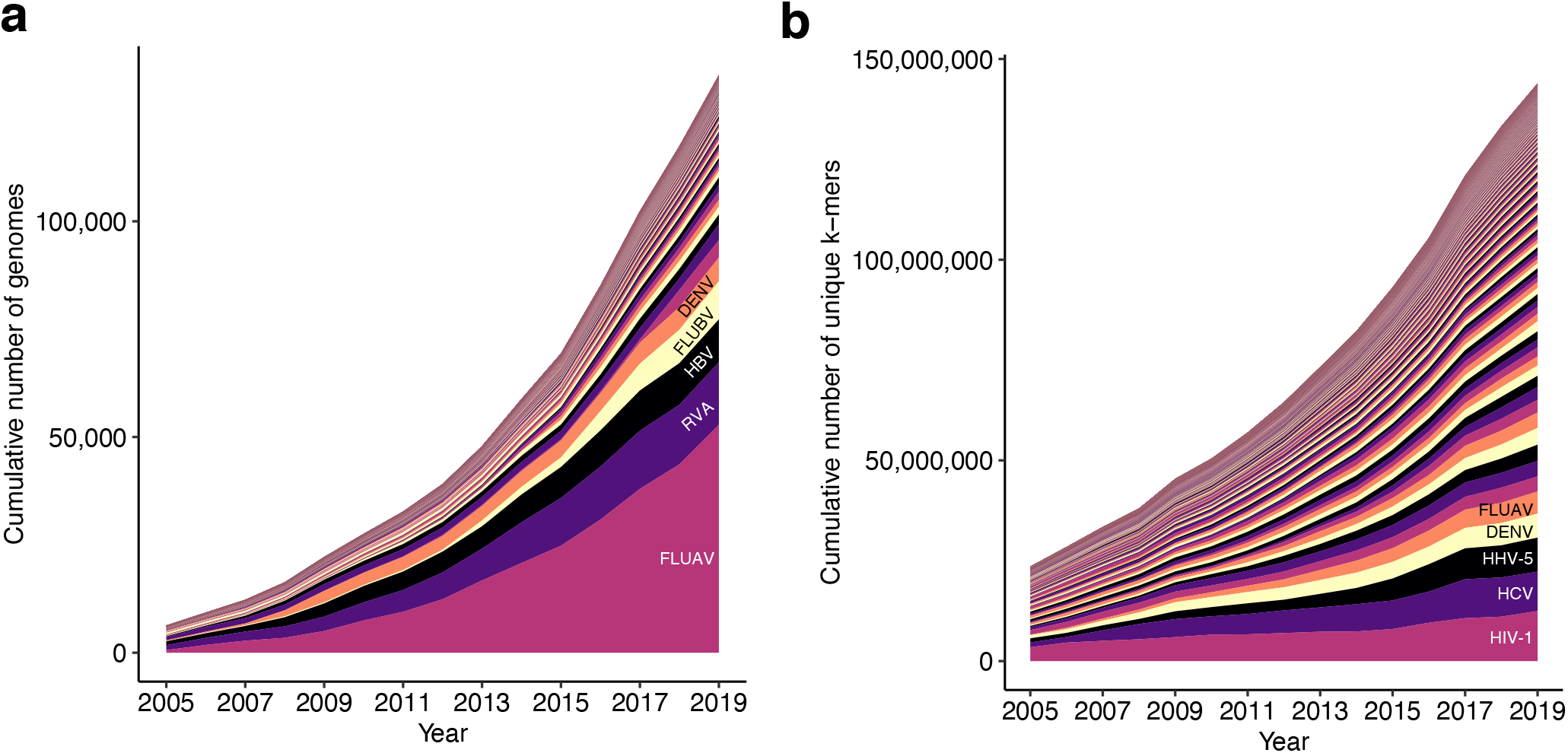
Growing number of viral genomes and diversity. Growth of data over time for 573 viral species known to infect humans. Each species is a color. **(a)** Cumulative number of genome sequences, counted from NCBI^31^ viral genome neighbors and influenza databases, for each species that were available up to each year. For genomes with multiple segments, this counts only the number of sequences of the segment that has the most sequences. 5 species with the most number of genomes are labeled. FLUAV, influenza A virus; RVA, rotavirus A; HBV, hepatitis B virus; FLUBV, influenza B virus; DENV, dengue virus. **(b)** Number of unique 31-mers for the genomes in (a), a simple measure of diversity. HIV-1, human immunodeficiency virus 1; HCV, hepatitis C virus; HHV-5, human betaherpesvirus 5. In both panels, year indicates the year of the entry creation date in the database.

**Supplementary Figure 2.**
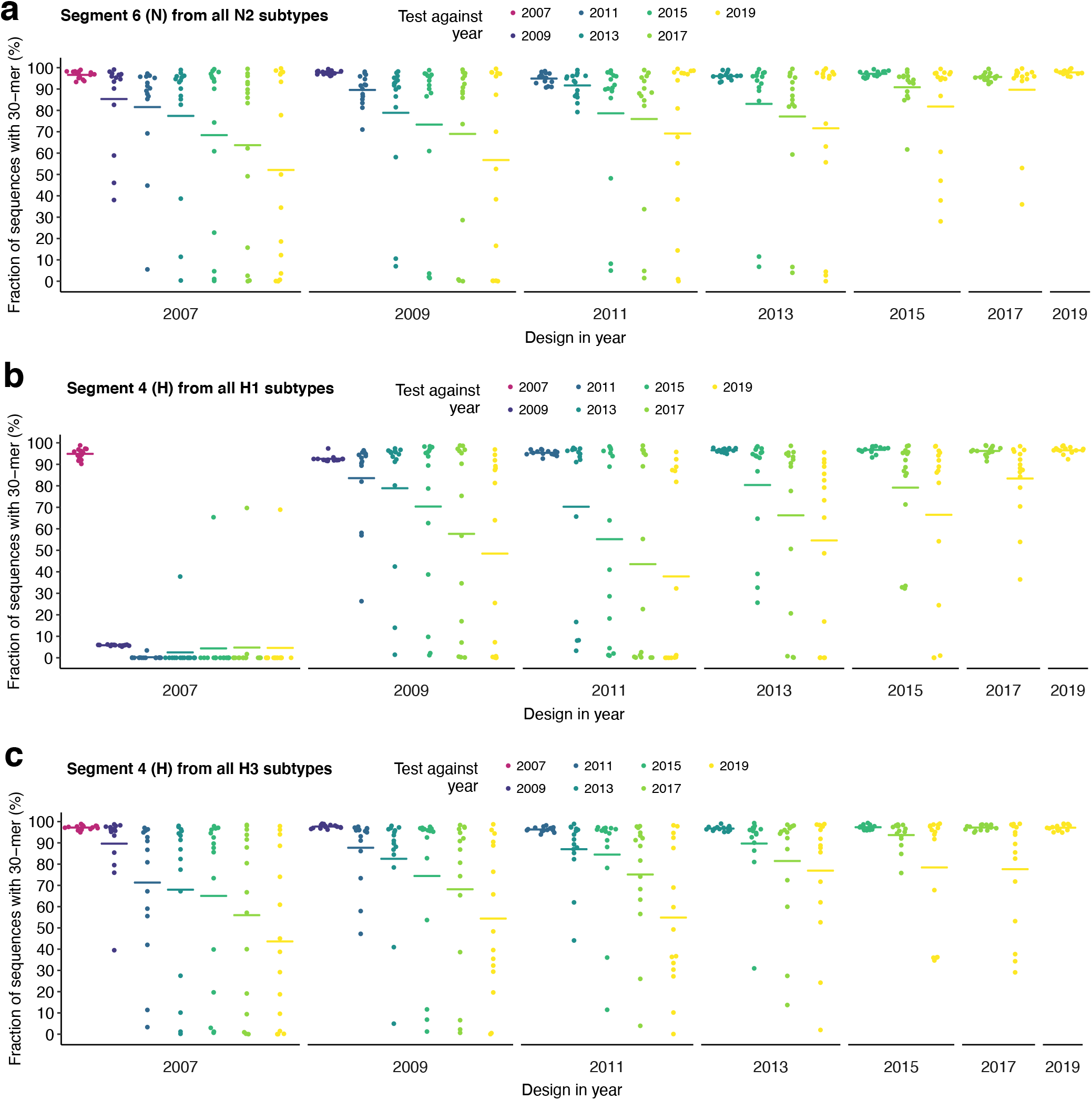
Comprehensiveness of conserved influenza A virus 30-mers over time. Even when considering the most conserved sequences, diagnostic performance of probes can degrade over time owing to genomic changes. At each year, we select the 15 most conserved non-overlapping 30-mers according to recent sequence data up to that year—a simple model for designing diagnostic probes at different years, without any consideration to other constraints such as specificity or activity. Each point represents a 30-mer from the year in which it was designed. We then measure the fraction of all sequences in subsequent years (colored) that contain each 30-mer—a simple test of comprehensiveness. Bars indicate the mean fraction of sequences containing the 15 30-mers at each combination of design and test year. To aid visualization, only odd years are shown. **(a)** Segment 6 (N) sequences from all N2 subtypes. **(b)** Segment 4 (H) sequences from all H1 subtypes. **(c)** Segment 4 (H) sequences from all H3 subtypes. Fig. 1a shows segment 6 (N) sequences from N1 subtypes.

**Supplementary Figure 3.**
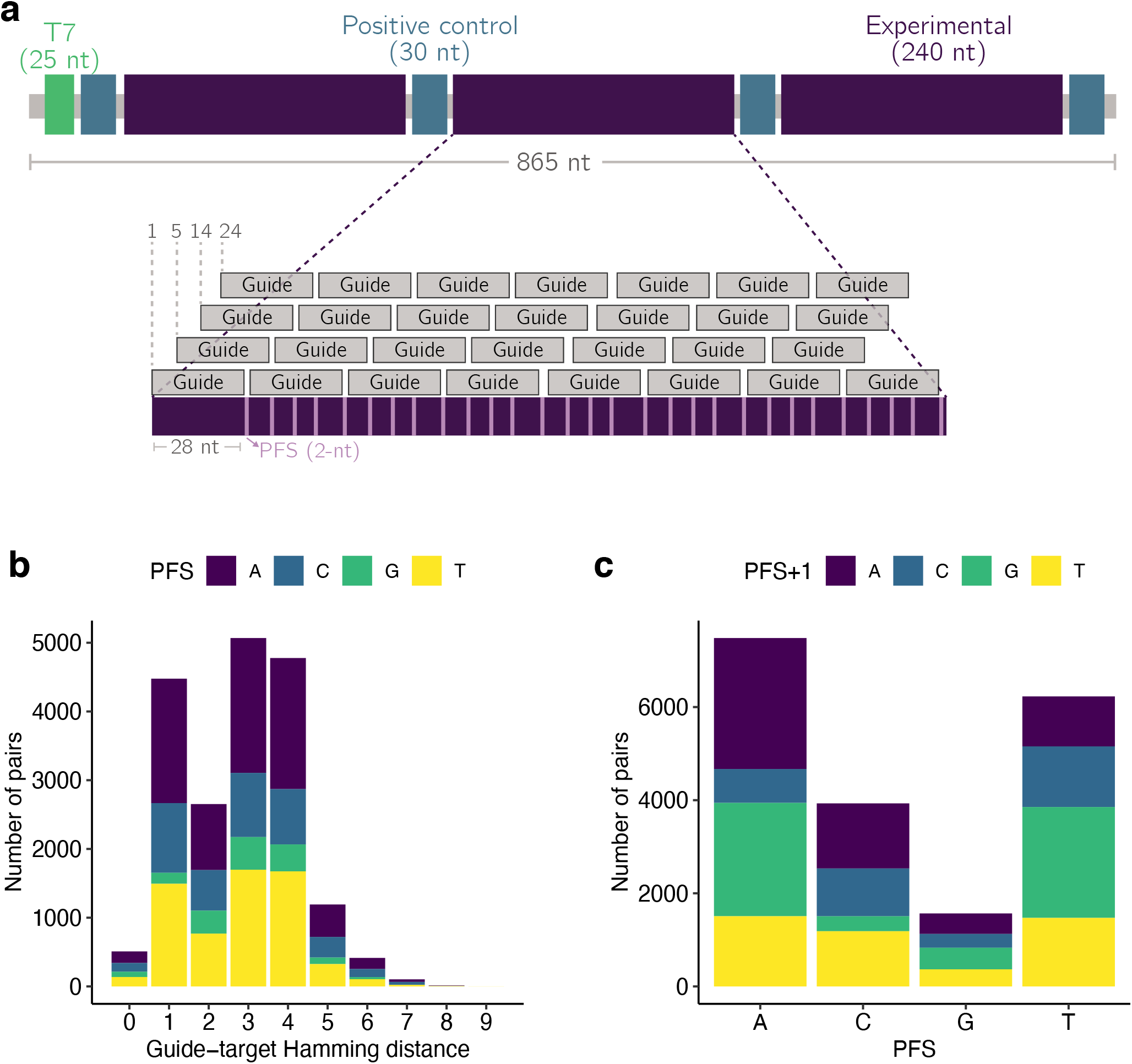
Guide-target library design. **(a)** Top, depiction of the wildtype target. The wildtype contains a T7 promoter on the 5’ end for transcription, four positive control regions, and three experimental regions. Each positive control region contains a unique guide that matches perfectly all targets, except negative control targets (not shown). Bottom, zoom of one experimental region from the wildtype target. Guide sequence comes from tiling along this region—29 guides per experimental region. There are also negative control guides (not shown) that only match the negative control targets. Other targets contain mismatches relative to the wildtype target, and thus contain mismatches relative to the guide sequences. **(b)** Distribution of the Hamming distance between guide and target across guide-target pairs. Color represents the number of pairs with each protospacer flanking site (PFS) at each Hamming distance. The 19,209 unique guide-target pairs included in our final, curated dataset (Methods) are shown. **(c)** Same as (b), but the distribution of PFS across the guide-target pairs. Color represents the number of pairs with each nucleotide immediately following the PFS (3’ end of protospacer).

**Supplementary Figure 4.**
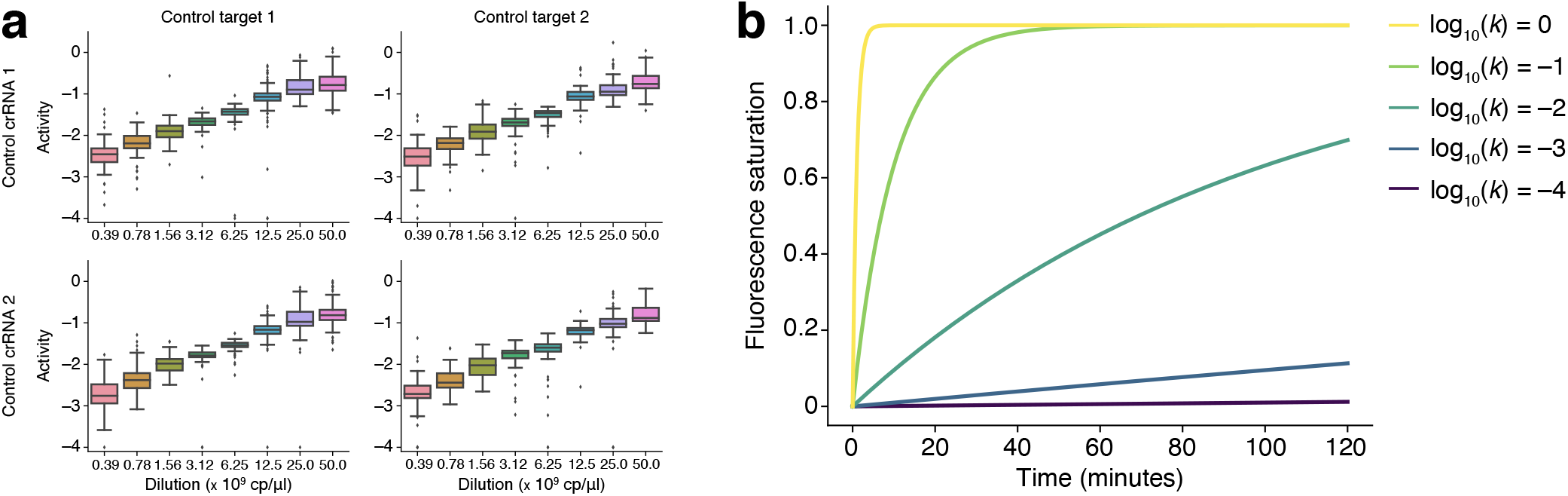
Assessing activity through CRISPR-Cas13a reaction kinetics. **(a)** Cas13 activity for a series of target concentrations, using two control targets and guides from our Cas13 library. We model fluorescence for each guide-target pair over time (Fig. 2a; Methods), fitting a curve of the form *C*(1 – *e^−kt^*) + *B* where *t* is time and *e^-kt^* represents remaining reporter presence over time. We take log_10_(*k*) to be the measure of Cas13 activity. Boxes show quartiles and whiskers extend, past the low/high quartiles, up to 1.5 times the interquartile range. **(b)** Theoretical fluorescence saturation over time—namely, the term 1 – *e^−kt^*—for five activity values. Over the time scale of our experiment (*t* up to ~120 minutes), when *k* is small we cannot observe reporter activation and the curve is approximately linear, making it difficult to estimate C and k together; these features motivate the use of an activity cutoff. Therefore, we label guide-target pairs with log_10_(*k*) ≤ −4 as inactive and those with log_10_(*k*) > −4 as active.

**Supplementary Figure 5.**
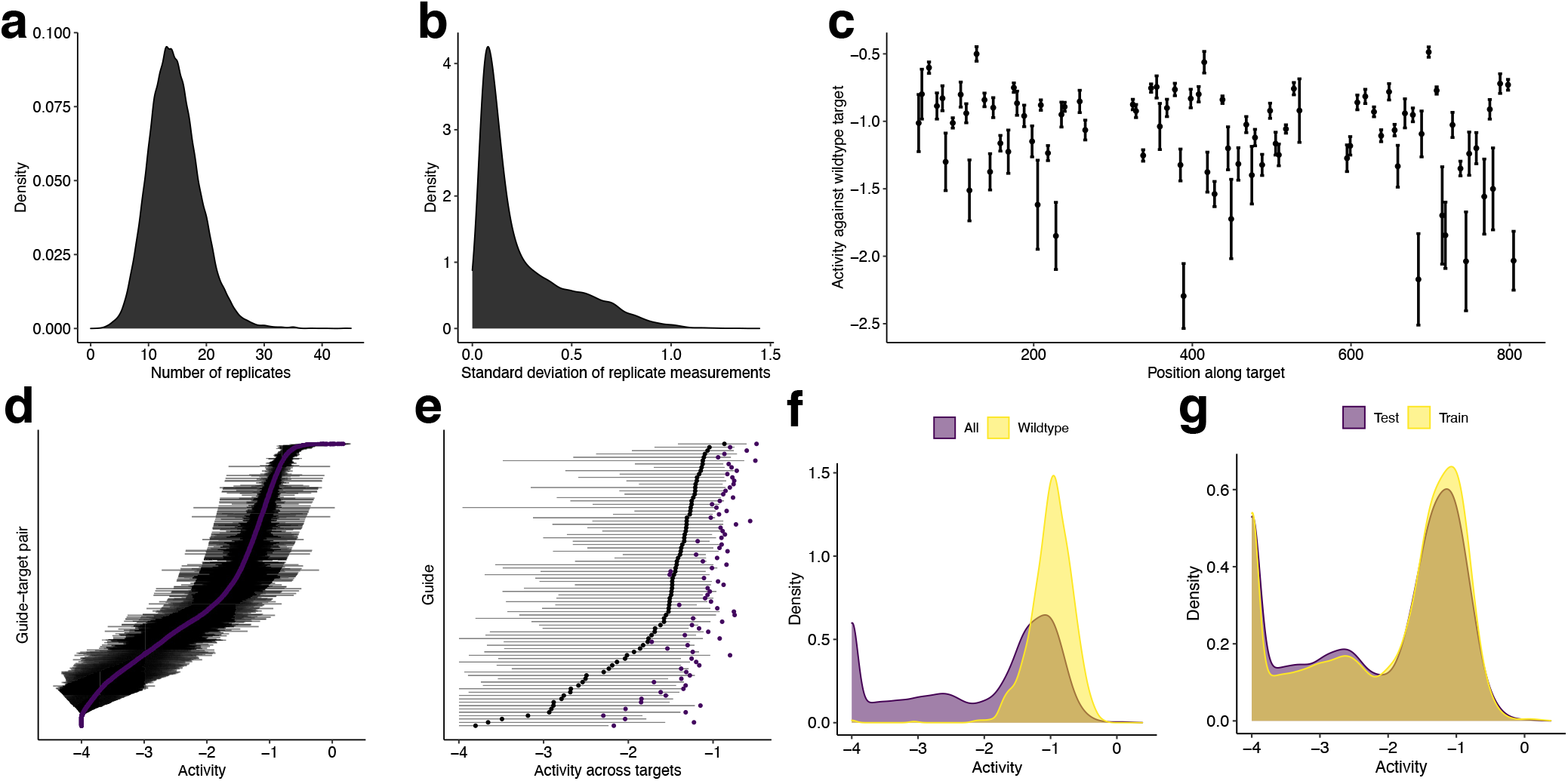
Dataset of CRISPR-Cas13a guide-target pairs. All panels show the 18,508 unique guide-target pairs in the dataset used for training and testing. Activity is defined in Methods. **(a)** Distribution of number of replicate activity measurements for each pair. **(b)** Distribution of standard deviation across replicate activity measurements for each pair. **(c)** Activity of each guide against the wildtype target (matching exactly), shown by their position along the target. Dot indicates the mean activity across the wildtype targets, shown with a 95% confidence interval. **(d)** Variation in activity across guide-target pairs and among replicate measurements. Each row represents a guide-target pair. Purple dot indicates the mean activity across replicate measurements; pairs are sorted vertically by this value. Bars indicate the 95% confidence interval for the mean. **(e)** Variation in activity between guides and across targets for each guide. Each row represents a guide. Black dot indicates the median activity across all targets and bars span the 20^th^ and 80^th^ percentiles of activity across all targets. Purple dot indicates the mean activity across the wildtype targets (matching the guide exactly). **(f)** Distribution of activity across all guide-target pairs and only pairs with the wildtype target. **(g)** Distribution of activity across all guide-target pairs in the training data and the pairs in the test data (the two sets do not overlap along the target or contain the same guides; Methods). In (d–g), there are 10 resampled replicate activity measurements for each guide-target pair. We set a lower threshold of −4 on the activity owing to measurement limitations (see Supplementary Fig. 4b and Methods for details), so density shown at −4 includes guide-target pairs with true activity below this threshold. (a), (b), (f), and (g) show kernel density esimates.

**Supplementary Figure 6.**
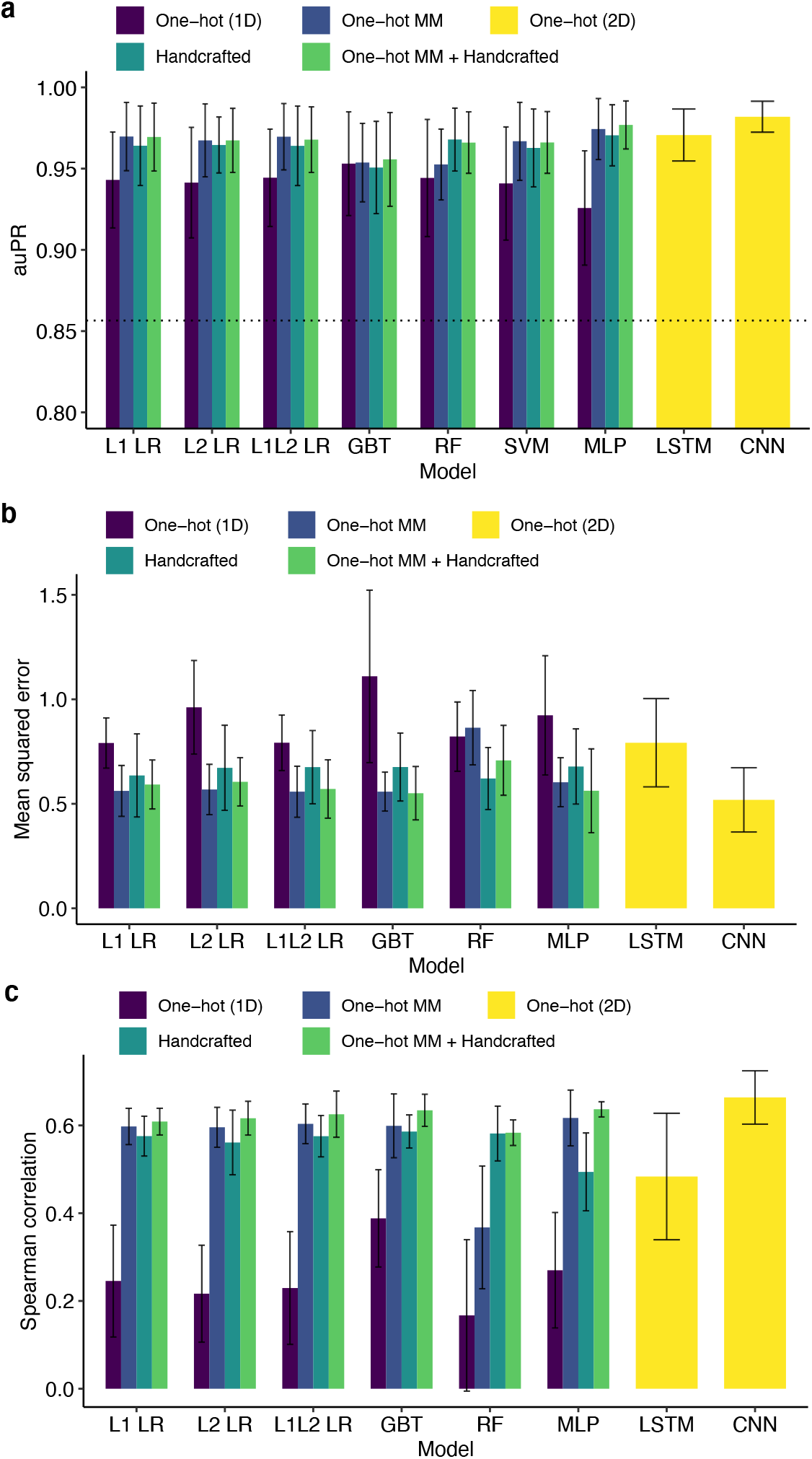
Nested cross-validation for classification and regression. For each model and input type (color) on each of five outer folds, we performed a five-fold cross-validated hyperparameter search. The plotted value is the mean of a statistic across the five outer folds, and the error bar indicates the 95% confidence interval. **(a)** Area under precision-recall curve (auPR) for different classification models. auROC is in Fig. 2c. L1 LR and L2 LR, logistic regression; L1L2 LR, elastic net; GBT, gradient-boosted classification tree; RF, random forest; SVM, support vector machine; MLP, multilayer perceptron; LSTM, long short-term memory recurrent neural network; CNN, convolutional neural network including parallel convolution filters of different widths and a locally-connected layer. One-hot (1D) is one-hot encoding of target and guide sequence independently, i.e., without encoding a pairing of nucleotides between the two; One-hot MM is one-hot encoding of target sequence nucleotides and of mismatches in guides relative to the target; Handcrafted is curated features of hypothesized importance (Methods); One-hot (2D) is one-hot encoding of target and guide sequence with encoded guide-target pairing. Dashed line is precision of random classifier (equivalently, the fraction of guide-target pairs that are active). **(b)** Mean squared error (MSE) for different regression models (lower is better). L1 and L2 LR, regularized linear regression; L1L2 LR, elastic net; GBT, gradient-boosted regression tree; RF, MLP, LSTM, and CNN are as in (a) except constructed for regression. Input types are as in (a). **(c)** Same as (b) but the statistic is Spearman correlation.

**Supplementary Figure 7.**
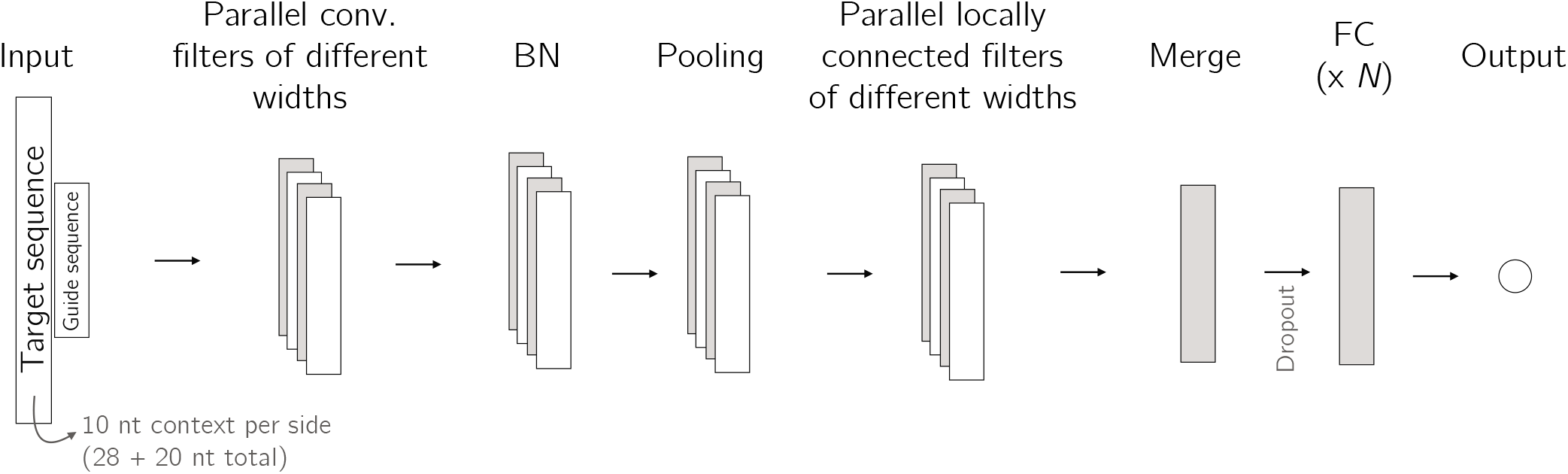
Architecture of convolutional neural network for guide-target activity prediction. Convolutional neural network (CNN) architecture for classifying and regressing activity; hyperparameter search and training is separate for each task. The inputs are one-hot encoded for the target and guides sequences (8 channels in total). There are multiple convolutional filters of different widths processing the input in parallel, as well as multiple locally connected filters of different widths; outputs of these different filters are concatenated in the merge layer. Pooling includes maximum, average, and both. ‘BN’ is batch normalization and ‘FC’ is fully connected. There are *N* fully connected layers. The dropout layers are in front of each fully connected layer.

**Supplementary Figure 8.**
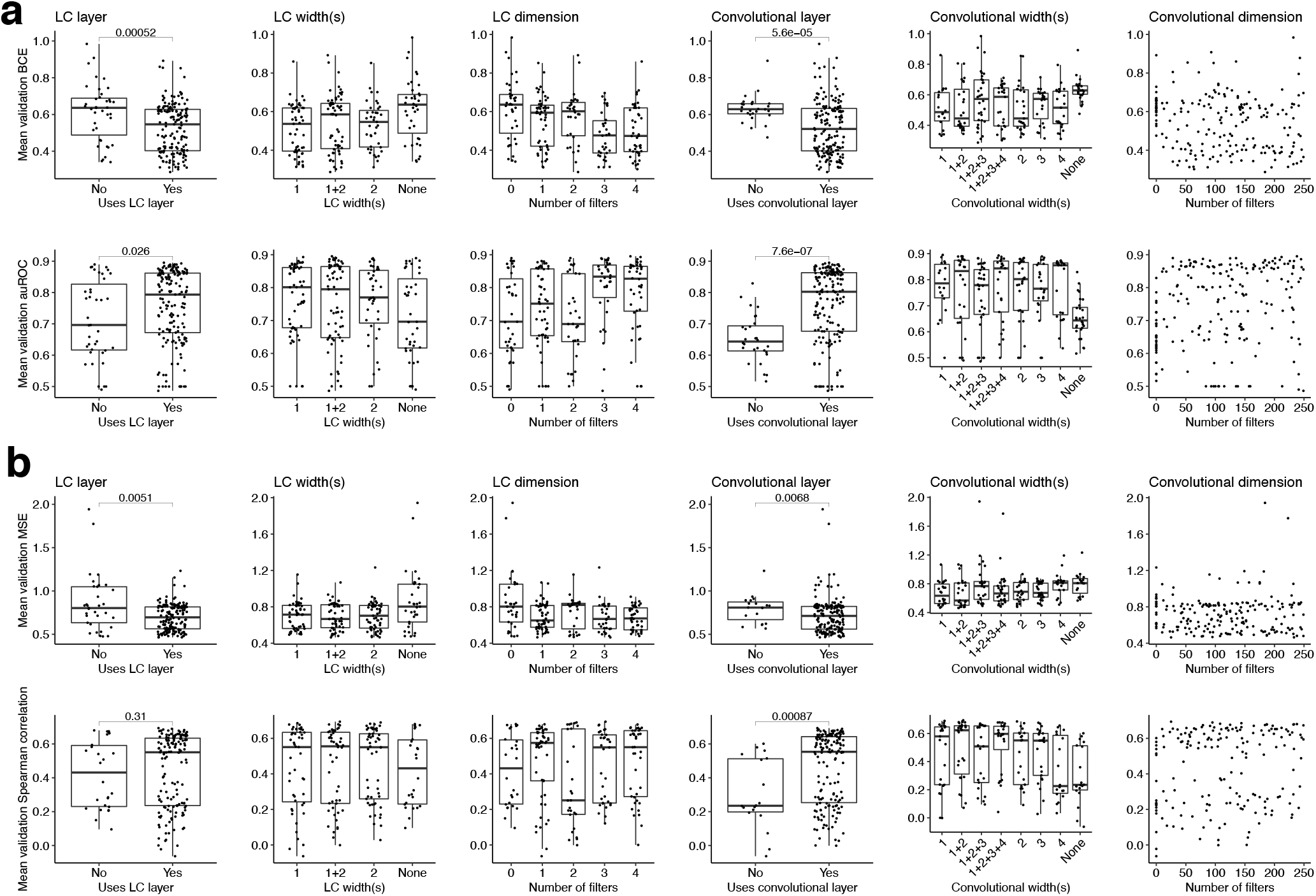
Hyperparameter search for convolutional neural networks. We used a random search over the hyperparameter space (200 draws) to select each convolutional neural network (CNN) model. Each plot corresponds to a hyperparameter and shows choices of that hyperparameter; see Methods for all hyperparameters. The evaluations are cross-validated: each dot indicates the mean of a metric, computed across five folds, for a draw of hyperparameters. ‘LC’, locally connnected. The ‘+’ in LC and convolutional widths separates different widths of parallel filters; ‘None’ indicates that the model does not use an LC or convolutional layer. *P*-values are computed from Mann-Whitney *U* tests (one-sided). **(a)** Results of hyperparameter search for classification. BCE, binary cross-entropy. **(b)** Results of hyperparameter search for regression. MSE, mean squared error.

**Supplementary Figure 9.**
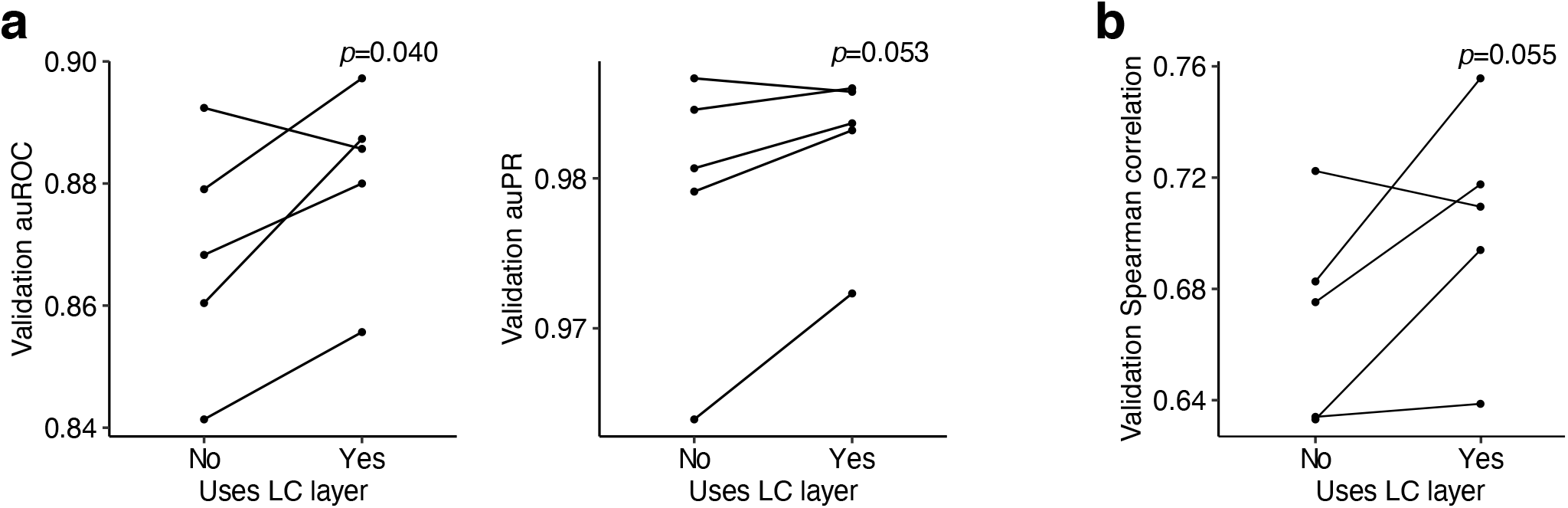
Effect of locally connected layers on model performance. Results of the forced inclusion or exclusion of locally connected layers (‘LC’; Supplementary Fig. 7) in convolutional neural networks for Cas13a guide-target activity prediction. We perform nested cross-validation: on each of five outer folds, we perform a five-fold cross-validated hyperparameter search to select a model, once using locally connected layers and once not using them. Plotted values are a statistic calculated on the validation data for each of the five outer folds. **(a)** auROC and auPR for classifying activity. **(b)** Spearman correlation for regressing activity on active guide-target pairs. *p*-values are computed from one-sided paired *t*-tests.

**Supplementary Figure 10.**
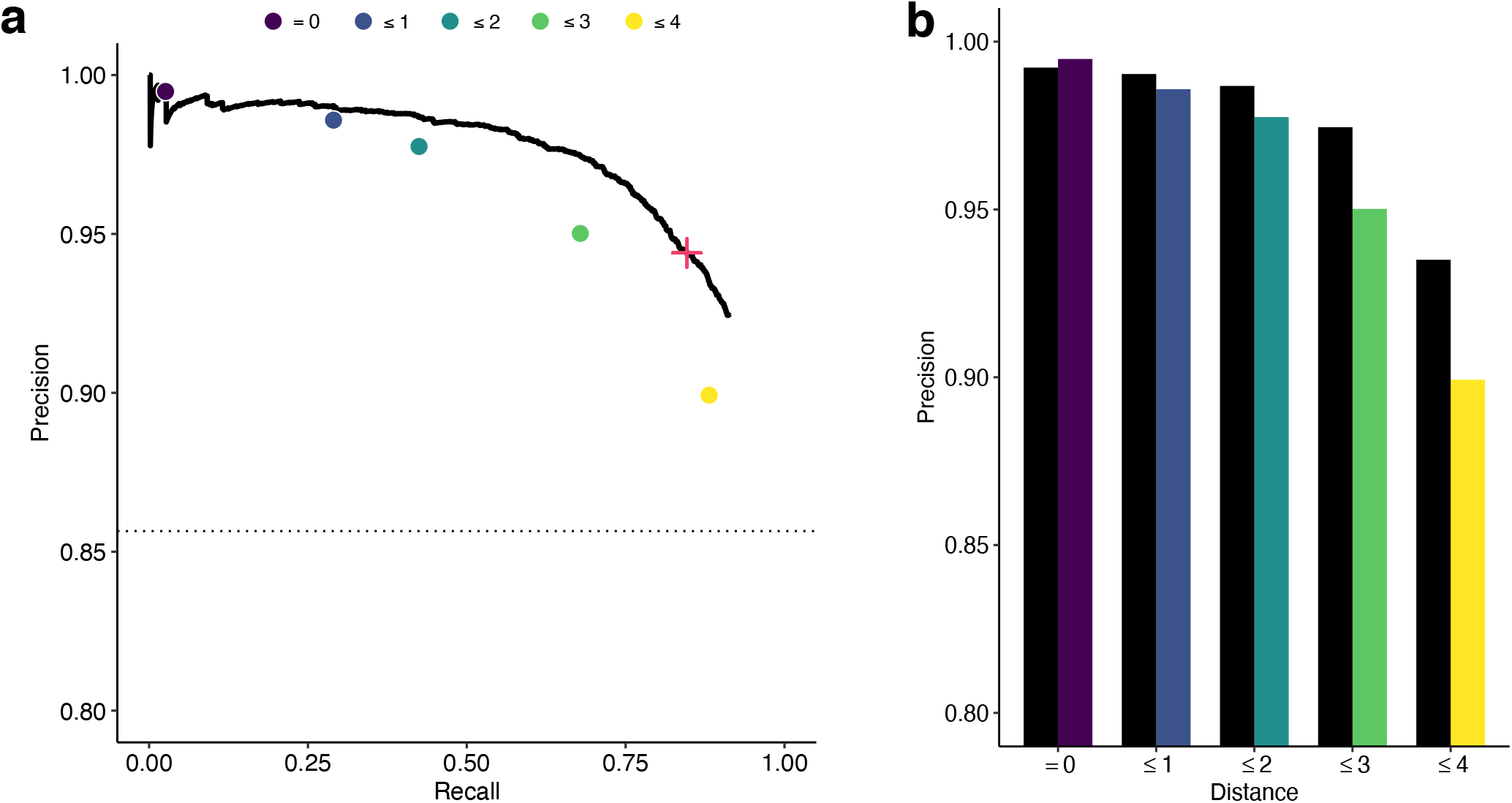
Precision-recall curve of classifier. **(a)** Precision-recall (PR) curve of CNN model, which is used in ADAPT, classifying pairs as inactive or active on a held-out test set. ROC curve is in Fig. 2d. Points indicate precision and recall for baseline heuristic classifiers, defined as choosing a guide-target pair to be active if and only if it has an active (non-`G`) PFS and the Hamming distance between the guide and target is less than the specified threshold (color). Red ‘+’ indicates the decision threshold in ADAPT. Dashed line is precision of a random classifier (equivalently, the fraction of guide-target pairs that are active). **(b)** Comparison of precision between CNN (black) and baseline classifiers (color as in (a)) at equivalent recall.

**Supplementary Figure 11.**
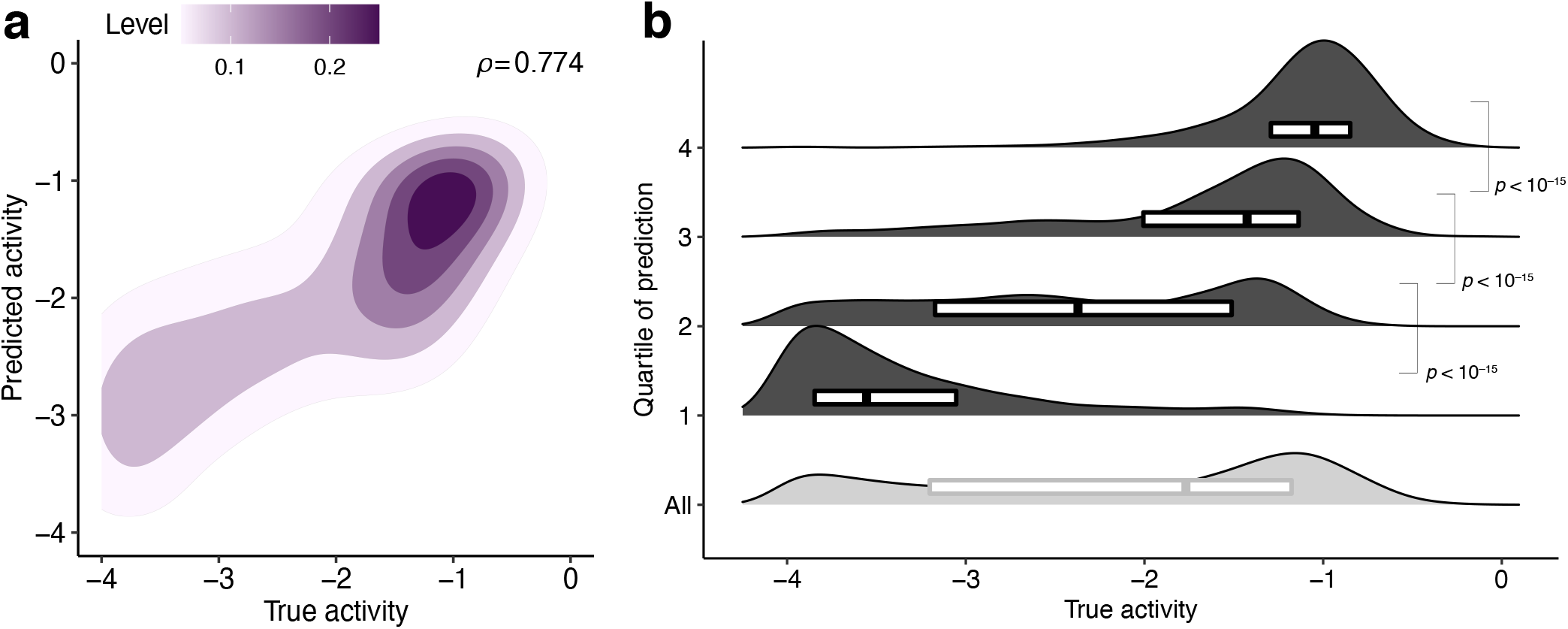
Regression results using all guide-target pairs. Results, on the held-out test set, of a CNN trained to regress activity using *all* guide-target pairs (other regression data are trained and tested only on active pairs). We set a lower threshold of −4 on the activity owing to measurement limitations (see Supplementary Fig. 4b and Methods for details), so activities are bounded below at −4. **(a)** Contour color, point density. *ρ*, Spearman correlation. **(b)** Same data as (a). Each row contains one quartile of pairs divided by predicted activity (top row is predicted most active), with the bottom row showing all guide-target pairs. Smoothed density estimates and interquartile ranges show the distribution of true activity for the pairs from each quartile. *P*-values are calculated from Mann-Whitney *U* tests (one-sided). The excess of inactive guide-target pairs in our data distorts the performance of this model and we do not use this model in ADAPT. We instead use the two-step hurdle model (Fig. 2d and 2e), as described in Methods, owing to the data’s distribution and the process we aim to model.

**Supplementary Figure 12.**
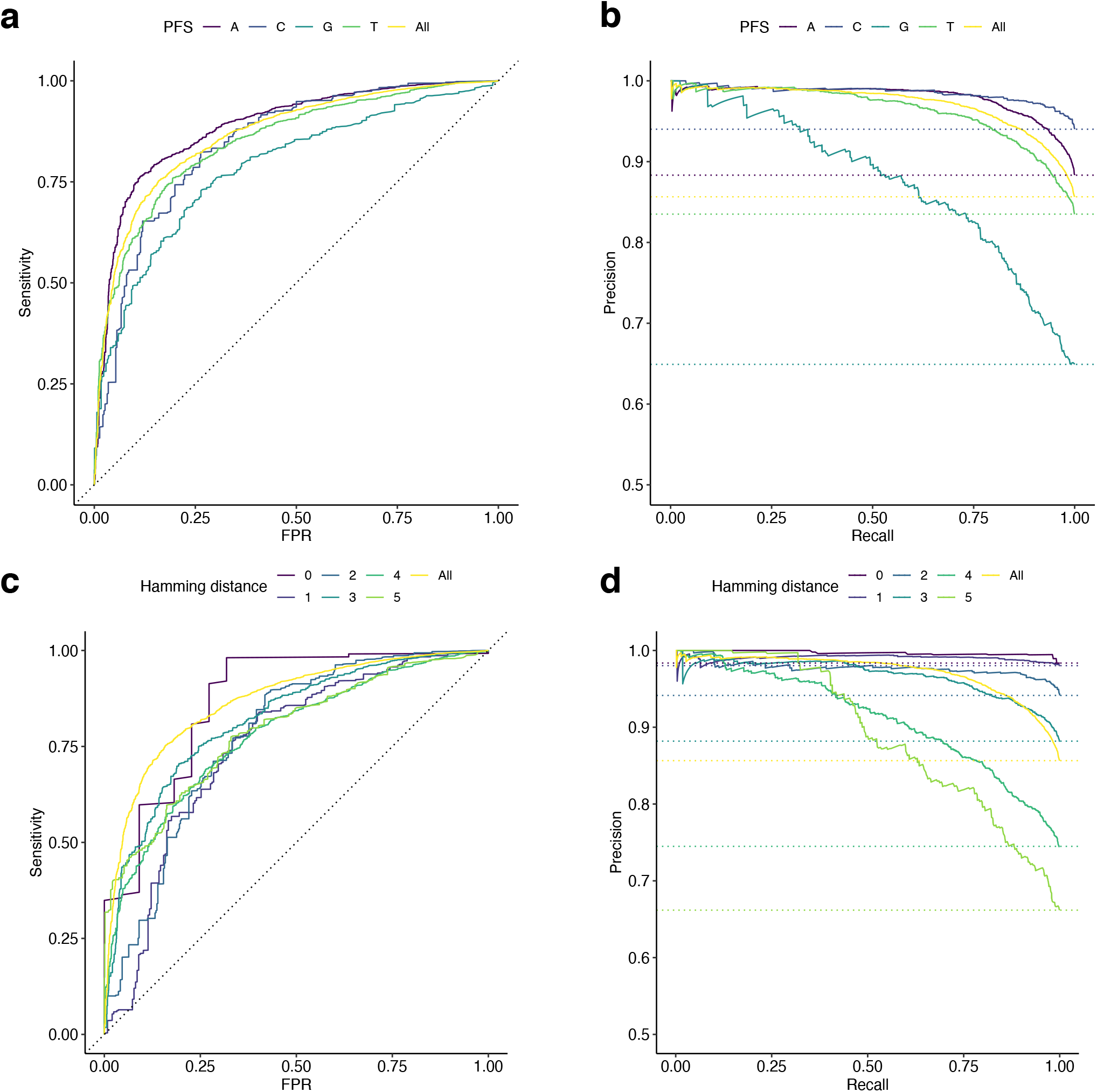
Classification performance on subsets of test data. Evaluations of classification on different subsets of the held-out test data corresponding to features that considerably affect activity. Here, the model is the same for all evaluations and only tested (not trained) on different subsets. **(a)** ROC curves computed from guide-target pairs with the different protospacer flanking site (PFS) nucleotides. **(b)** Precision-recall (PR) curves computed from pairs with the different PFS nucleotides. Dashed lines are precision of random classifiers for each PFS (equivalently, the fraction of guide-target pairs that are active with each PFS). **(c)** ROC curves computed from pairs with different Hamming distances between guide and target. **(d)** PR curves computed from pairs with different Hamming distances between guide and target. Dashed lines are precision of random classifiers for each choice of Hamming distance (equivalently, the fraction of guide-target pairs that are active at each Hamming distance). In all panels, yellow curve is for all test data.

**Supplementary Figure 13.**
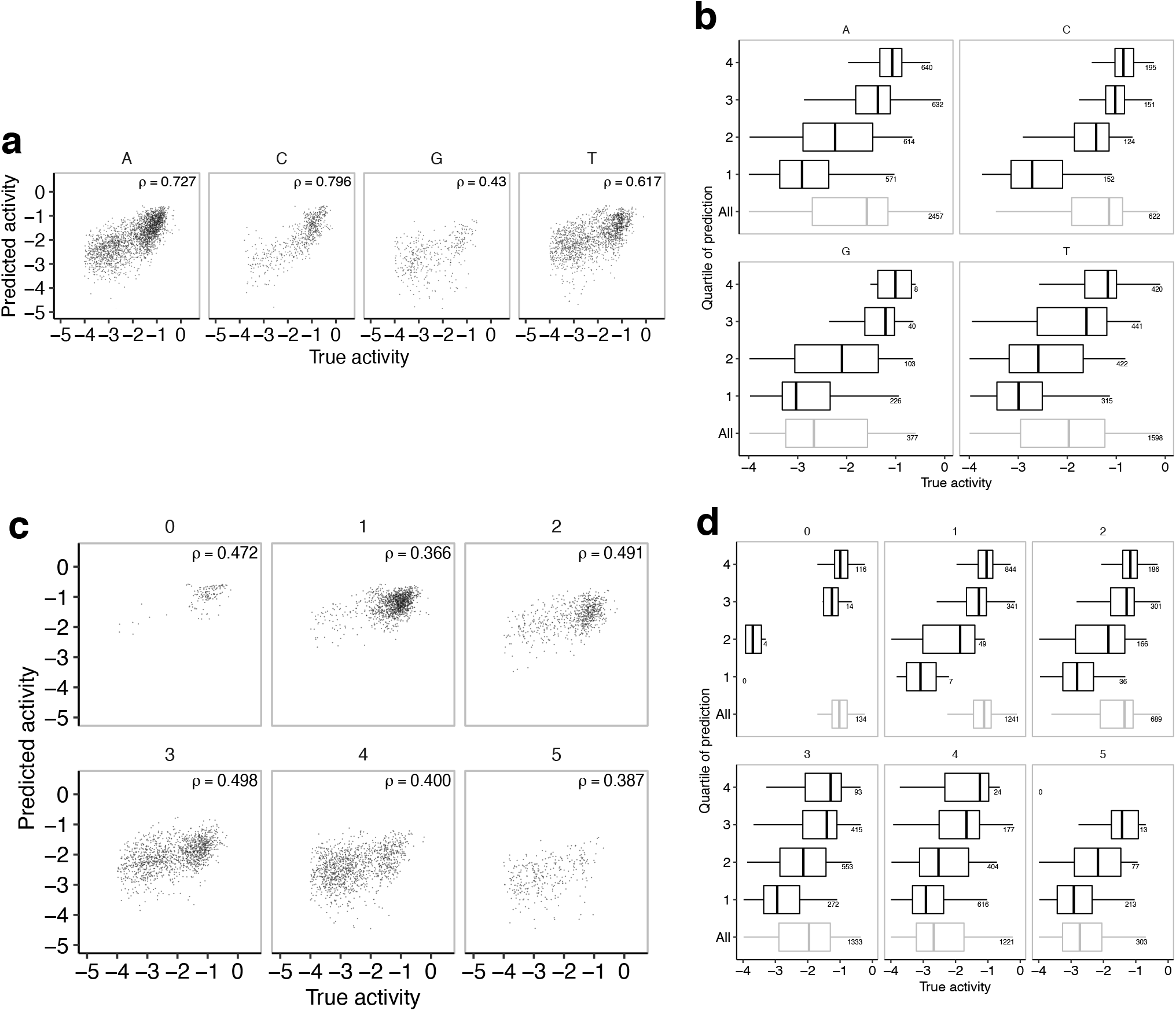
Regression performance on subsets of test data. Evaluations of regression on different subsets of active guide-target pairs in the held-out test data, where the subsets correspond to variables that considerably affect activity. Here, the model is the same for all evaluations and only tested (not trained) on different subsets. **(a)** Pairs separated by the different protospacer flanking site (PFS) nucleotides, indicated above each plot. Each point is a guide-target pair. **(b)** Pairs separated by the different PFS nucleotides. Each row contains one quartile based on their predicted activity (top row is predicted most active), with the bottom row showing all active pairs with the PFS. Numbers indicate the number of pairs in each quartile; the quartile for each pair is based on its predicted activity across all PFS nucleotides, not only the PFS for each plot. **(c)** Same as (a), except separated by different Hamming distances between guide and target. **(d)** Same as (b), except separated by Hamming distance. In (a) and (c), *ρ* is Spearman correlation. In (b) and (d), boxes show quartiles and whiskers extend, past the low/high quartiles, up to 1.5 times the interquartile range (outliers not shown).

**Supplementary Figure 14.**
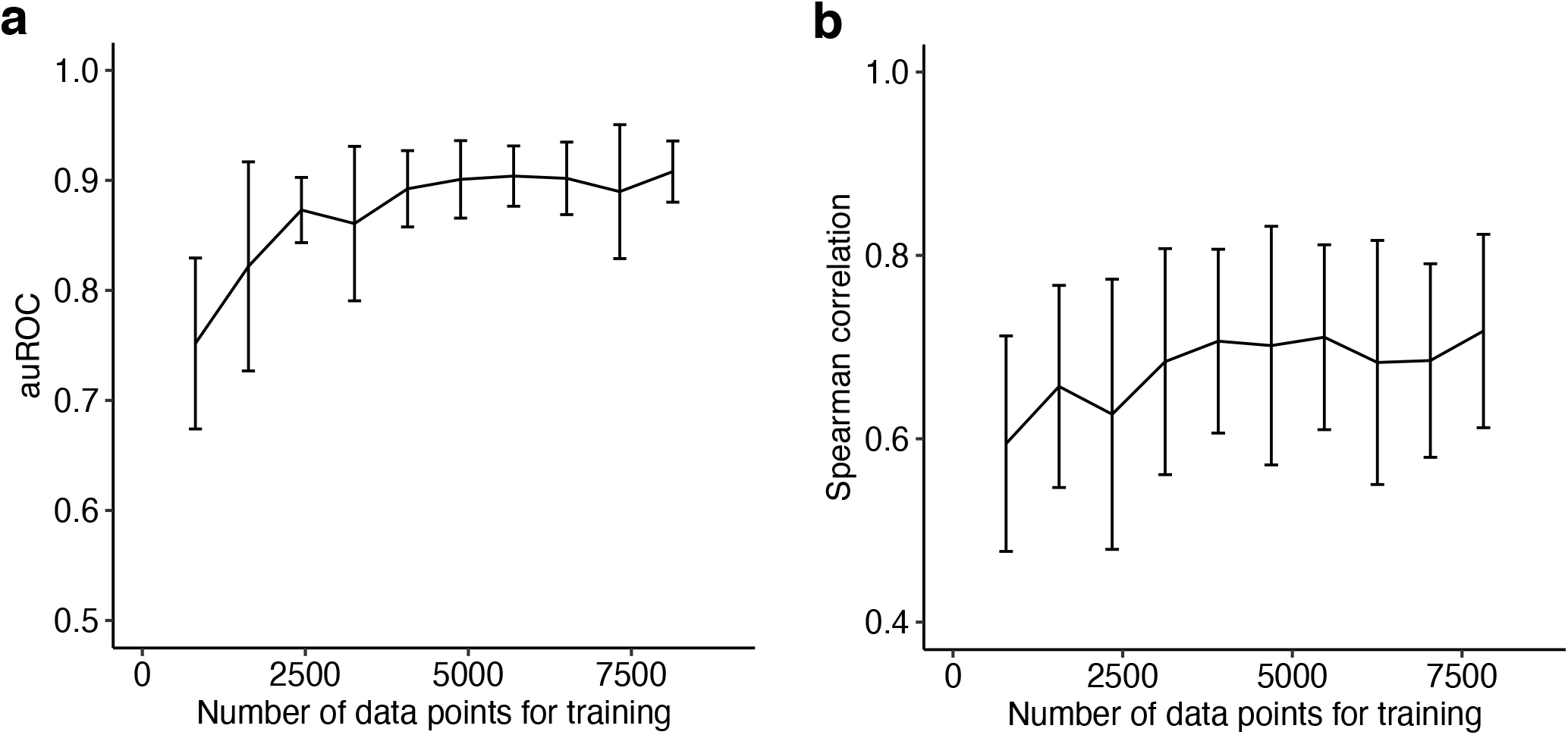
Learning curves. Learning curves for the convolutional neural networks used in ADAPT, which assess whether additional data could benefit model performance. At each number of input training data points, we perform nested cross-validation to select models: on each of five outer folds, we perform a five-fold cross-validated hyperparameter search to select a model. Line indicates the mean of a statistic on the validation data across the five selected models and error bars give a 95% confidence interval. **(a)** Learning curve selecting models for classification. **(b)** Learning curve selecting models for regression.

**Supplementary Figure 15.**
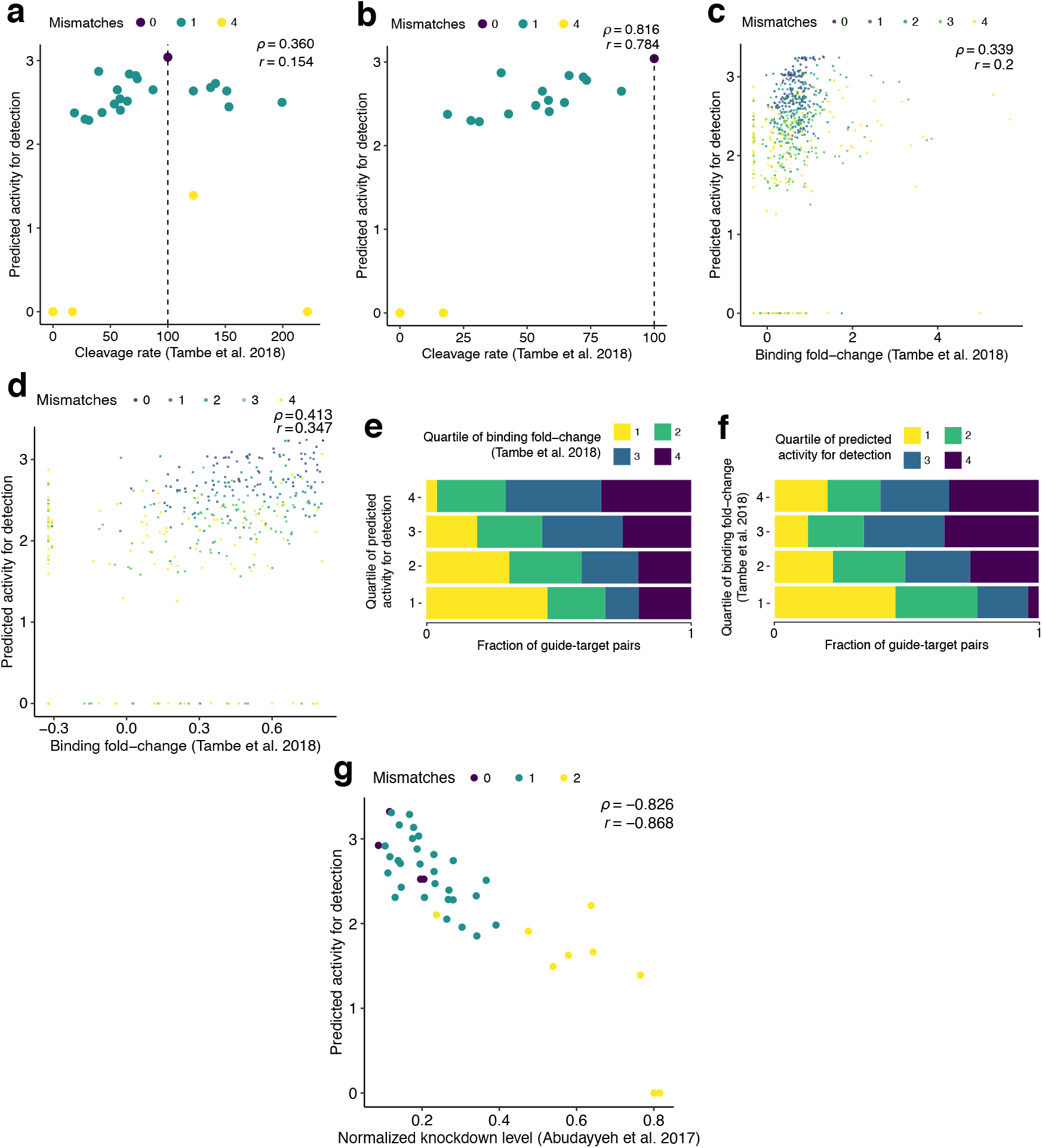
Comparison of predictions with independent Cas13a datasets. **(a)** Each point is a guide-target pair from experiments in ref. 37 measuring LbuCas13a nuclease activity. Horizontal axis is the measured, normalized percent cleavage rate relative to no mismatches and dashed line at 100 shows the value for no mismatches. Vertical axis is our model’s predicted activity for LwaCas13a collateral cleavage. Colors indicate the number of mismatches in the guide-target pair. *ρ*, Spearman correlation; *r*, Pearson correlation coefficient. **(b)** Same as (a), but only for points where mismatches decrease the LbuCas13a cleavage rate—that is, points to the left of the dashed line. Considering this subset helps to mitigate differences arising from LbuCas13a’s higher overall collateral activity compared to LwaCas13a. **(c)** Each point is a guide-target pair from experiments in ref. 37 measuring LbuCas13a–RNA binding affinity. Horizontal axis is the measured, regularized fold-change enrichment for binding to a target. Vertical axis is our model’s predicted activity for LwaCas13a collateral cleavage. **(d)** Same as (c), but only for points where mismatches decrease the binding affinity compared to no mismatches. **(e)** Same data as (c). Each row contains one quartile of pairs divided by our model’s predicted activity (top row, 4, is predicted most active). Colors in each bar indicate the fraction of pairs belonging to each quartile of the binding affinity measurements (4 is highest binding). **(f)** Same data as (c). Each row contains one quartile of pairs divided by binding affinity measurements (top row, 4, is highest binding). Colors in each bar indicate the fraction of pairs belonging to each quartile of the predicted activities (4 is predicted most active). We do not expect a strong correlation in (c–f), but results are consistent with binding being necessary, though not sufficient, for collateral activity. **(g)** Each point is a guide-target pair from experiments in ref. 36 measuring knockdown levels from LwaCas13a on-target (*cis*-) cleavage. Horizontal axis is the measured knockdown level. In the normalized measurements, the non-targeting guide was set to a knockdown level of 1. Vertical axis is our model’s predicted activity.

**Supplementary Figure 16.**
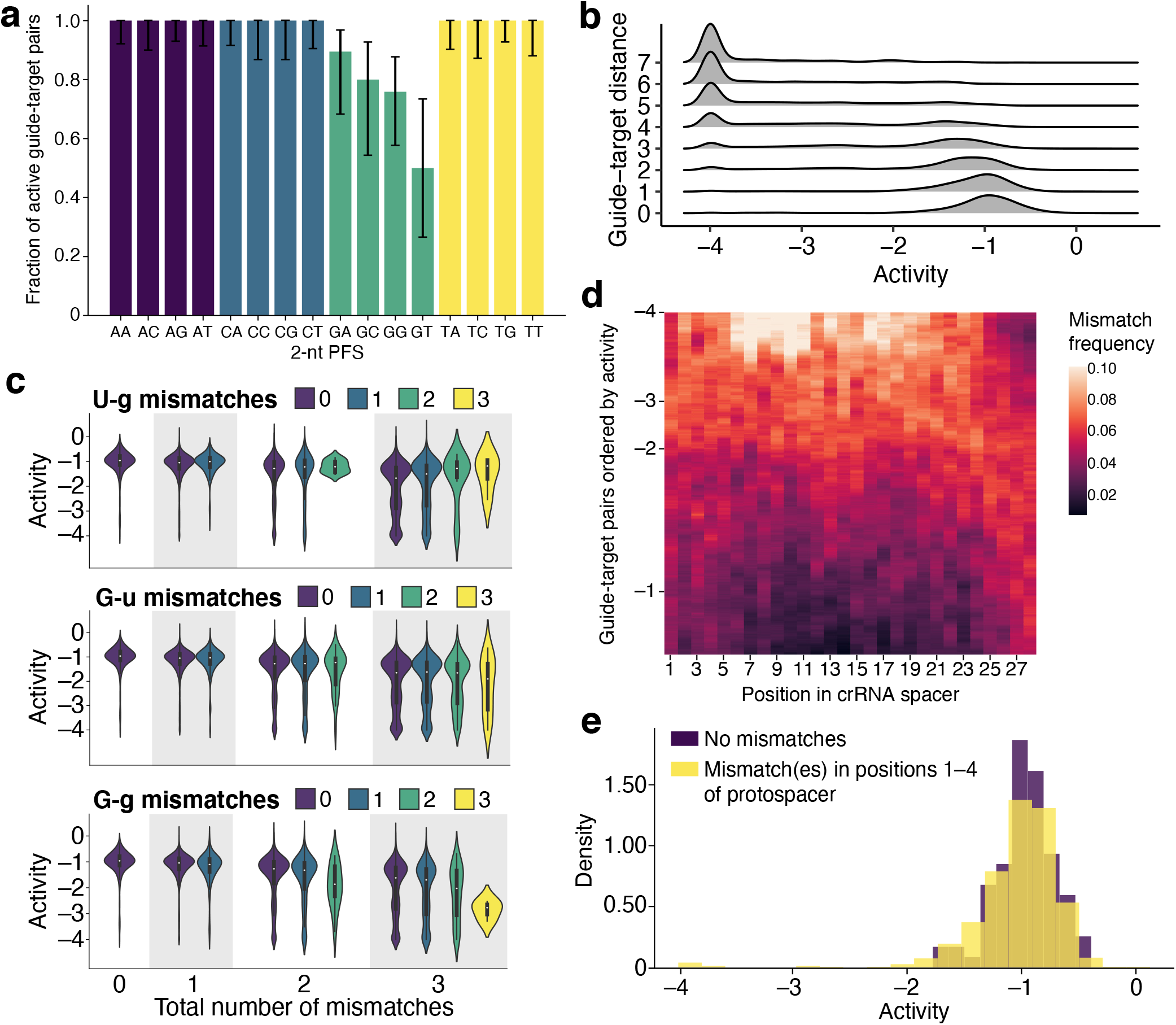
CRISPR-Cas13a guide-target activity. **(a)** Fraction of guide-target pairs that are active for each 2-nt protospacer flanking site allele (PFS; i.e., the canonical Cas13a PFS together with the nucleotide adjacent on the 3’ side of the protospacer). This analysis considers only matching guide-target pairs (i.e., no mismatches) and determines a pair to be active if the median log(*k*) value across replicates is > −2. Error bars represent 95% exact binomial confidence intervals. **(b)** Density of activity for different numbers of mismatches between guides and targets. Here, the number of mismatches is equivalent to Hamming distance. **(c)** Effect of G-U pairing on activity. Top panel highlights U in the target and `G` in the crRNA spacer (U-g). Horizontal groupings (0, 1, 2, 3) are the total number of mismatches in guide-target pairs and the distributions in each grouping separate the pairs by the number of U-g mismatches, showing the density and interquartile ranges of activity; the yellow distribution shows pairs with 3 mismatches, all of which are U-g. Middle panel highlights G-u mismatches and bottom panel, as a benchmark, shows G-g. **(d)** Profile of mismatches among guide-target pairs with similar activity. Each row in the heatmap represents a guide-target pair, ordered by activity, with those having the least activity on top; values on the left indicate activity. For each row *y*, we consider the 1,000 guide-target pairs with activity closest to the pair represented by *y*. Then, at each position *x* in the crRNA spacer sequence, we consider all mismatches at x across our dataset and calculate the fraction of them to which the 1,000 guide-target pairs, centered at *y*, contribute. We plot this fraction; higher values at a row indicate a preponderance of mismatches among the guide-target pairs with the activity represented by that row. **(e)** Density of guide-target pairs that have no mismatches (purple) compared to those that have at least one mismatch in the first four positions of the protospacer and no mismatch elsewhere (yellow). As in (b), here a G-U pair is counted as a mismatch. Positions in (d) are relative to the crRNA spacer sequence, while positions in (e) (and elsewhere) are relative to the target. In our analyses we set a lower threshold of −4 on the activity owing to measurement limitations (see Supplementary Fig. 4b and Methods for details), so in (b–e) guide-target pairs shown at −4 include pairs with true activity below this threshold; in (b) and (c), densities drop slightly below −4 owing to smoothing.

**Supplementary Figure 17.**
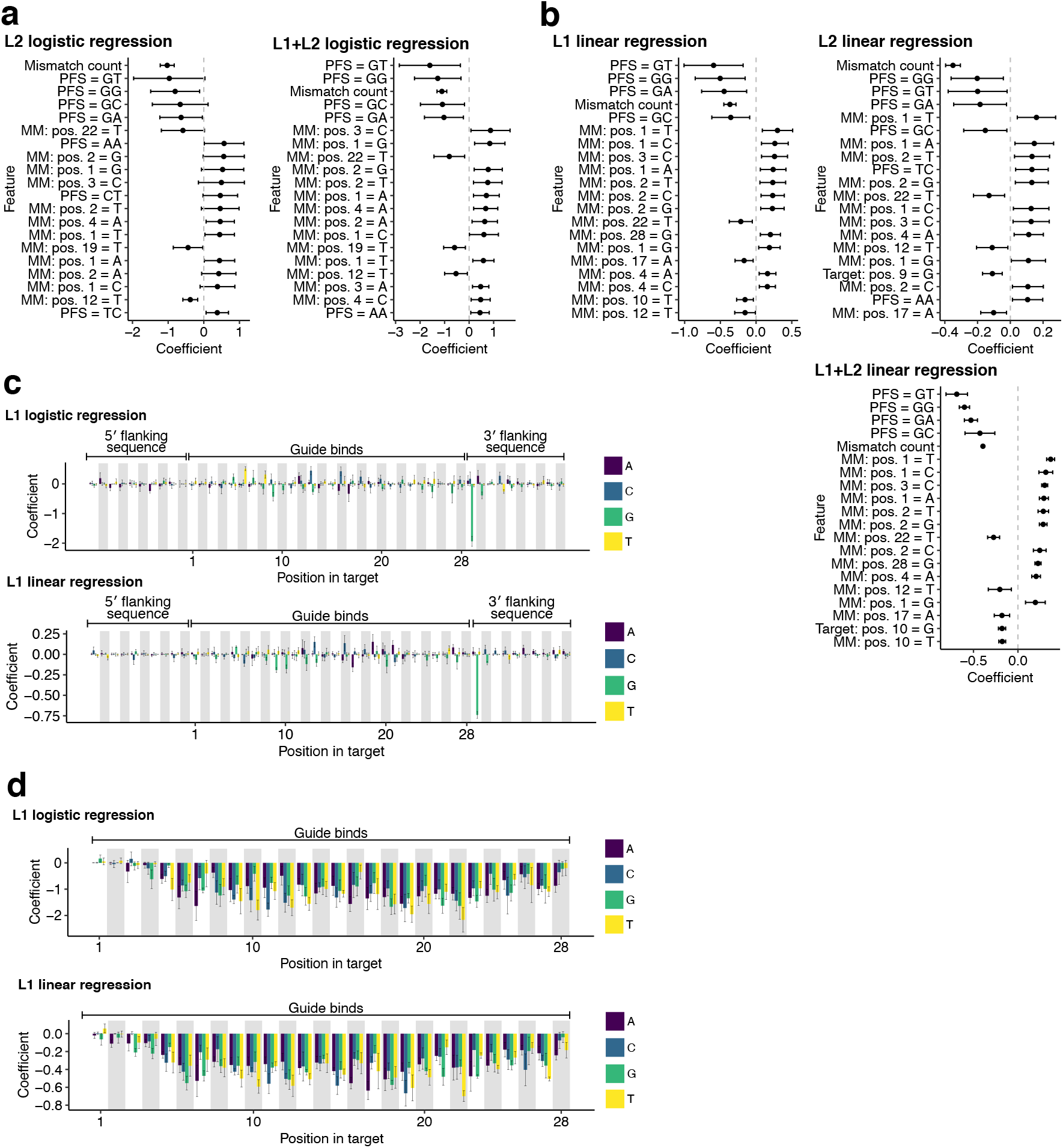
Importance of features in linear models. Feature coefficients in linear models for predicting guide activity. In all panels, plotted value is the mean of the coefficient across training on five folds and error bar is the 95% confidence interval. Positions are along the target, where the guide binds at positions 1–28: position 1 is the 5’ end of the protospacer (position 28 of the guide spacer); position 28 is the 3’ end of the protospacer (position 1 of the guide spacer); positions `−9-0` are 10-nt of context flanking the protospacer on the 5’ end; positions 29-38 are 10-nt of context flanking the protospacer on the 3’ end. **(a)** Linear models for classifying guide-target activity. Coefficients are ranked by absolute value and the top 20 are shown. Models use the ‘One-hot MM + Handcrafted’ input, which combines one-hot encoding of target sequence nucleotides and of mismatches in guides relative to the target with curated features of hypothesized importance (details in Methods). PFS, protospacer flanking site (flanking on 3’ side) including nucleotides at two positions. ‘MM:’ indicates a mismatch at the given position with the base representing the complement of the nucleotide in the guide’s spacer. ‘Target:’ indicates a base in the target sequence, matching with the guide, at the given position. L1 logistic regression is in Fig. 2g. **(b)** Same as (a) but for regression models on active guide-target pairs. **(c)** Coefficients for nucleotide composition of the target, from the L1 logistic regression model used for classifying activity and the L1 linear regression model used on active guide-target pairs. Models use the ‘One-hot MM’ input, which combines one-hot encoding of target sequence nucleotides and of mismatches in guides relative to the target (it leaves out the handcrafted features, including number of mismatches and two-nucleotide PFS interaction, present in (a) and (b) so that they do not affect the coefficients along the target). Colors represent nucleotides in the target sequence. Outlier at position 29 indicates the effect of a `G` PFS. **(d)** Same model and input as in (c), but for the features representing guide-target mismatches along the target sequence in the region to which the spacer binds. Colors represent the complement of the nucleotide in the guide’s spacer; for example, `A` indicates `T` in the spacer sequence that is mismatched with either `C, G`, or `T` in the target sequence.

**Supplementary Figure 18.**
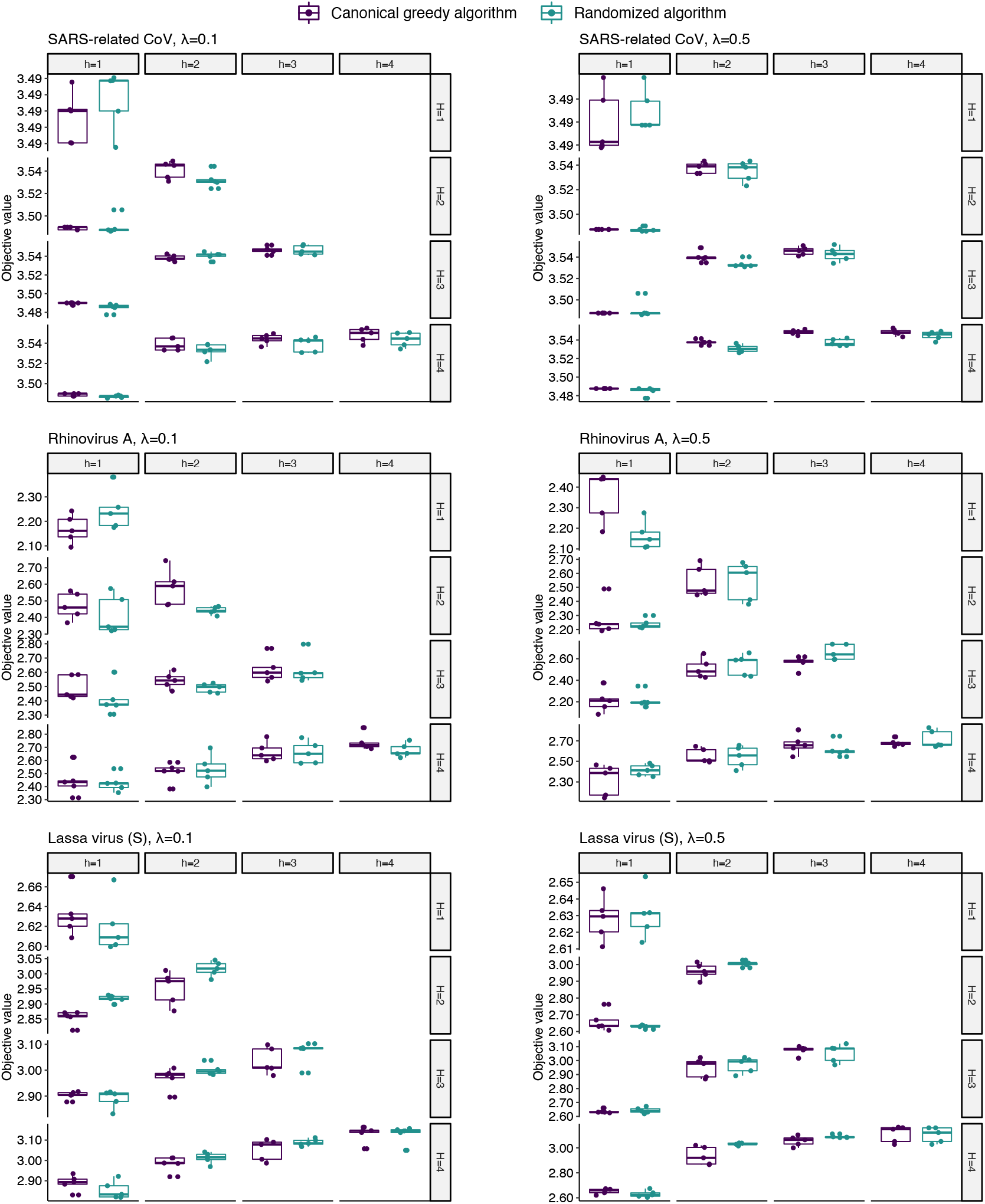
Comparison of algorithms for submodular maximization. Objective values of the optimal solutions identified by two algorithms for submodular maximization: the canonical greedy algorithm for monotone functions^43^ and a randomized algorithm with provable guarantees on non-monotone functions^42^. The function here is non-monotone, but neither algorithm clearly outperforms the other. Three viral species are shown. Each was evaluated for two choices of the weight on the soft constraint/penalty, indicated by λ, as well as different choices of the soft cardinality constraint (h) and hard constraint (*H*) with *h* ≤ *H*. Supplementary Note 1 contains a definition of the objective function, including the penalty weight and cardinality constraints. Each point indicates the result of one of 5 runs; differences account for randomness both in the randomized greedy algorithm and in constructing the ground set.

**Supplementary Figure 19.**
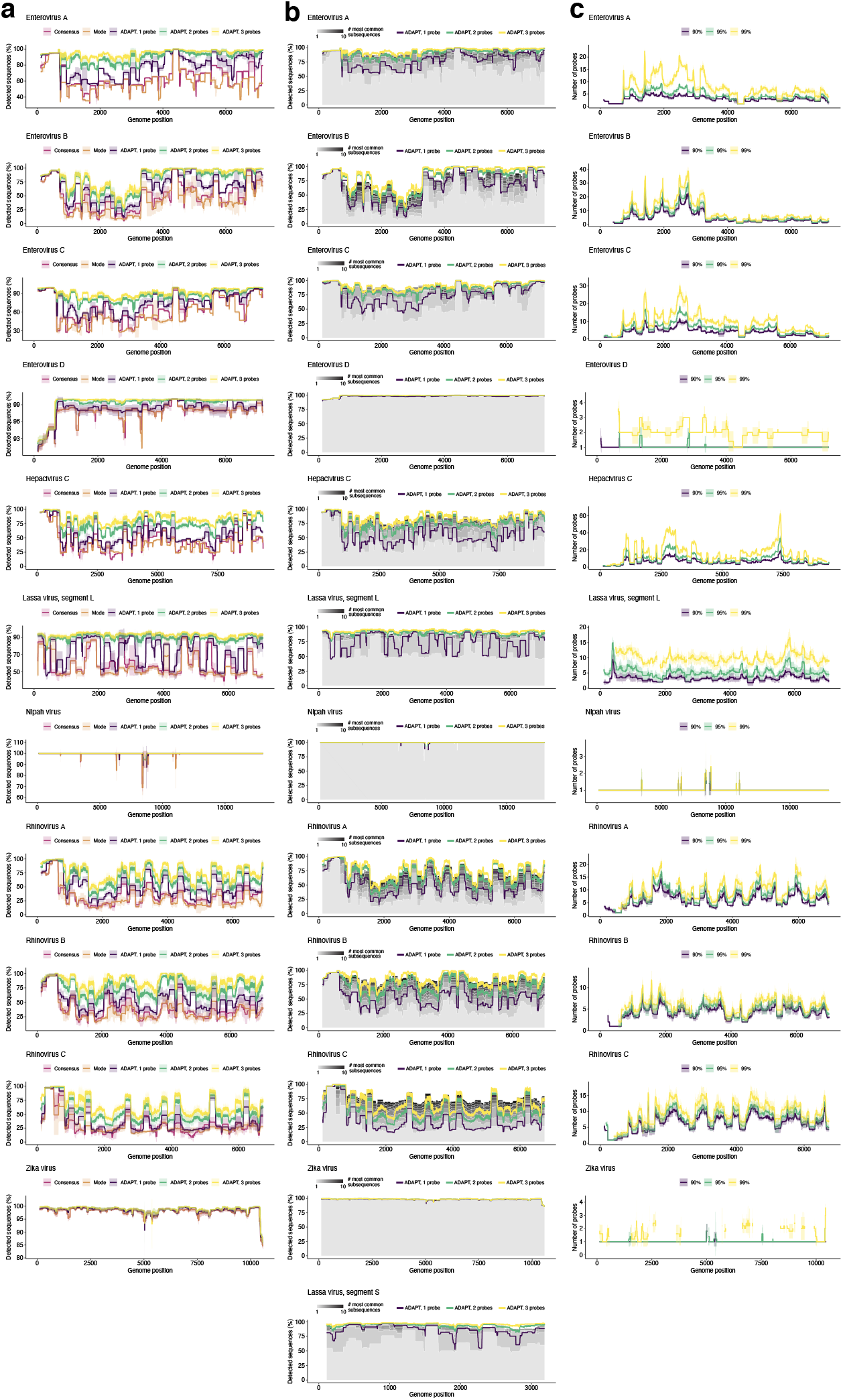
Comprehensiveness of probe design. Comparison of ADAPT’s comprehensiveness in designing probe sequences to baseline methods, across 11 viral species. **(a)** Fraction of genome sequences detected, with different design strategies in a 200 nt sliding window, using a model in which 30 nt probes detect a target if they are within 1 mismatch, counting G-U pairs at matches. ‘Consensus’, probe-length consensus subsequence from the window that detects the most number of genomes; ‘Mode’, most common probe-length subsequence within the window. Our approach (ADAPT) uses hard constraints of 1–3 probes and maximizes activity. **(b)** Same as (a), but generalizing the ‘Mode’ beyond one probe. Stacked grays show the cumulative fraction of genome sequences detected using the *n* probes representing the n most common subsequences at a site, ranging n from 1 (lightest gray) to 10 (darkest gray). Top of the lightest gray area corresponds to ‘Mode’ in (a) (1 probe) and top of all the grays is using 10 probes. **(c)** Number of probes identified by ADAPT when solving a dual objective: minimizing the number of probes to detect >90%, >95%, and >99% of LASV genomes using the model in (a). Gaps at a site are present when it is not possible to construct a probe set that reaches the desired coverage, owing to gaps or missing data. In (a) and (c), lines show the mean and shaded regions around them are 95% pointwise confidence bands across genomes sampled for each virus calculated by bootstrapping, i.e., randomly sampling genomes to be input to the design process; (b) shows only the mean, to ease visualization. Figure 3b,c show results from panel (a) and (c) for Lassa virus, segment S.

**Supplementary Figure 20.**
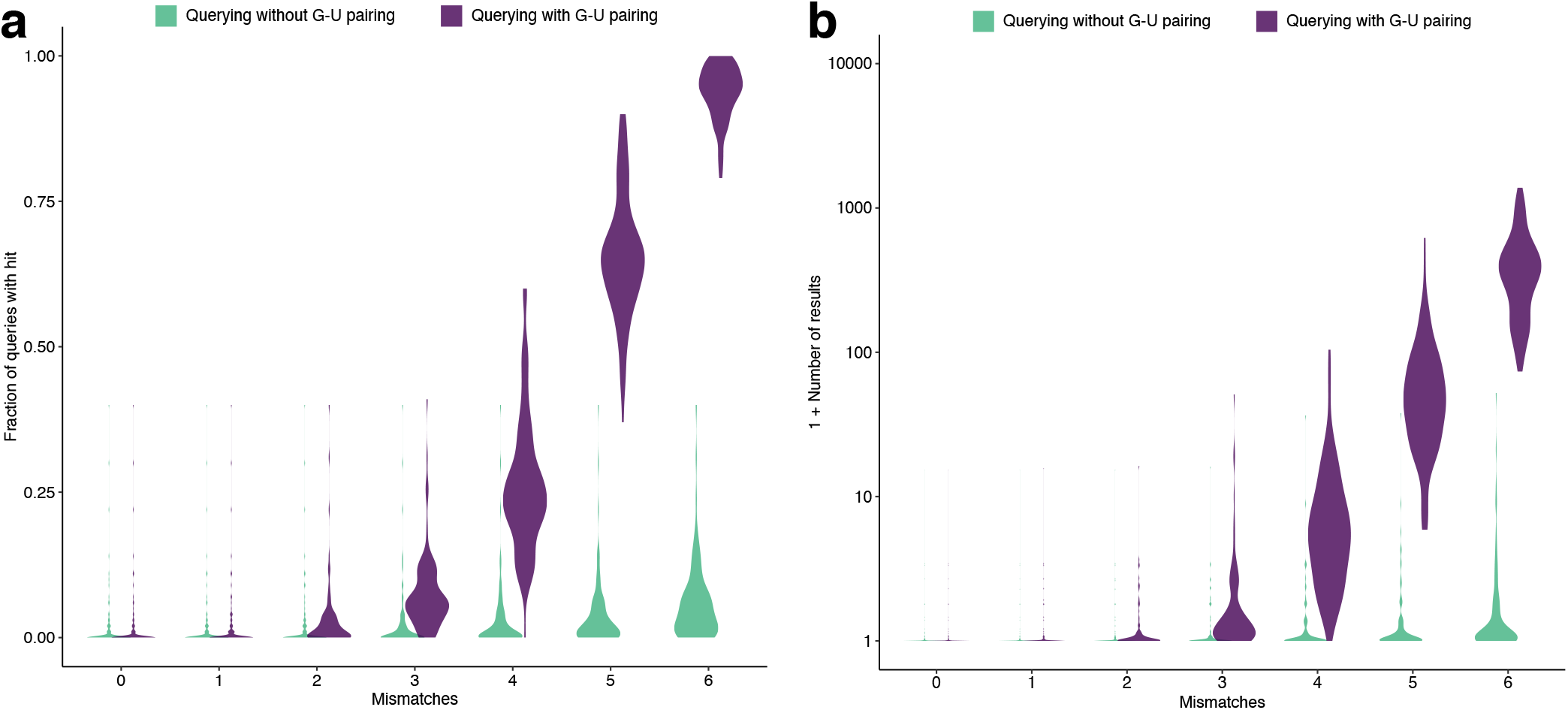
Potential hits with tolerance of G-U base pairing. Being tolerant of G-U base pairing increases the potential for non-specific hits of a *k*-mer. We built an index of ~1 million 28-mers from 570 human-associated viral species. For each of 100 randomly selected species, we queried 28-mers for hits against the other 569 species (details in Methods). We performed this for each choice of *m* mismatches, counting a non-specific hit as one within *m* mismatches of the query, both being sensitive to G-U base pairing (purple; counting it as a match) and not being sensitive to it (green; counting it as a mismatch). Violin plots show the distribution, across the selected species, of the mean of the measured value taken over the queries for each species. **(a)** Fraction of queries that yield a non-specific hit. The measured value for a query is 0 (no hit) or 1 (≥ 1 hit), so the mean represents the fraction of queries with a hit. **(b)** Number of non-specific hits per query.

**Supplementary Figure 21.**
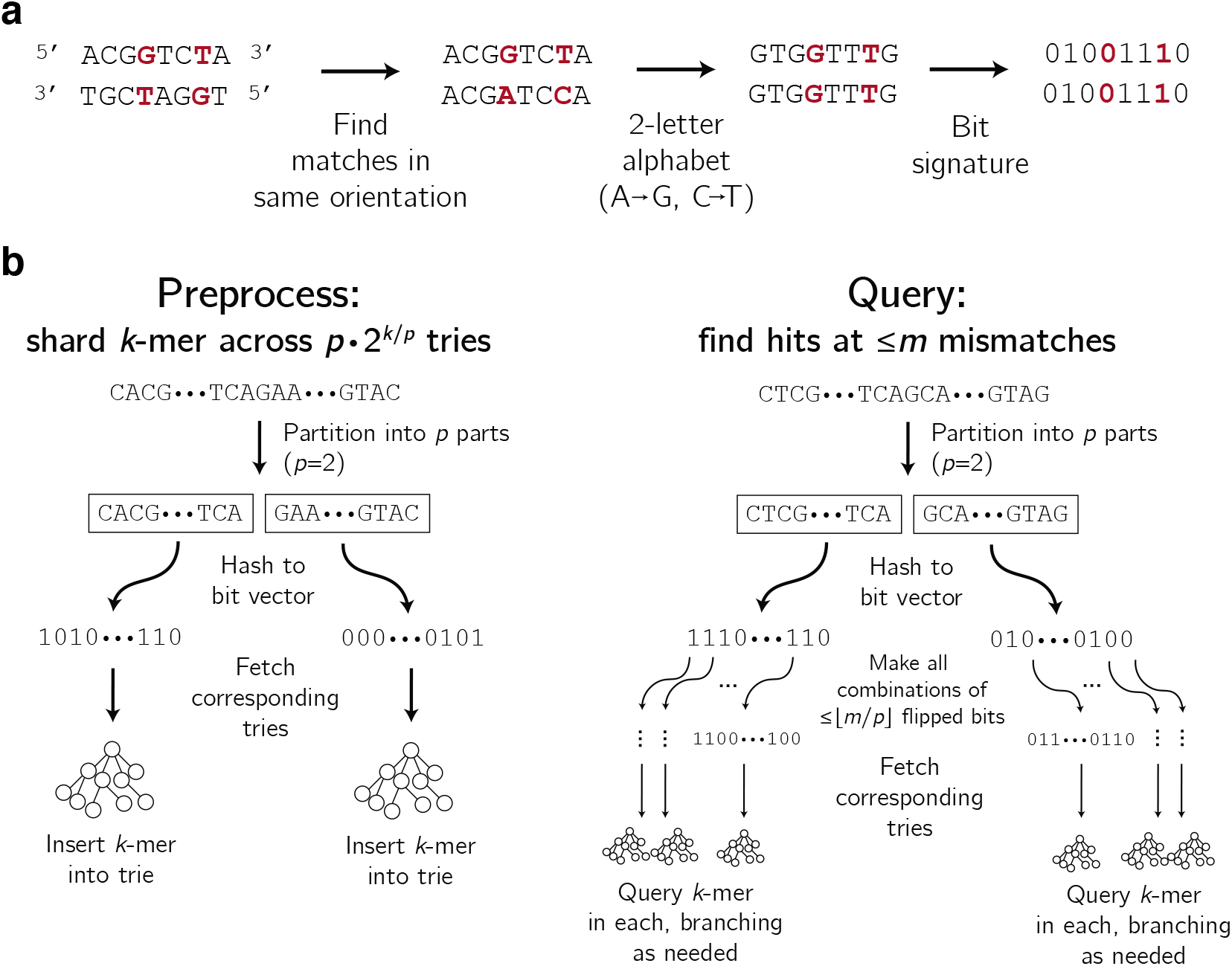
Sharding k-mers across tries for specificity queries. **(a)** Constructing a bit signature after transforming a string to a two-letter alphabet, described in Supplementary Note 2b. Two strings that match up to G-U base pairing (shown here as G-T) have the same bit signature. **(b)** Left: Inserting a *k*-mer into the data structure of tries. Each k-mer is inserted into *p* tries, and there are *p* · 2^*k/p*^ tries in total. Right: Querying a *k*-mer for near neighbors (within m mismatches, sensitive to G-U base pairing as a match).

**Supplementary Figure 22.**
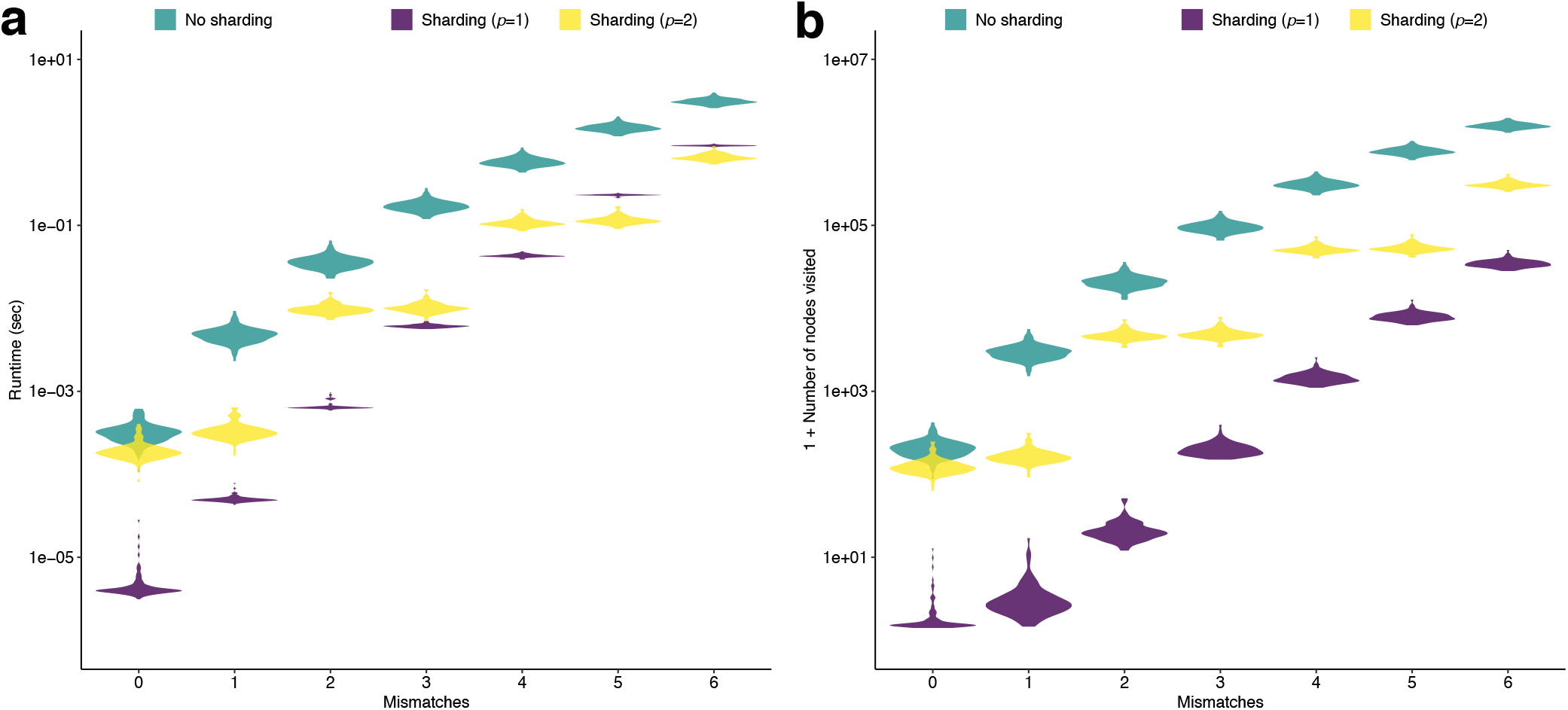
Benchmarking of specificity queries. **(a)** The runtime of querying using an index of ~1 million 28-mers across 570 human-associated viral species. For each of 100 randomly selected species, we queried 28-mers for hits against the other 569 species. Violin plots show the distribution, across the selected species, of the mean runtime for each query. Green shows results on a single, large trie of 28-mers; purple (*p* = 1) and yellow (*p* = 2) show results on the approach described in Supplementary Note 2d, with two choices of the partition number *p*. **(b)** Same as (a), but showing total number of nodes visited across the trie(s). The decrease in this value using our approach suggests that parallelizing the approach—by searching within multiple tries in parallel—may provide a further speedup.

**Supplementary Figure 23.**
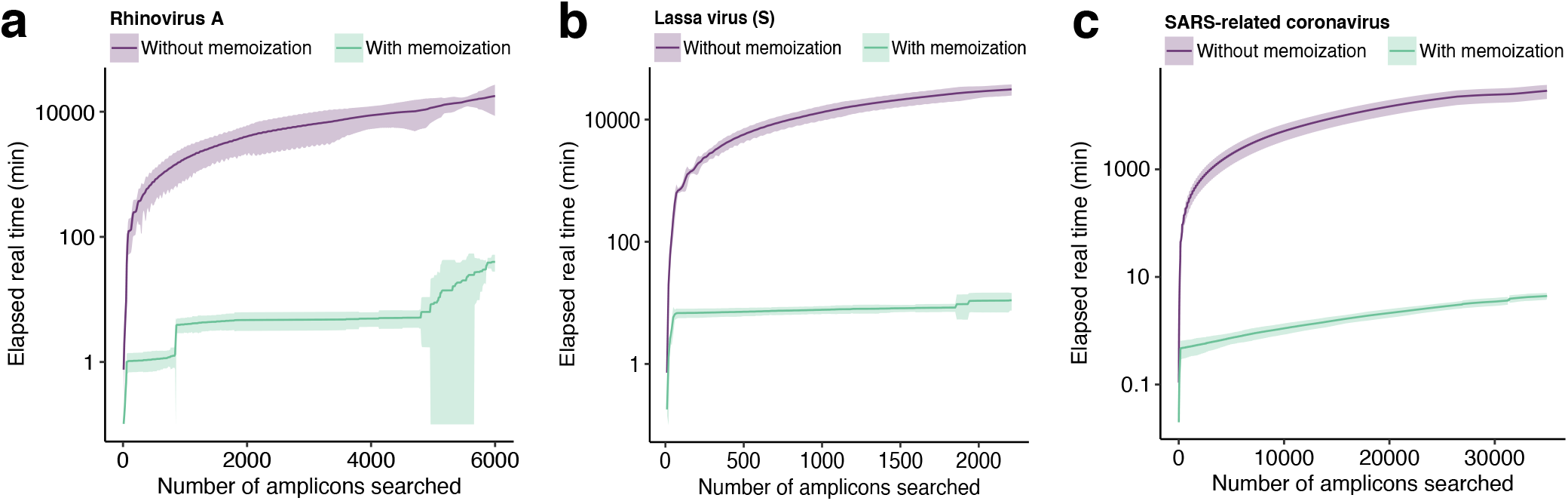
Runtime improvement provided by memoization. Runtime of ADAPT’s search with and without memoizing computations, for three species. We plot the cumulative elapsed real time (minutes) at each successive window (amplicon) that ADAPT considers during its search. Shaded regions indicate a 95% confidence interval calculated across 3 runs (same input) and line is the mean. **(a)** Rhinovirus A. The lower end of the confidence interval is cutoff at 0.1. Memoization provides a 99.71% reduction in runtime (mean). **(b)** Lassa virus, segment S. Memoization provides a 99.75% reduction in runtime (mean). **(c)** SARS-related coronavirus. Memoization provides a 99.96% reduction in runtime (mean). For (b) and (c), the search without memoization was ended before its completion; thus, for these, the reduction is a lower bound assuming a continued faster growth of the runtime without memoization.

**Supplementary Figure 24.**
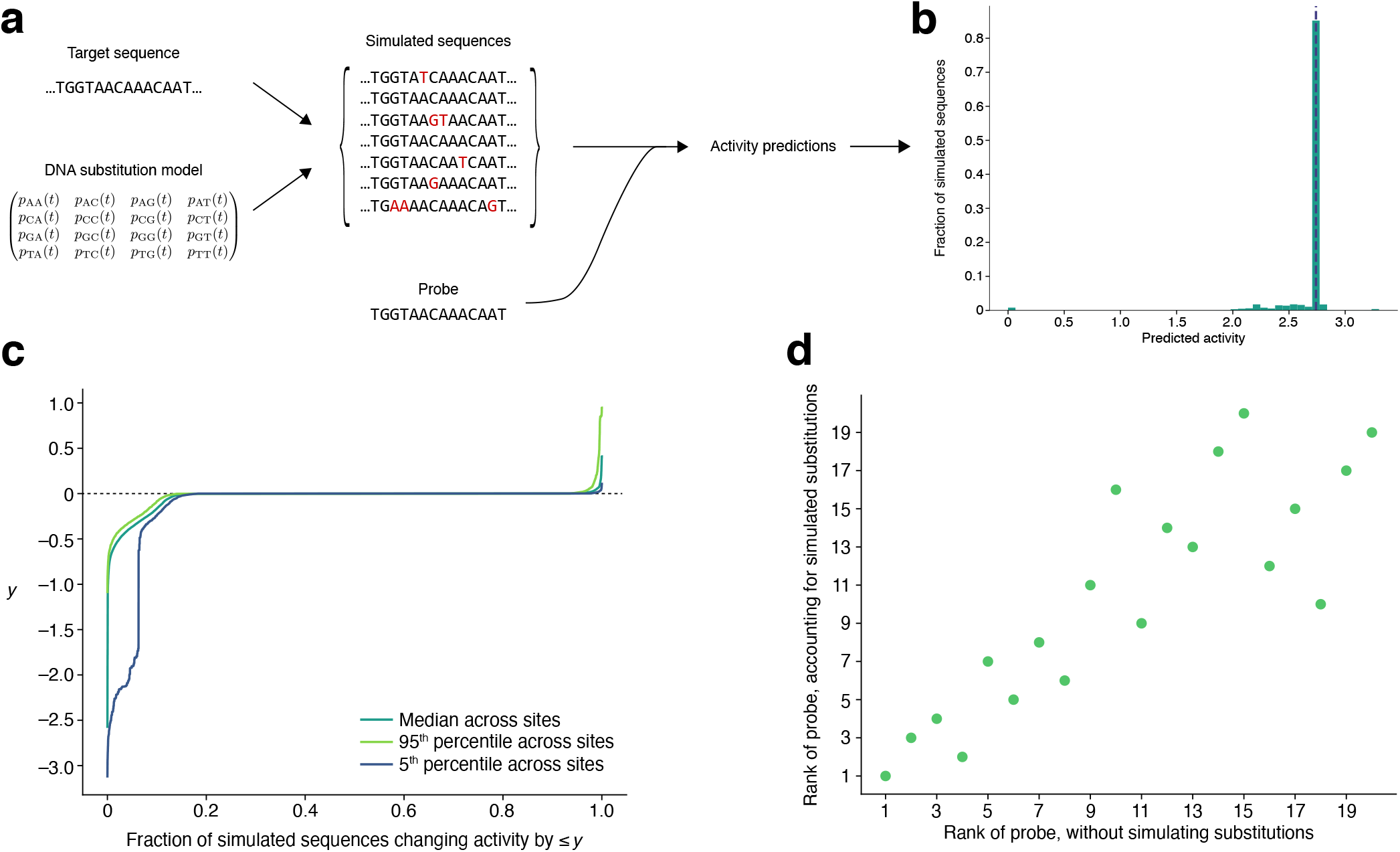
Estimating probe activity under simulated substitutions. **(a)** Sketch of proactive scheme to estimate probe performance, after a period of time, by simulating relatively likely nucleotide substitutions. Starting with a target sequence and a GTR substitution model, we sample from a distribution of substitutions made to the original target sequence. In analyses that follow, we sample after 5 years. Against each simulated sequence, we predict the detection activity of a given probe using our Cas13a predictive model; across the simulated sequences, these predictions provide a distribution of activity under potential substitutions. **(b)** Distribution of predicted detection activity across 10,000 simulated sequences that originate from a target sequence at one site in the SARS-CoV-2 genome. The probe is complementary to the original target sequence, except we randomly introduce a single mismatch. Most simulated sequences do not introduce any substitutions (i.e., they are identical to the original); the peak in the histogram (and vertical dashed line) represents these ones. Other than these simulated sequences, most simulated substitutions degrade the activity of the probe (left of the dashed line). Some enhance its activity (right of the dashed line), for example, by reversing the existing mismatch. **(c)** Inverse CDF of the change in predicted detection activity after simulating substitutions, summarized across 1,000 random sites in the SARS-CoV-2 genome; (b) showed one such site. At each of these 1,000 sites, we simulate 10,000 target sequences according to our substitution model and construct a distribution of the change in the probe’s predicted detection activity compared to its activity in detecting the original sequence. As in (b), at each site the probe is complementary to the original target sequence, except with one random mismatch. Plotted is the median change taken across sites, as well as the 95^th^ and 5^th^ percentiles. The faster the curve rises to 0, the less likely there is to be degraded activity. That the 5^th^ percentile curve shows a sharp drop for low values (~0.1) on the horizontal axis indicates that some sites may experience a pronounced degradation in detection activity over time, but that even for these sites it is unlikely (~10% chance). **(d)** Effect of simulating substitutions on the ordering of ADAPT’s designs. We begin with the top 20 design options output by ADAPT for targeting SARS-CoV-2 genomes and, for this analysis, consider only the probe (Cas13a guide) from each design option. Each point represents one of the 20 probes. We rank the probes according to their mean predicted detection activity across the genomes; this ranking is on the horizontal axis. Then, for each genome, we simulate 10,000 sequences according to our substitution model (at the site where a probe binds) and compute the 5^th^ percentile of the predicted detection activities between the probe and these simulated sequences. We rank the probes accounting for simulated substitutions (vertical axis) according to the mean of this 5^th^ percentile value taken across the genomes. In this analysis, we use only 500 randomly sampled genomes from the set of genomes used to design the 20 probes with ADAPT, in order to reduce runtime.

**Supplementary Figure 25.**
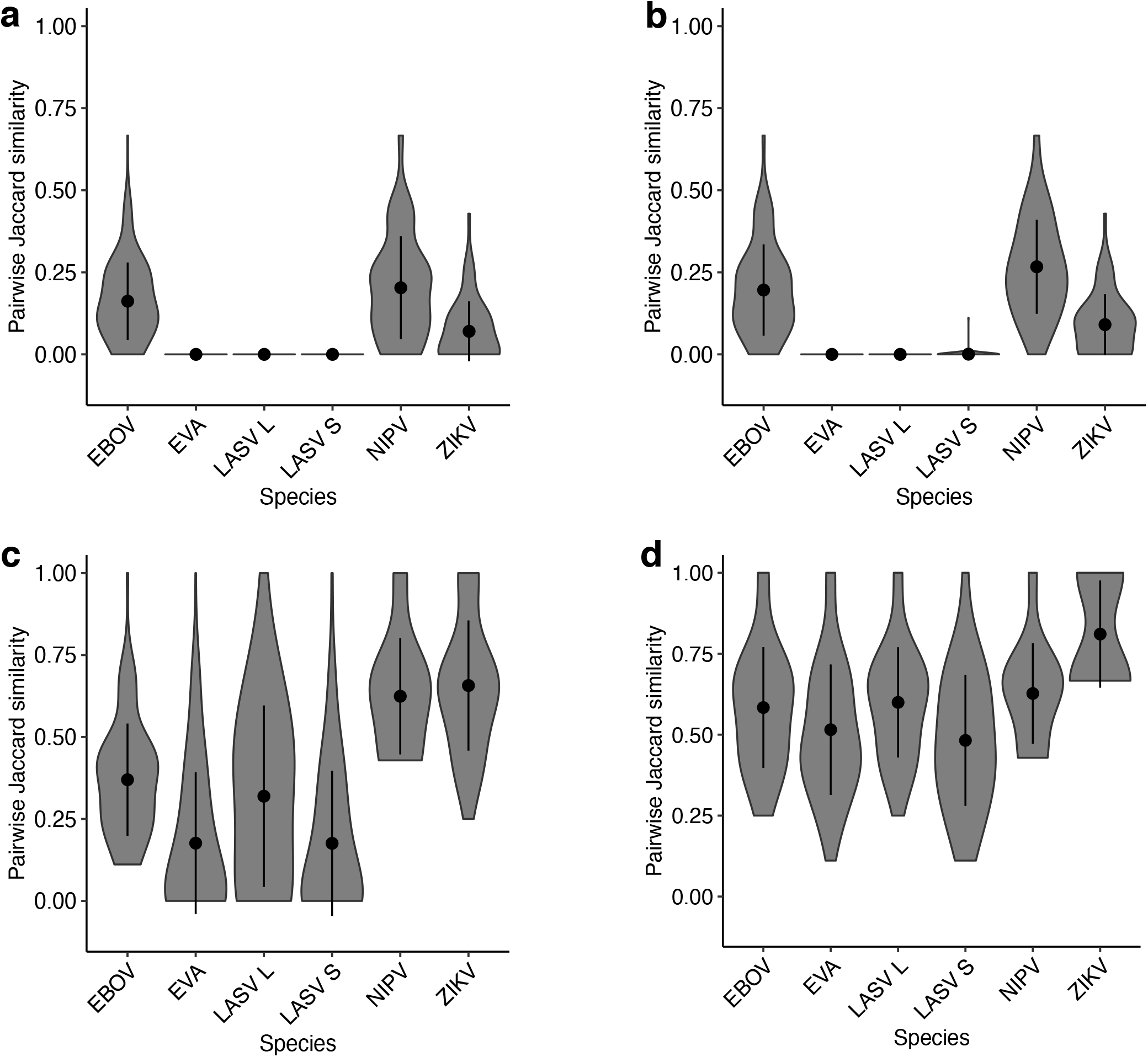
Dispersion in ADAPT’s designs. For each species, we ran ADAPT 20 times. For each pair of runs, we calculated the Jaccard similarity comparing the top 5 design options from each. Violin plots show a smoothed density estimate of the pairwise Jaccard similarities; dot indicates the mean and bars show 1 standard deviation around the mean. Lower values indicate more variability in ADAPT’s design outputs across runs. The panels show different methods of providing input genomes and of comparing a pair of design outputs. **(a)** Using resampled input genomes for each run and considering two design options to be equal if they have exactly the same primers and probes. **(b)** Using the same input genomes for each run and considering two design options to be equal if they have exactly the same primers and probes. **(c)** Using resampled input genomes for each run and considering two design options to be equal if their endpoints are within 40 nt of each other. **(d)** Using the same input genomes for each run and considering two design options to be equal if their endpoints are within 40 nt of each other. When using resampled input genomes, the comparisons account for algorithmic randomness and input sampling. When using the same input genomes, the comparisons account only for algorithmic randomness. EBOV, Zaire ebolavirus; EVA, Enterovirus A; LASV L/S, Lassa virus segment L/S; NIPV, Nipah virus; ZIKV, Zika virus.

**Supplementary Figure 26.**
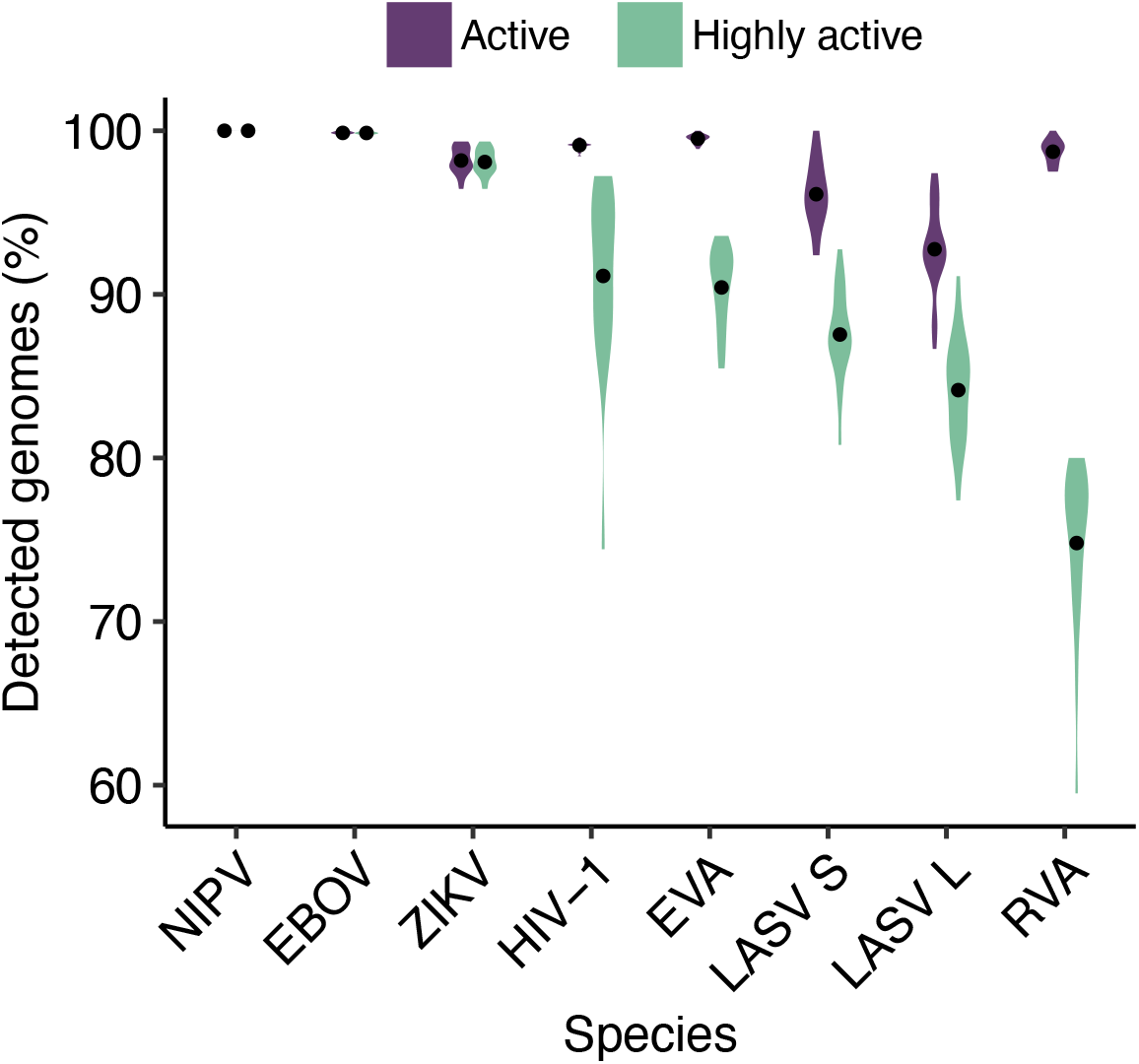
Cross-validated evaluation of detection with relaxed design parameters. Crossvalidated evaluation of detection using more relaxed design parameters than the choices in Fig. 4b; the relaxed parameters (Methods) tolerate more complex assay designs (e.g., more guides) to achieve higher sensitivity. For each species, we ran ADAPT on 80% of available genomes and estimated performance, averaged over the top 5 design options, on the remaining 20%. Distributions are across 20 random splits and dots indicate mean. Purple, fraction of genomes detected by primers and for which Cas13a guides are classified as active. Green, same except Cas13a guides also have regressed activity in the top 25% of our dataset. NIPV, Nipah virus; EBOV, Zaire ebolavirus; ZIKV, Zika virus; LASV S/L, Lassa virus segment S/L; EVA, Enterovirus A; RVA, Rhinovirus A.

**Supplementary Figure 27.**
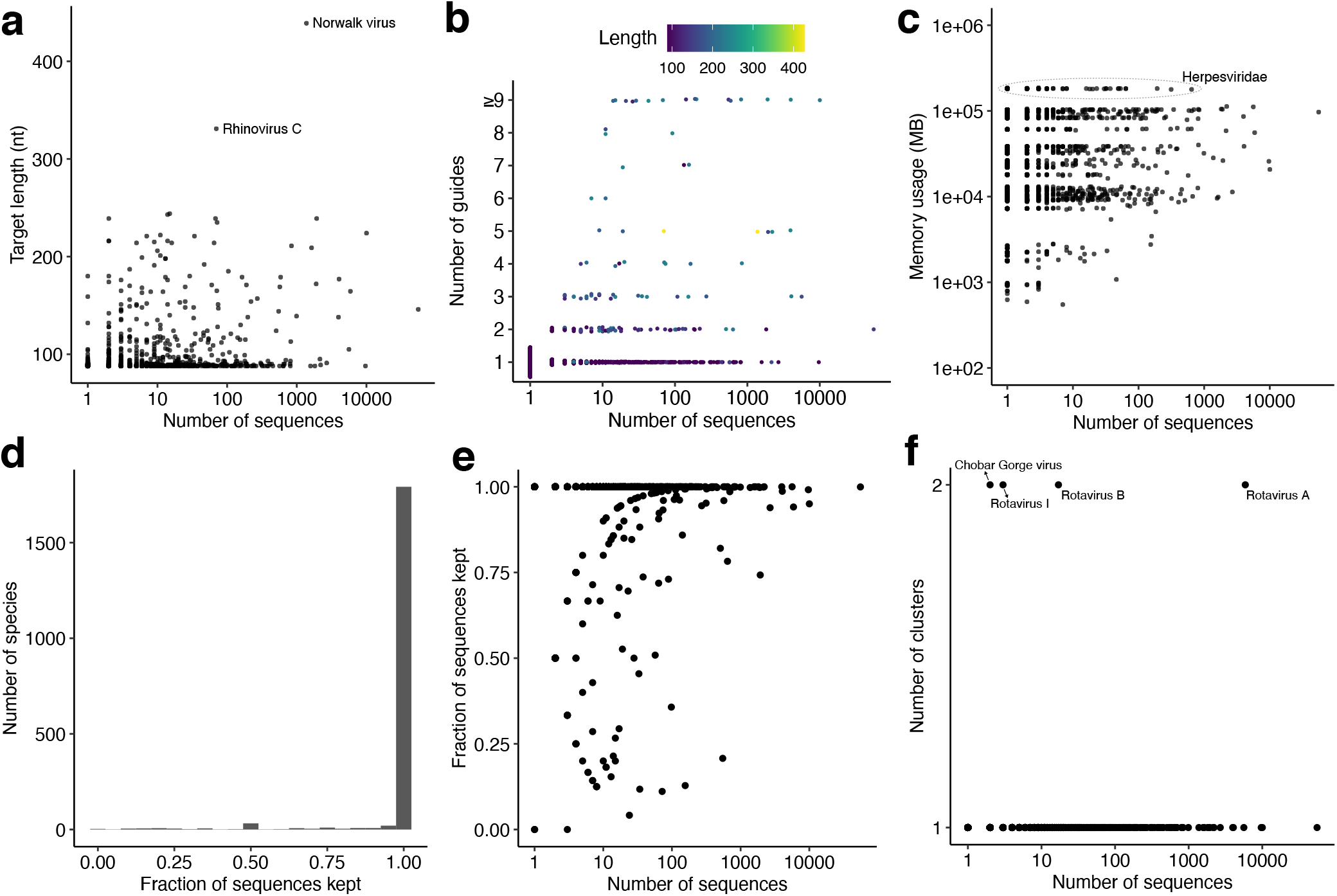
Results of ADAPT’s designs for 1,926 vertebrate-infecting viruses. Running ADAPT on 1,933 vertebrate-infecting species produced designs on 1,926 (Methods). **(a)** Length of each target region, i.e., amplicon, in nt of the highest-ranked design output by ADAPT for each species. As part of the design we restricted the length to ≤ 250-nt for all species except two (Methods). Horizontal axis is the number of input sequences for design. **(b)** Number of Cas13a guides in the highest-ranked design option for each species, produced using the objective function in which we minimize the number of guides subject to detecting > 98% of sequences with high activity. Color indicates the length of the targeted region (amplicon) in the design. **(c)** Maximum resident set size (RSS), in MB, of the process running ADAPT on each species. **(d)** Distribution, across species, of the fraction of input sequences passing curation. **(e)** Fraction of input sequences passing curation for each species compared the number of input sequences for that species. **(f)** Number of clusters for each species compared to the number of input sequences for that species. In (a–c) and (e–f), each point is a species.

**Supplementary Figure 28.**
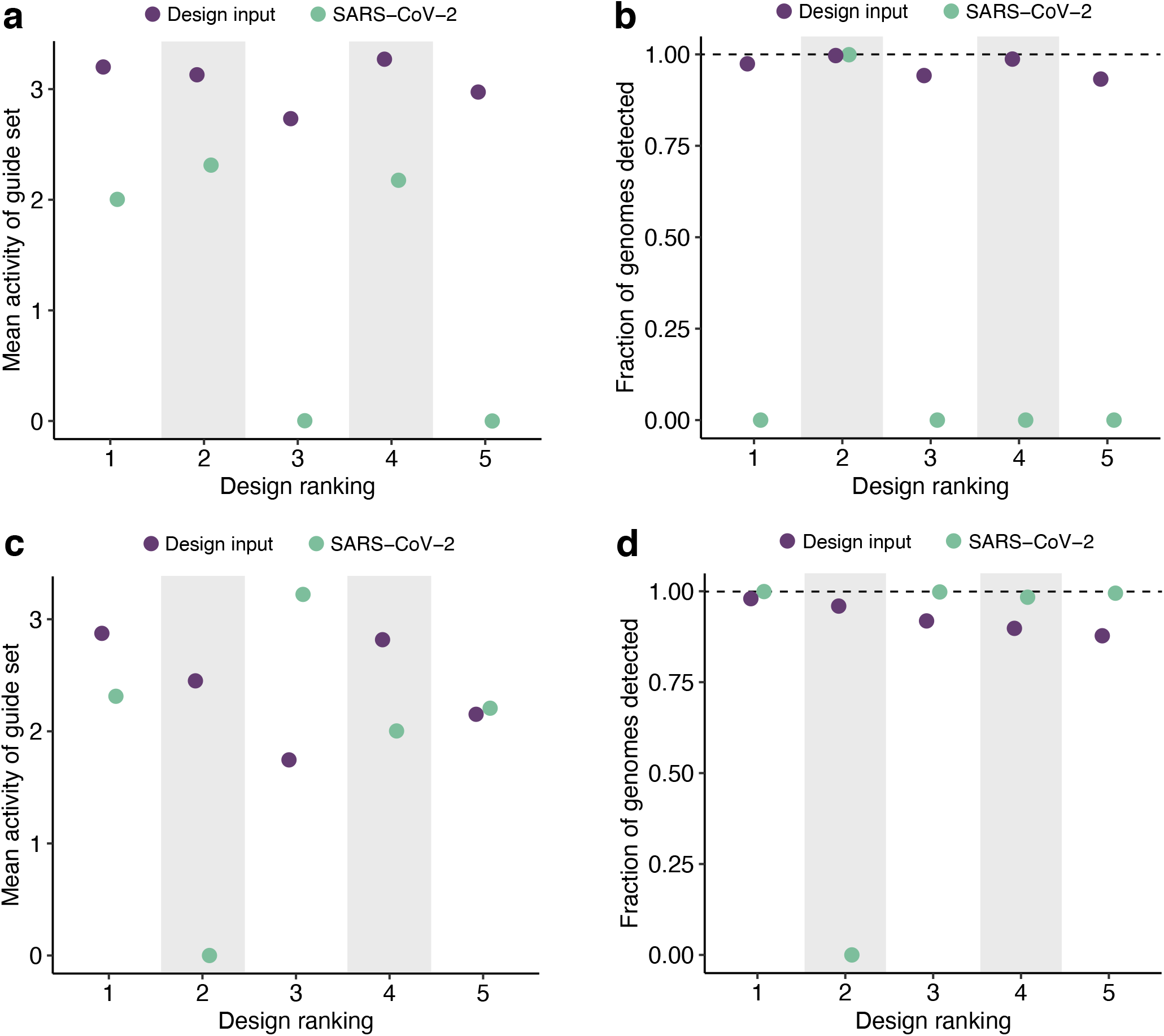
Evaluation of SARS-CoV-2 detection with assays designed, using genomes through 2018, to detect SARS-related coronavirus. The SARS-related CoV designs were generated using genomes available through the end of 2018, which simulates the design of broadly-effective assays a year before SARS-CoV-2’s emergence. SARS-related CoV is a species that encompasses SARS-CoV-2, as well as SARS-CoV-1 and viruses sampled from animals. **(a)** Performance of Cas13a guides from each of the five highest-ranked design outputs from ADAPT (ordered by ranking; 1 is best). Points indicate the mean predicted activity of each design’s guides in detecting targeted genomes. Purple, mean across the 311 genomes used for the design (all SARS-related CoV genomes through the end of 2018). Green, mean across the 184,197 SARS-CoV-2 genomes available through November 12, 2020. **(b)** Fraction of genomes predicted to be detected by each design’s assay, accounting for both the primers and guides in the assay (details in Methods). Designs were produced as in (a) and colors are as in (a). **(c)** Same as (a), except the designs used input that downsampled SARS-CoV-1 to a single genome, effectively down-weighing consideration to SARS-CoV-1 in the design. Purple, mean across the 49 genomes used for the design. Green, mean across the 184,197 SARS-CoV-2 genomes available through November 12, 2020. **(d)** Fraction of genomes predicted to be detected by each design’s assay, accounting for both the primers and guides in the assay. Designs were produced as in (c) and colors are as in (c). In (a) and (c), values at 0 indicate Cas13a guides that are classified to be inactive; values above 0 are classified to be active.

**Supplementary Figure 29.**
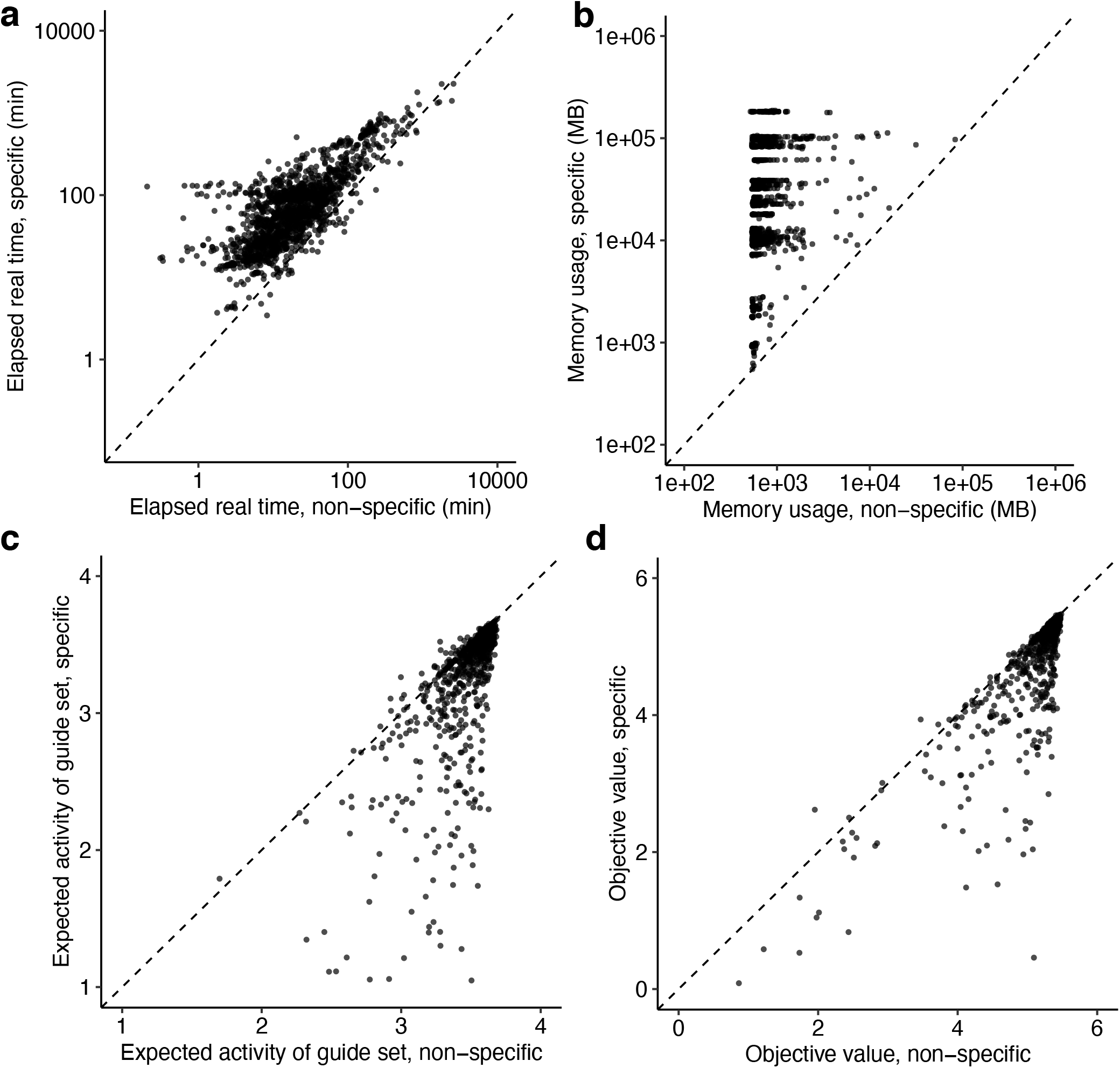
Effects of enforcing specificity on ADAPT’s designs for 1,926 vertebrate-infecting viruses. In each panel, each point is a species and comparisons are with and without enforcing species-level specificity within each family, **(a)** End-to-end elapsed real time running ADAPT. **(b)** Maximum resident set size (RSS), in MB, of the process running ADAPT. **(c)** Mean activity of the guide set, from the highest-ranked design option, across input sequences. **(d)** Objective value of the highest-ranked design option, which incorporates expected activity of the guide set, the number of primers, and the target region length. Not shown, 9 species with objective value < 0. In all panels, 1,926 species are shown (7 of the 1,933 vertebrate-infecting species did not produce designs; Methods).

**Supplementary Figure 30.**
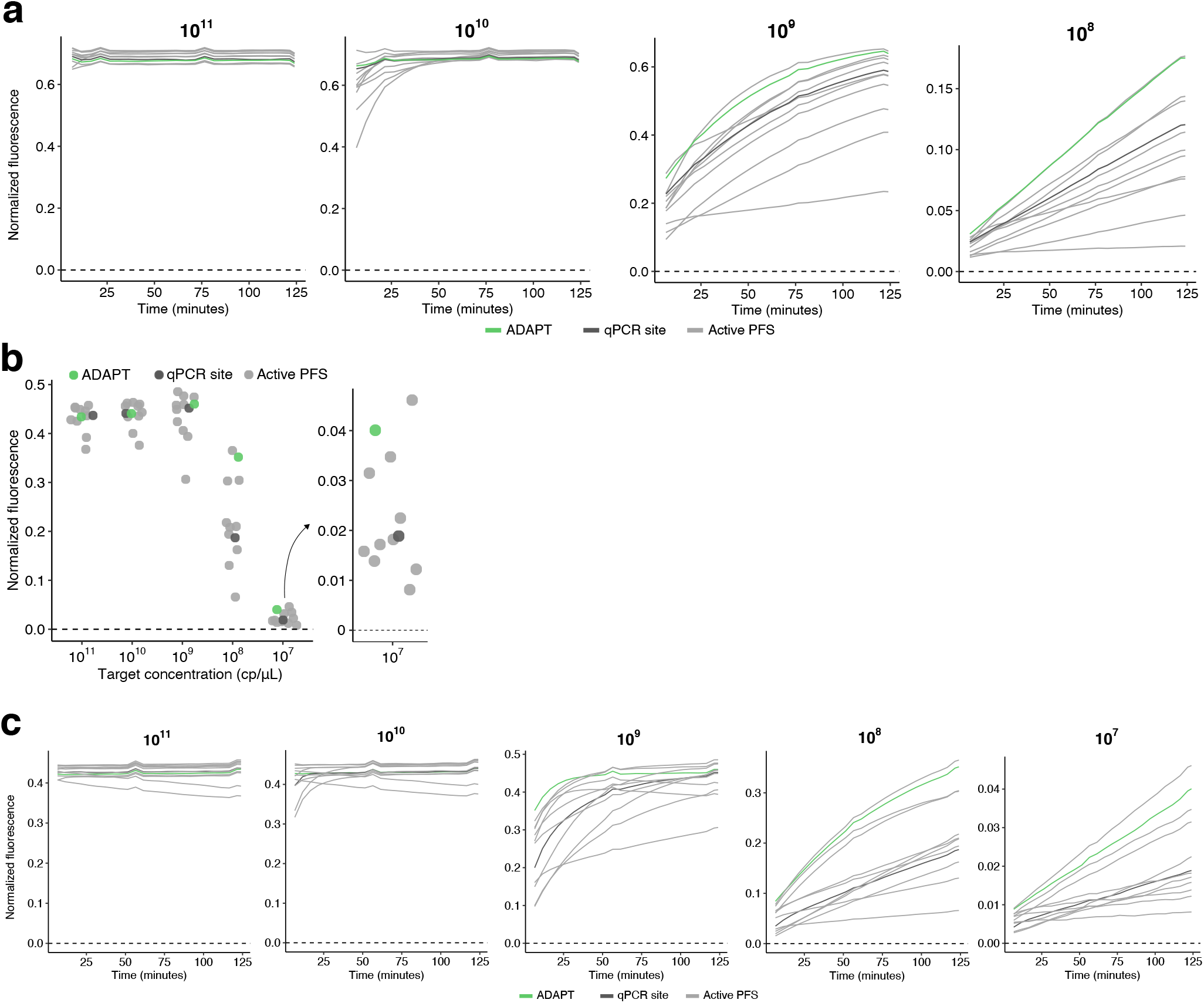
Benchmarking ADAPT’s designs in US CDC SARS-CoV-2 amplicons. **(a)** Fluorescence over time for Cas13 guides at varying target concentrations (top of each plot, in cp/μL) within the US CDC’s SARS-CoV-2 N1 RT-qPCR amplicon. Compared guides are ADAPT’s design (green), a guide with an active (non-G) PFS at the site of the qPCR probe from the N1 assay (dark gray), and 10 randomly selected guides with an active PFS (light gray). Target concentration of 10^7^ cp/μL is shown in Fig. 5b. **(b)** Fluorescence for Cas13 guides at varying target concentrations within the US CDC’s SARS-CoV-2 N2 RT-qPCR amplicon. The final time point is shown (124 minutes). Guides are as in (a). **(c)** Fluorescence over time for Cas13 guides at varying target concentrations (top of each plot, in cp/μL) within the US CDC’s SARS-CoV-2 N2 RT-qPCR amplicon. Compared guides are as in (a).

**Supplementary Figure 31.**
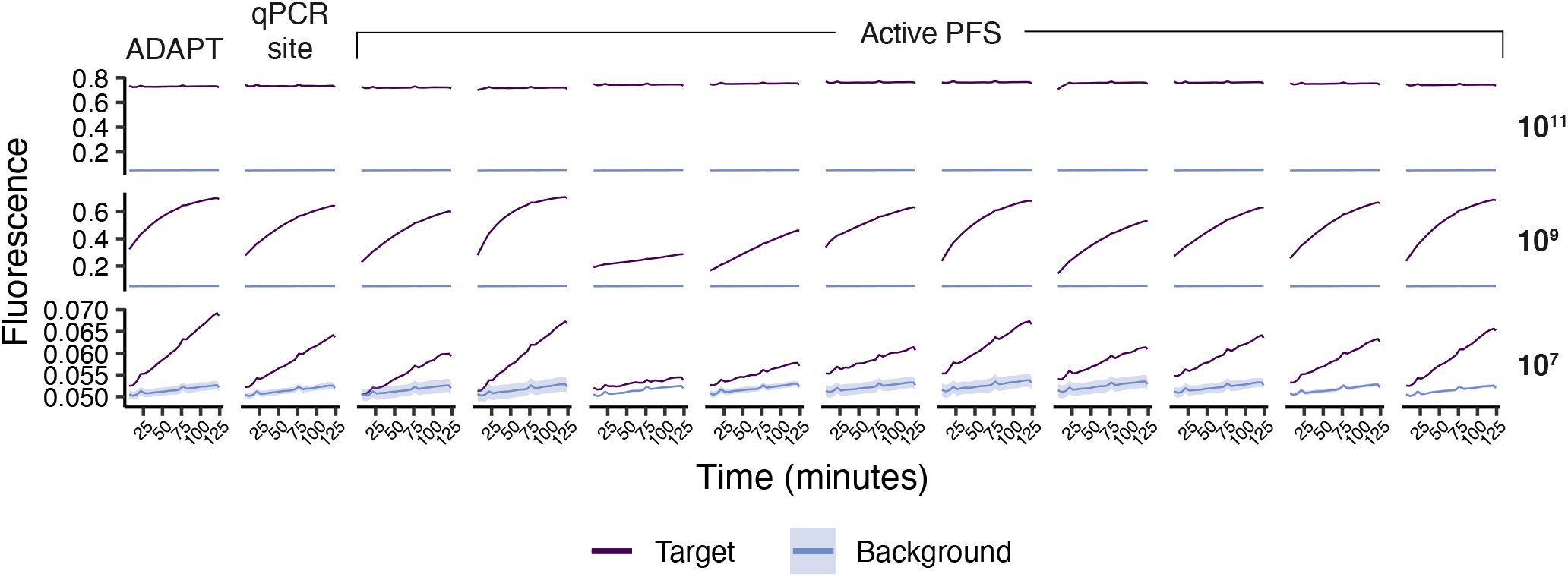
No-template background control fluorescence. Fluorescence over time against the no-template background control (water; blue) and against the template (purple) for each guide tested in the US CDC’s SARS-CoV-2 N1 RT-qPCR amplicon. Guides, separated by columns, are: ADAPT’s design, a guide with an active (non-G) PFS at the site of the qPCR probe, and 10 randomly selected guides with an active PFS. Labels on the right indicate target concentration in cp/μL (irrelevant for the background values). Shaded regions around the background values are 95% pointwise confidence bands across *n* = 7 replicates. Unlike in other plots of fluorescence, here the plotted values are not background-subtracted.

**Supplementary Figure 32.**
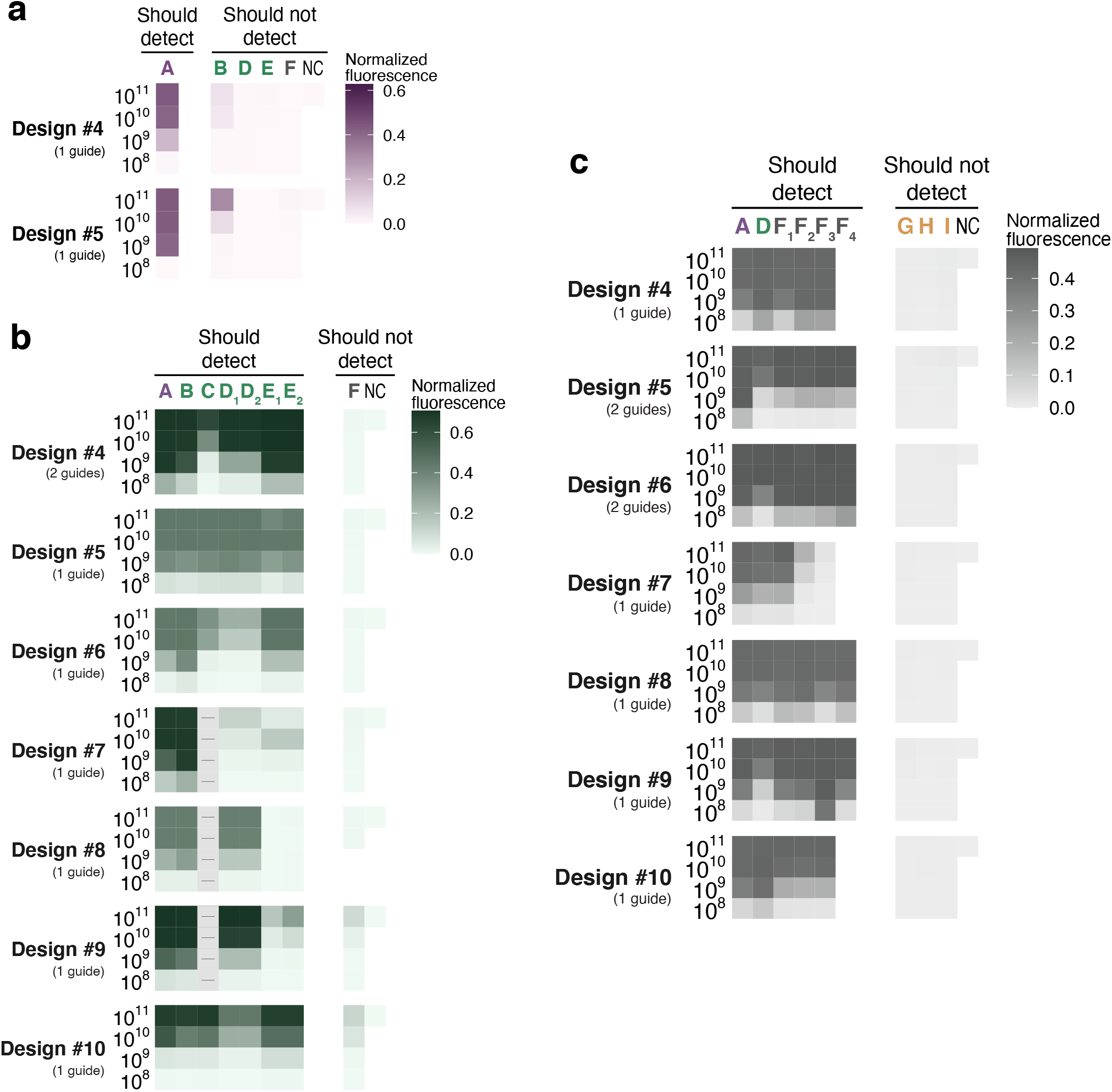
Sensitivity and specificity of additional designs for SARS-related taxa. Fluorescence for ADAPT’s designs specific to **(a)** SARS-CoV-2, **(b)** SARS-CoV-2–related, and **(c)** SARS-related coronavirus species. Fig. 5c shows phylogenetic relationships of these taxa. Assays ranked from 4 through 10 are shown (only 5 tested for SARS-CoV-2); the top 3 are shown in Fig. 5. Target definitions are in Fig. 5c and the Fig. 5 legend provides additional details about each panel. In (c), clade F required a fourth representative target (F_4_) in only some amplicons. In all panels, parenthetical numbers are the number of Cas13 guides in ADAPT’s design.

**Supplementary Figure 33.**
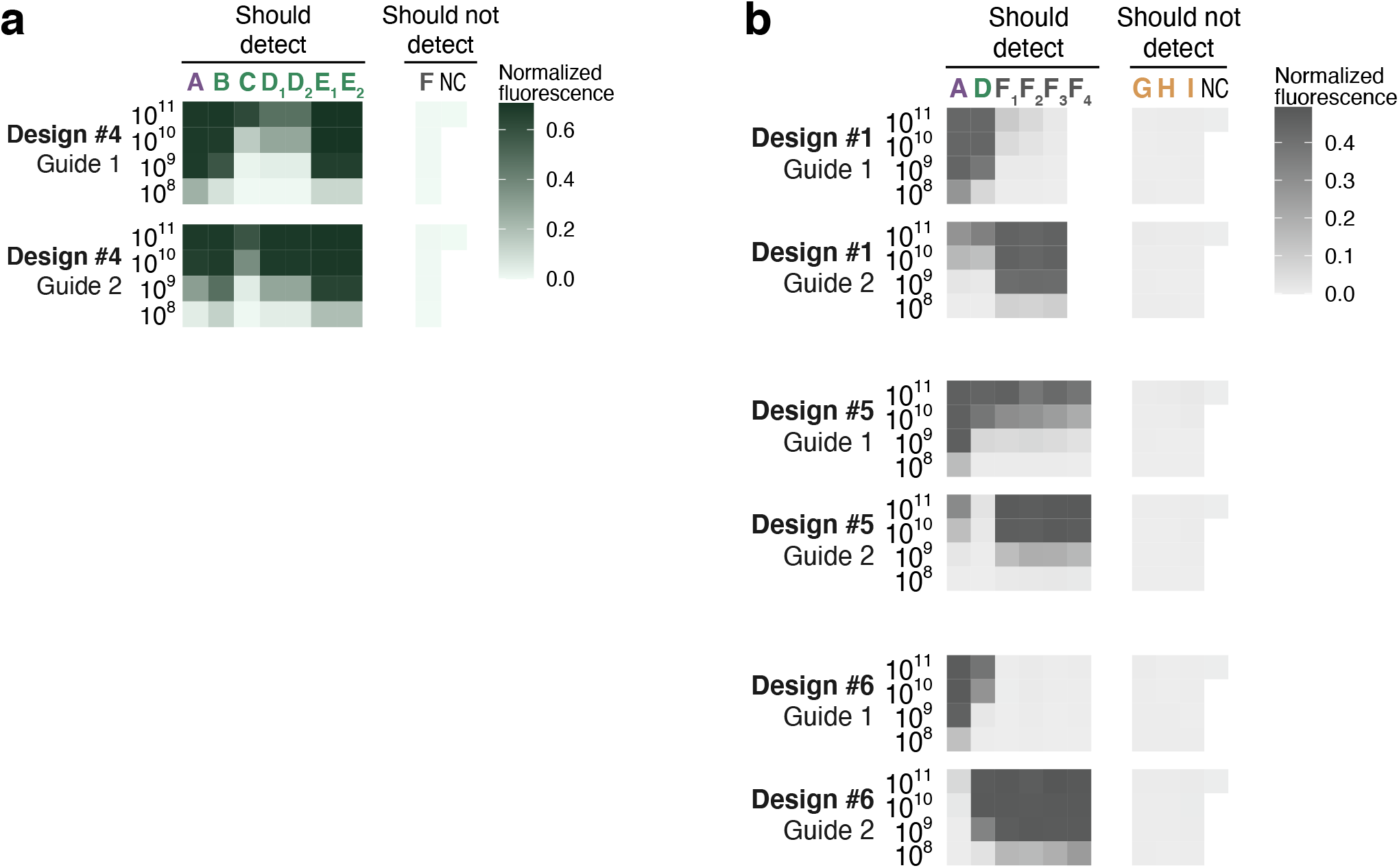
Separate guides for multi-guide designs in SARS-related taxa. Fluorescence for ADAPT’s design options that use more than one Cas13 guide, separated by guide. Left label indicates target concentration (cp/μL). **(a)** The two guides in Design #4 to detect SARS-CoV-2–related taxon (Supplementary Fig. 32b). **(b)** The two guides in Design #1 (Fig. 5e) and in Designs #5 and #6 (Supplementary Fig. 32c) to detect the SARS-related coronavirus species. In other figures, plotted value is the maximum across multiple guides.

**Supplementary Figure 34.**
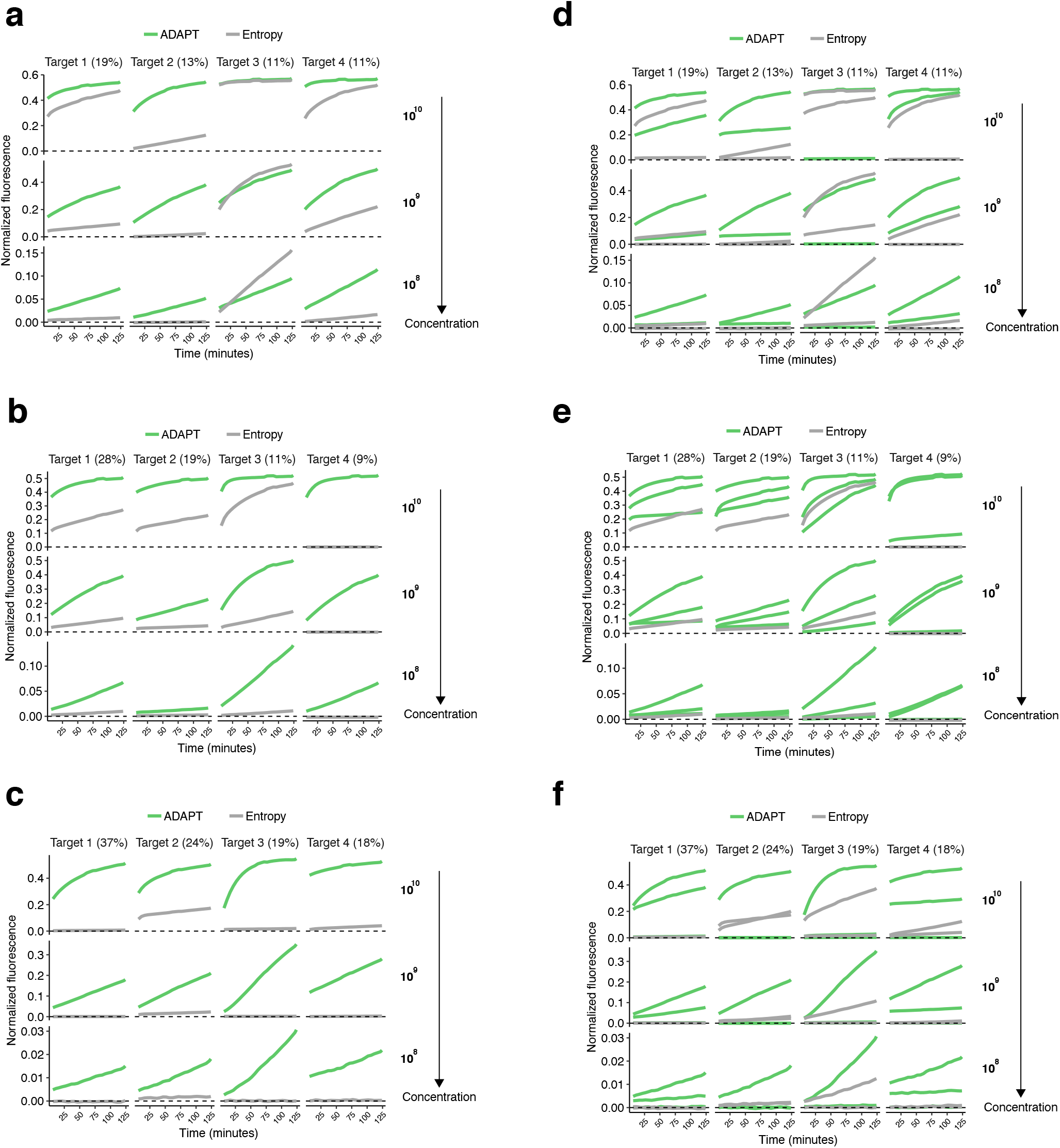
Kinetic curves of designs for detecting Enterovirus B. **(a–c)** Fluorescence over time for ADAPT’s designs in detecting EVB at varying target concentrations (right of each plot in cp/μL), for the 4 targets representing the largest fraction of EVB genomic diversity within the corresponding amplicon. (a) Design #1 (highest ranked output design by predicted performance); (b) Design #2; (c) Design #3. Plots at the target concentration of 10^8^ cp/μL are also shown in Fig. 5h. The Entropy guide (gray) targets the site from ADAPT’s amplicon with an active PFS and minimal Shannon entropy. **(d–f)** Same as (a–c) except with a separate line for each guide, when there are multiple guides in ADAPT’s design or 2 guides tested for the entropy-based approach. (d) Design #1; (e) Design #2; (f) Design #3. When there are two Entropy guides, they have an active PFS and the least and second-least Shannon entropy in the amplicon of ADAPT’s design.

**Supplementary Figure 35.**
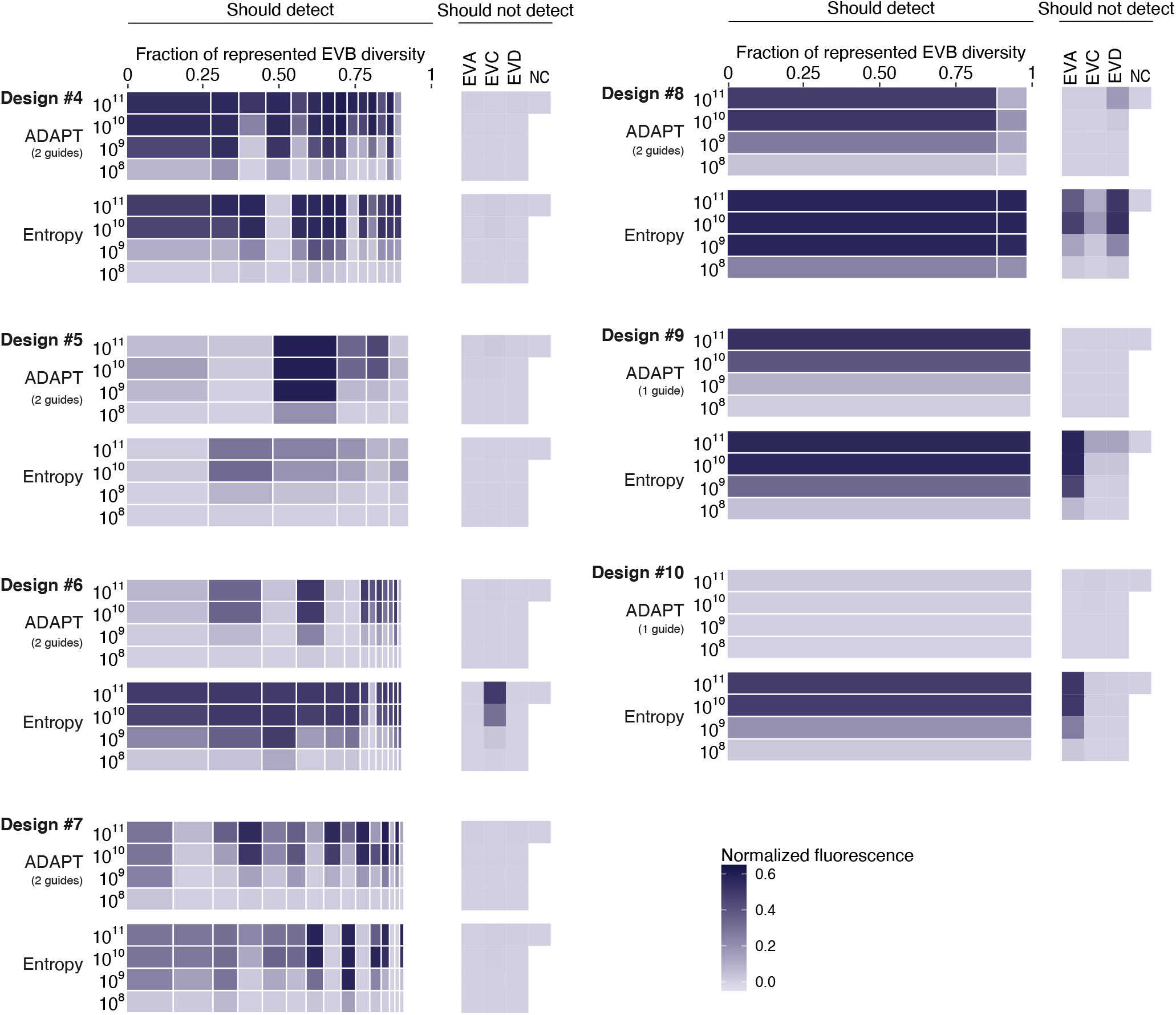
Sensitivity and specificity of additional designs for Enterovirus B. Fluorescence for ADAPT’s Enterovirus B (EVB) designs in detecting EVB and representative targets for Enterovirus A/C/D (EVA/C/D). Assays ranked from 4 through 10 are shown; the top 3 are shown in Fig. 5g. Each band is an EVB target having width proportional to the fraction of EVB genomic diversity represented by the target, within the amplicon of ADAPT’s design. Immediately under each ADAPT design is one baseline guide (“Entropy”) from the site in the amplicon with an active PFS and minimal Shannon entropy. Values immediately to the left of the bands indicate target concentration (cp/μL), and parenthetical numbers are the number of Cas13 guides in ADAPT’s design.

**Supplementary Figure 36.**
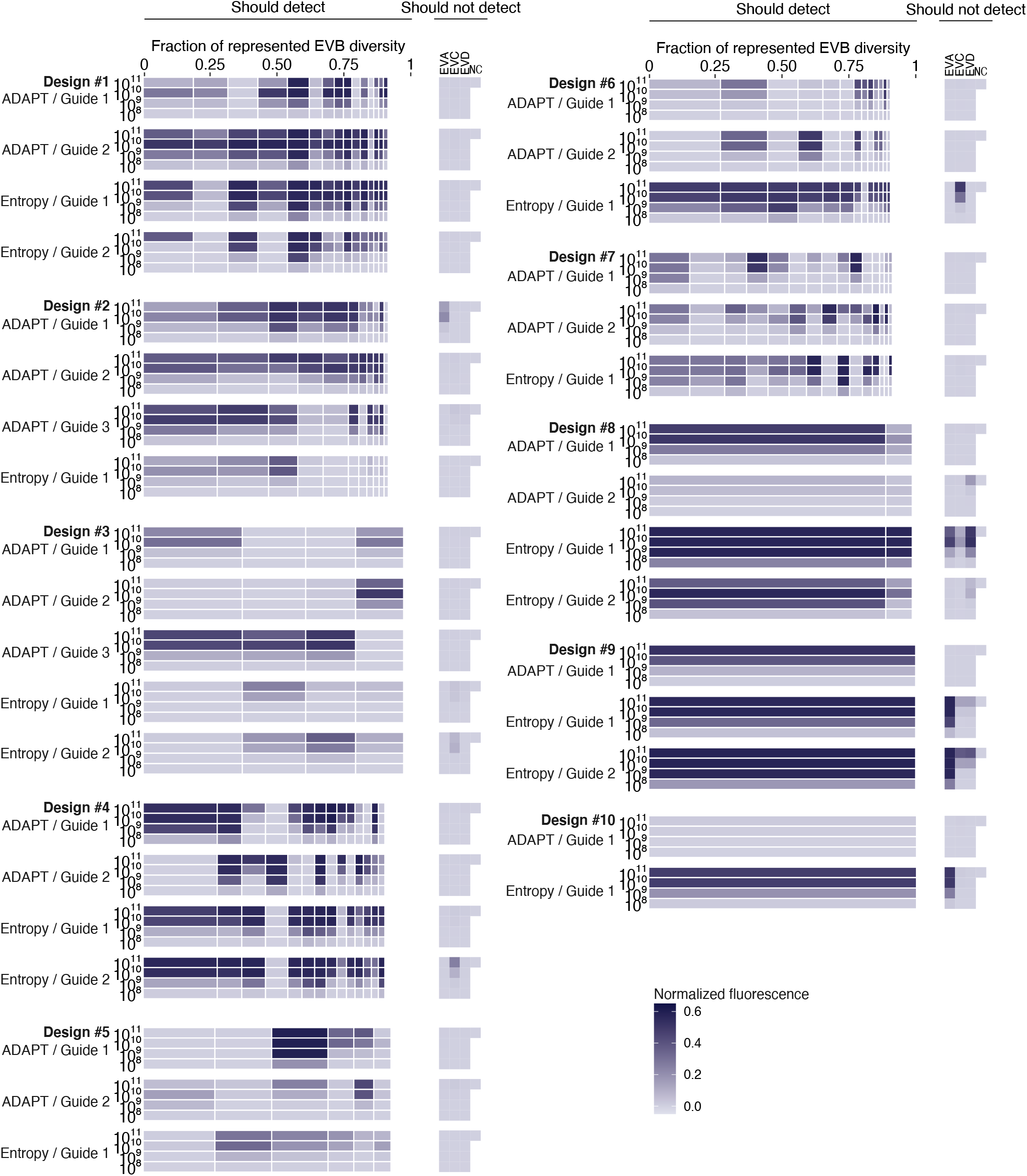
Separate guides in designs for detecting Enterovirus B. Fluorescence for ADAPT’s Enterovirus B (EVB) designs in detecting EVB and representative targets for Enterovirus A/C/D (EVA/C/D), separated by guide. Each band is an EVB target having width proportional to the fraction of EVB genomic diversity represented by the target, within the amplicon of ADAPT’s design. Values immediately to the left of the bands indicate target concentration (cp/μL). Immediately under each ADAPT design is one baseline guide (“Entropy”) from the site in the amplicon with an active PFS and minimal Shannon entropy; when there are two Entropy guides, the second is from the site with an active PFS and the second-least entropy. In Fig. 5g and Supplementary Fig. 35, plotted value for ADAPT is the maximum across multiple guides.

## Footnotes

1 Although high, this is only an upper bound and is meant to restrict the search space and thus restrict runtime.

2 In particular, by sampling nucleotides—i.e., concatenating locality-sensitive hash functions drawn from a Hamming distance family—and taking the consensus of sequences within each cluster, where clusters are defined by their hash values.

3 We synthesize the reverse complement of *g* and use that for detection, so these rules correspond to permitting G-T base pairing.

4 We typically add a fixed constant (4) to the objective values before reporting them to the user, which we find makes the values more interpretable to users because it makes them more likely to be non-negative. This shift has no impact on the design options or their rankings.

5 One technicality: many species have segmented genomes. For these, ADAPT also needs the label of the segment. ADAPT effectively treats each segment as a separate taxonomy—i.e., for species that are segmented, the *t_i_*s are actually pairs of taxonomy ID and segment.

6 The “reference” sequences are determined by NCBI, but can also be provided by the user. They are manually curated, high-quality genomes and encompass major strains.

